# Evolutionary dynamics in the decades preceding acute myeloid leukaemia

**DOI:** 10.1101/2024.07.05.602251

**Authors:** Caroline J. Watson, Gladys Y. P. Poon, Hamish A.J. MacGregor, Adriana V.A. Fonseca, Sophia Apostolidou, Aleksandra Gentry-Maharaj, Usha Menon, Jamie R. Blundell

## Abstract

Somatic evolution in ageing tissues underlies many cancers. However, our quantitive understanding of the rules governing this pre-cancerous evolution remains incomplete. Here we exploit a unique collection of serial blood samples collected annually up to 15 years prior to diagnosis of acute myeloid leukaemia (AML) to provide a quantitative description of pre-cancerous evolutionary dynamics. Using deep duplex sequencing and evolutionary theory, we quantify the acquisition ages and fitness effects of the key driver events in AML development. The first driver mutations are typically acquired in the first few decades of life when the blood remains highly polyclonal. These early slow-growing clones subsequently acquire multiple further driver mutations which confer selective advantages up to 100-fold larger than the early drivers. These faster-growing clones harbouring multiple driver mutations can cause complete somatic sweeps of the blood decades before diagnosis, a feature strongly associated with future AML. Once established in the blood, the dynamics of driver mutations are highly predictable. Trajectories are shaped by strong clonal competition between lineages with limited evidence of other extrinsic factors playing a major role. Our data show that the clonal dynamics of blood are consistent with a set of remarkably simple evolutionary rules which strike a balance between chance and determinism.

## Introduction

Cancers develop via a process of Darwinian evolution within tissues ^*1*–*4*^. Mutation, drift, and selection in the large cell populations making up our tissues can result in the emergence of a clone that has acquired multiple genetic and epigenetic “driver” alterations that disrupt tissue homeostasis and promote uncontrolled proliferation ^*5*–*8*^. Whilst large-scale sequencing efforts over the last 20 years have provided comprehensive catalogues of the driver genes responsible for cancer development ^*9*–*11*^, a complete quantitative understanding of the evolutionary dynamics governing how these drivers arise, expand and compete, is lacking. Why are cancer driver mutations widespread across many healthy human tissues ^*12*–*16*^, yet cancers of many of these tissues are comparatively rare? Why in some people do initial clonal expansions progress to lethal cancers while in others they remain benign? What are the relative roles of chance and determinism in the evolution of cancer? Answers to these questions could guide the development of more rational approaches to early cancer detection ^*17*^, risk prediction ^*18*–*22*^ and personalised intervention ^*23,24*^.

Theories of oncogenesis based on somatic evolution have existed since the 1950s and were used to explain the non-linear increases in cancer incidence with age ^*25*^. These multi-stage models highlighted important roles for mutation ^*25,26*^, selection ^*27,28*^, and stem-cell population sizes ^*29*^. However, these models have proven difficult to falsify with age-incidence data alone. The dynamics of molecular evolution, read-out using deep sequencing of serial samples, provides a more direct measure of the evolutionary dynamics and has transformed our understanding of microbial evolution ^*30*–*34*^. Applying similar approaches to the study of cancer development, however, has proven challenging as serial samples of pre-cancerous tissue from asymptomatic individuals are difficult to obtain. Even when these tissues are available they typically provide only a static snapshot of the somatic diversity present, missing key aspects the evolution that went before and what came after. To better understand the dynamics of pre-cancerous evolution requires serial samples collected with high temporal resolution.

The haematopoietic system provides a unique opportunity to understand and quantify pre-cancerous evolution because serial blood samples in asymptomatic individuals can be collected antecedent to cancer development. Healthy blood displays ubiquitous and dramatic age-related changes driven by positive selection for genetic alterations that are recurrently mutated in AML (e.g. *DNMT3A, TET2* and *ASXL1*) ^*18*–*22,35,36*^. This phenomenon, termed clonal haematopoiesis (CH), increases the risk of future blood malignancies such as AML ^*18*–*22*^. Whilst some aspects of CH evolution have begun to be understood quantitatively ^*37*–*42*^, our understanding of how these age-related changes evolve into AML, is incomplete. We reasoned that serial blood samples collected regularly over decade long timescales in people who subsequently develop AML could be used to better understand the evolutionary origins of AML. By leveraging developments in error correctable sequencing ^*43,44*^ and insights from evolutionary theory ^*31,45,46*^ we sought to characterise when in life the key driving events of AML occur, and gain insights into the rules governing their establishment, growth and competition.

## Results

### Deep sequencing of longitudinal blood preceding AML

The United Kingdom Collaborative Trial of Ovarian Cancer Screening (UKCTOCS) study enrolled 202,638 cancer-free post-menopausal women, aged 50-74 years old, between 2001 and 2005, of whom 50,640 were randomised into a “multi-modal arm” involving annual blood serum samples for up to 11 years ^*47,48*^. We identified all the women who were cancer-free at enrolment but subsequently developed AML during the >12 years follow-up, of whom 50 had sufficient blood sample available from ≥2 timepoints prior to AML diagnosis. These 50 women were diagnosed with AML at an average age of 72 years (range 53-83 years) and had up to 10 serial blood samples (median 5) available prior to diagnosis (Fig. 1a, Extended Data Fig. 1). We obtained 262 pre-AML blood serum samples from these 50 women, as well as 262 timepoint-matched serial blood serum samples from 50 age-matched controls who remained cancer-free. To precisely track the evolutionary dynamics in these longitudinal samples, we designed a highly sensitive sequencing strategy called TETRIS-seq. This approach incorporates duplex error-corrected sequencing ^*43*^ of a panel that comprehensively covers CH and AML-associated genomic alterations, including gene mutations, chromosomal rearrangements and mosaic chromosomal alterations (mCAs), in combination with a custom *in silico* de-noising algorithm (Fig. 1b, methods, Supplementary Note 1). Using TETRIS-seq we reliably achieved a duplex consensus depth of >1500x, enabling robust detection of single nucleotide variants (SNVs) and indels at variant allele frequencies (VAFs) ≥0.1% (Supplementary note 1E). Using the longitudinal nature of our samples we were able to phase single nucleotide polymorphisms (SNPs) on sections of chromosomes affected by mCAs enabling detection of chromosomal gains, losses and copy-neutral loss of heterozygosity (CN-LOH) events down to 0.1% cell fraction (Supplementary note 1G). Together this approach enabled us to sensitively and precisely track the dynamics of a broad range of AML-associated genomic alterations in the decade preceding AML diagnosis.

**Fig. 1.**
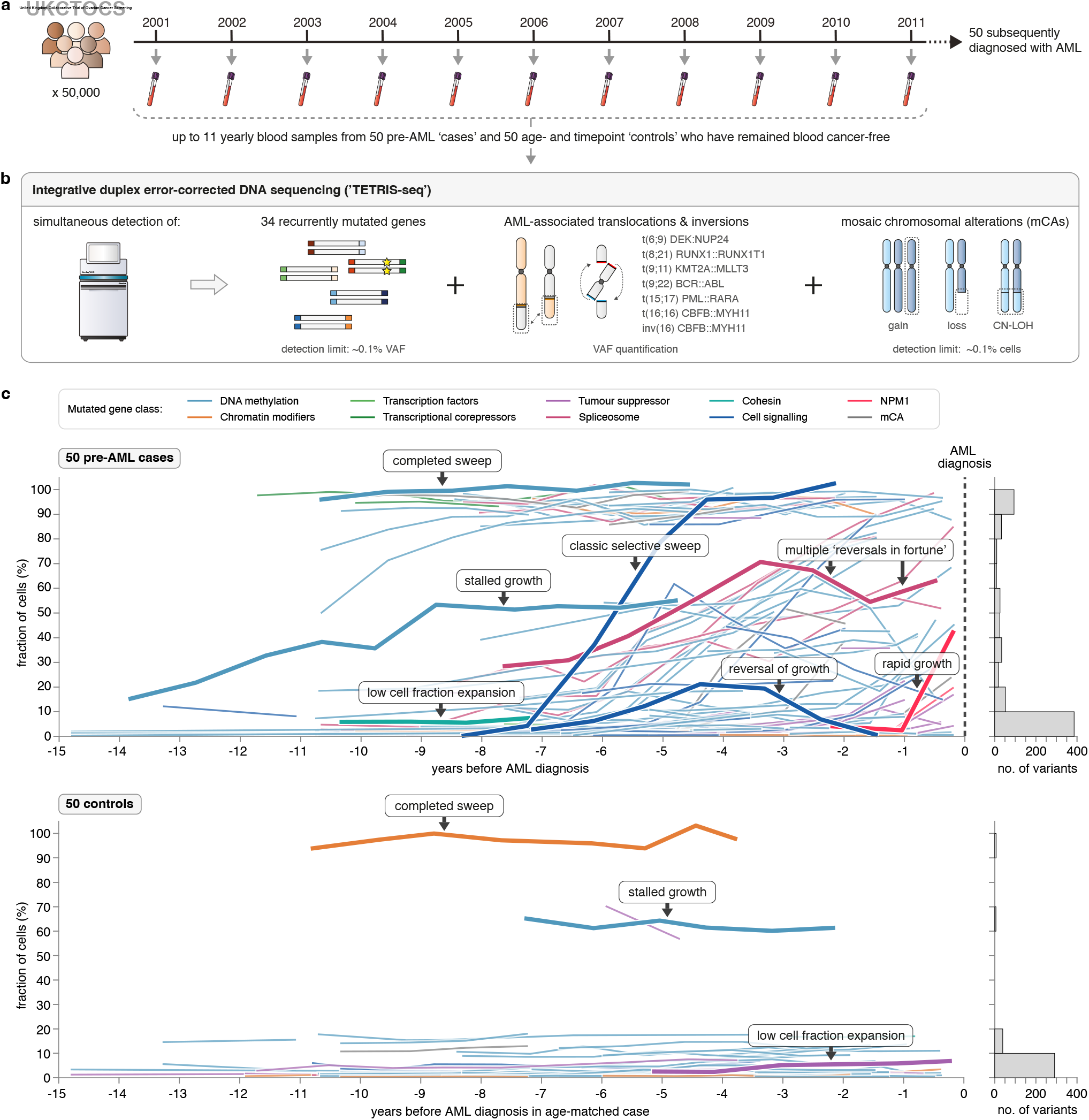
Deep sequencing of serial blood samples in the decades preceding AML. **a**. Longitudinal blood samples collected annually from 50 future AML cases and 50 age-matched controls from the UKCTOCS cohort. **b**. TETRIS-seq workflow for deep duplex targeted sequencing of AML associated genetic alterations combined with an *in-silico* de-noising algorithm. **c**. Cell fraction trajectories of all variants detected in the pre-AML cases (above) and controls (below). Characteristic dynamic behaviours are labelled and highlighted. Histograms of variant frequencies for the two groups are shown on the right-hand side.

### Mutation dynamics in the decade preceding AML

Ap-plying TETRIS-seq to these longitudinal samples revealed strikingly different somatic evolutionary dynamics between the pre-AML cases and the controls (Fig 1c). In controls, we detected CH in 78% (39/50) of people, consistent with previous studies ^*36,38*^. Variants observed in controls were typically at cell fractions <20% and trajectories appeared relatively stable over time (Fig. 1c). These variants were mostly in well-known CH driver genes, including *DNMT3A* and *TET2* (Extended Data Fig 2). In pre-AML cases, we detected CH in 90% (45/50) of people; a similar fraction to controls. In contrast to controls, however, variants observed in pre-AML cases were substantially more abundant and exhibited large cell fraction changes over the sampling period (Fig. 1c). These variants occurred in a broader range of AML-associated genes including spliceosome, cell signalling and tumour suppressor genes (Extended Data Fig 2).

The cell fraction trajectories observed in pre-AML samples exhibit a broad range of qualitative behaviours, which are distinct from the controls (highlighted trajectories Fig. 1c). We observed many examples of classic selective sweeps whereby a single variant expanded to dominate the blood. The majority of these somatic sweeps had already reached fixation prior to the start of the sampling period and were ∼19-fold *(19 vs 1)* more common in the pre-AML cases relative to controls. We also observed examples of clonal expansions whose growth stalled, reversed, or experienced multiple reversals in fortune (Fig. 1c) suggestive of frequent and strong clonal interference between parallel lineages within the same individual. These behaviours were almost exclusively observed in the pre-AML cases. Clones expanding at low cell fraction were a common feature of both pre-AML cases and controls, but were more common in pre-AML cases (histograms, Fig. 1c) and appeared to expand at faster rates pre-AML.

### Reconstruction of clonal evolutionary histories

The highly quantitative time-series data generated by TETRIS-seq enabled us to unambiguously infer the order of driver mutation acquisition and patterns of co-occurrence in many of the 50 pre-AML cases (Fig 2, see ‘Inferring clonal architecture’ in Methods and Supplementary Table 6). In 54% (27/50) of pre-AML cases, we observed ‘linear evolution’, characterised by the sequential acquisition and expansion of driver mutations along a single dominant ‘trunk’, with minimal evidence of side-branching (Fig. 2a-d, Supplementary Fig. 16). Within this class, however, there was substantial variation in the clonal composition of the blood in the decade prior to diagnosis. In 26% (13/50) of cases, we observed clear evidence of clonal interference, characterised by the acquisition and expansion of multiple driver clones on parallel evolutionary branches, which grew to high cell fractions (>20%) and interfered with each others growth, causing complex evolutionary trajectories (Fig. 2e-h, Supplementary Fig. 17). There was substantial variation in the dynamics observed within this class too, with examples of competition between quadruple-mutant clones (Fig. 2e), triple-mutant clones (Fig. 2h) and double mutant clones (Fig. 2f, g). In both the linear and clonal interference patterns of evolution the majority of the blood already derived from clones that were harbouring at least one pathogenic mutation in the decade pre-AML diagnosis. The remaining 20% (10/50) of cases are characterised by ‘late evolution’, where the pre-leukaemic clone was only detectable 1-2 years pre-AML, if at all (Figure 2c, Supplementary Fig. 17). This suggests that in some cases AML may evolve via the acquisition of uncommon driver mutations not covered by our targeted sequencing panel, that the pre-leukaemic evolution may play out whilst clones are below our detection limit, or that the acquisition of the key drivers in these cases occurs without a long pre-leukaemic phase.

**Fig. 2.**
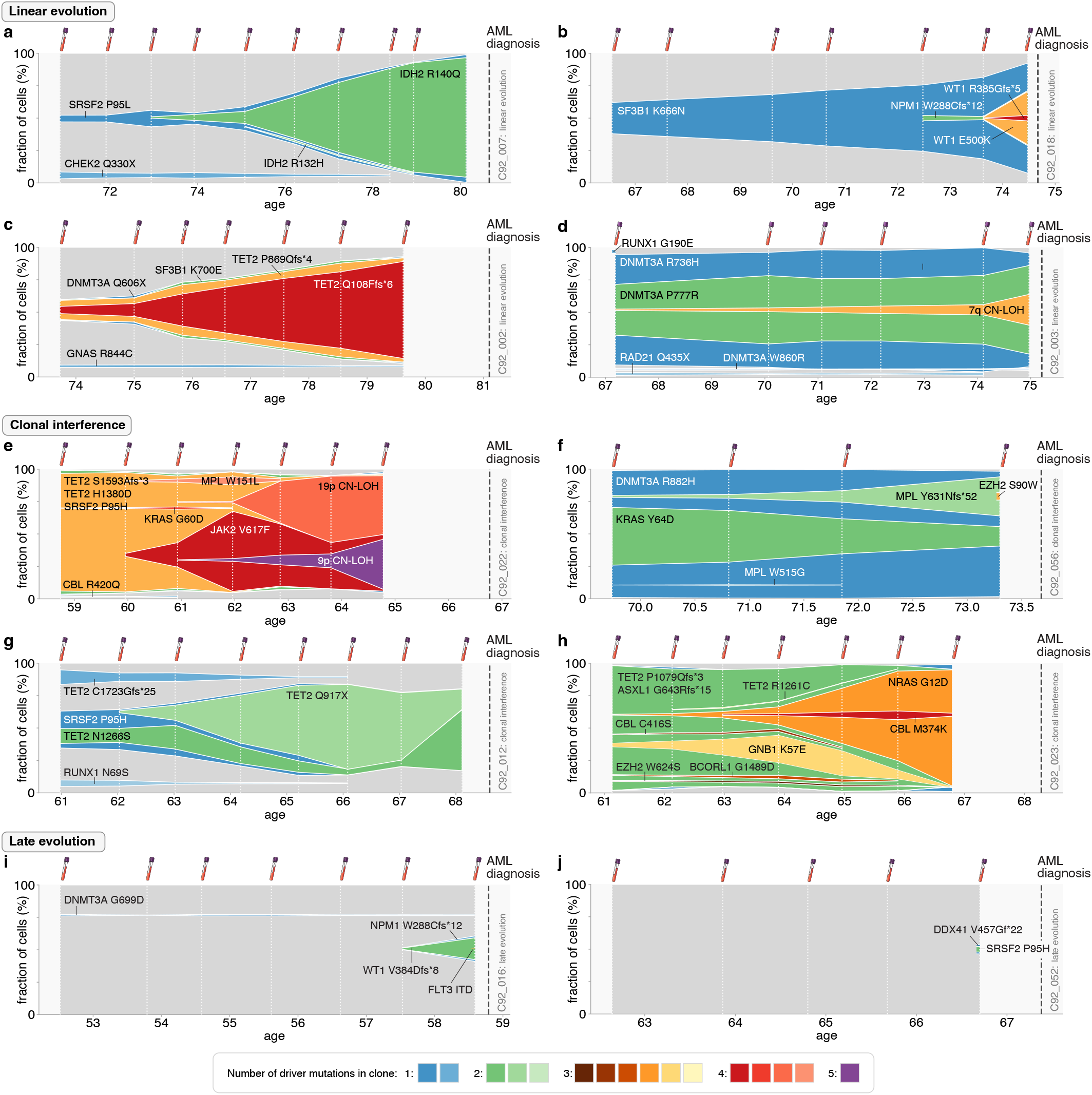
Reconstruction of clonal evolutionary histories in AML. Muller plots exhibiting clonal dynamics in 10 pre-AML cases showing “linear evolution”, “clonal interference” or “late evolution”. Colouring denotes the number of observed driver mutations in each clone (see legend). White vertical dashed lines indicate timing of UKCTOCS blood samples. Thick grey vertical dashed line indicates time of AML diagnosis. **a-d**. 4 pre-AML cases exhibiting a “linear” mode of evolution. **e-h**. 4 pre-AML cases exhibiting a “clonal interference” mode of evolution. **i-j**. 2 pre-AML cases exhibiting “late evolution”.

### Quantitative dynamics of driver mutations

We reasoned that the high sensitivity and temporal resolution of our data could be used to gain a quantitative understanding of how driver mutations expand and compete with each other in the decades preceding AML diagnosis (Fig. 3). We developed a mathematical framework for predicting how clones expand and compete through time inspired by models of microbial evolution ^*31,45*^. In this framework the dynamics of a given clone are determined by its occurrence time, fitness effect and the simplest interaction term: an increasing “mean fitness” of all other clones in the population that ensures the population frequencies of all clones sum to one. The dynamics of a specific variant are determined by summing the frequency of the original parental clone (i.e. the first clone to acquire the variant) and all daughter clones which inherited the variant (Fig. 3a-c).

**Fig. 3.**
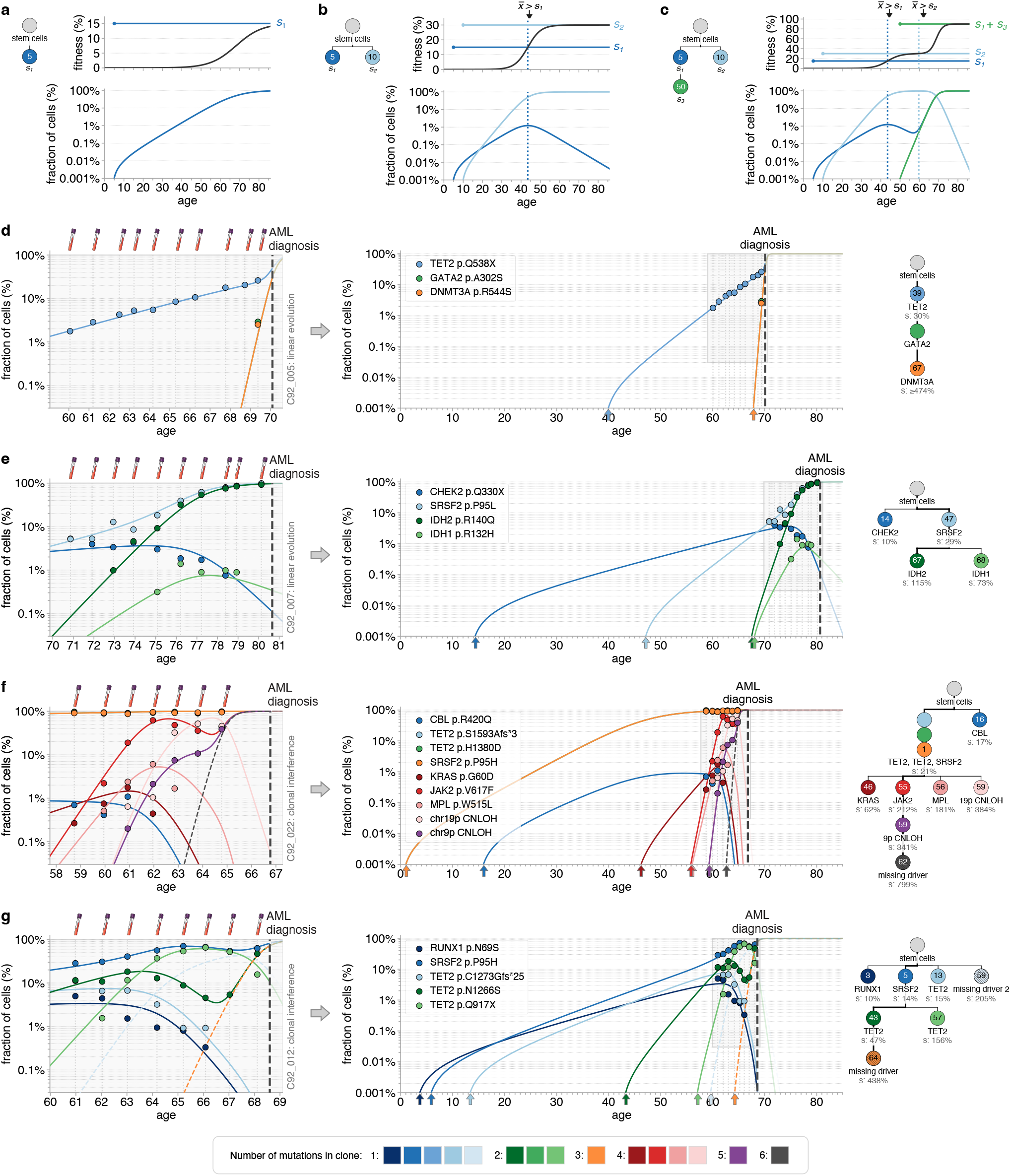
Quantitative dynamics of driver mutations in the decades before AML diagnosis. **a-c**. Schematics demonstrating how the clonal composition of the population (left-hand-side: phylogenetic trees) and estimates of the fitness and occurrence times of each clone (top plot) interact via the mean population fitness (black line, top plot) to produce different behaviours in the observed variant frequency trajectories (bottom plot). **d-h**. Observed variant frequency trajectories (data points) compared with predicted variant frequency trajectories (coloured lines) due to the most likely fitness and occurrence time estimates of clones, across the period of UKCTOCS blood sampling (left-hand-side plots) and since the birth of the individual (right-hand-side plots). Grey vertical dashed lines indicate timing of blood samples. Thick black vertical dashed line indicates time of AML diagnosis. Phylogenetic trees (right-hand-side) show the inferred clonal composition of the population, occurrence times of each clone (number in circles, years) and their fitness effect (% per year). The number of driver mutations in the clone is indicated by its colour (see legend). **d, e**. Examples of two pre-AML cases that demonstrated linear evolution. **f, g**. Examples of two pre-AML cases that demonstrated clonal interference and where we infer the presence of missing drivers (dashed trajectory lines).

After surviving an initial stochastic drift phase, a single clone carrying a driver mutation expands exponentially at a rate determined by its fitness. This clonal expansion causes the population mean fitness 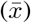 to increase, eventually caus-ing the clonal expansion to slow down as it sweeps to fixation (Fig. 3a). Competition between two independent clones with different fitnesses can lead to the variant frequency trajectory of the less fit clone decreasing once its fitness is overtaken by the population mean fitness (Fig. 3b). Competition between clones within a branch and between branches can drive more complex variant frequency dynamics involving multiple reversals of fortune due to clones jumping ahead of, or falling behind, the population mean fitness (Fig. 3c). Using this framework we developed a maximum likelihood approach (methods) to infer fitness and occurrence time estimates for each clone in the population and compare the variant frequency trajectories predicted by the maximum likelihood inference (lines) to the measured variant frequency trajectories (data points) in pre-AML cases (Fig. 3d-g, Extended Data Figs. 3-9, Supplementary Fig. 22) and controls (Supplementary Figs. 23-30).

We observe a remarkably close agreement between the variant frequency trajectories predicted by this simple framework and the observed data. We first consider a case of linear evolution with no evidence of clonal competition (C92_005, Fig. 3d). Here we infer that a *TET2* variant was acquired at ∼ 39 years of age and expanded exponentially with a doubling time of ∼2.4 years (fitness of 30% per year). Approximately 30 years later (at age ∼ 67) this lineage acquires 2 further driver mutations which cause rapid clonal expansion with a doubling time of ∼8 weeks (fitness of ≥474% per year). Diagnosis occurs three years later at 70 years of age. Case C92_007 also exhibits a predominantly linear pattern of evolution (Fig. 3e). Here we infer that the first pre-leukaemic clone (*SRSF2* p.P95L) is acquired at ∼47 years of age and expands with a doubling time of ∼2.5 years (fitness of 29% per year). Approximately 20 years later, (at age ∼67), this lineage acquires an *IDH2* p.R140Q mutation which creates a double-mutant clone that expands rapidly with a doubling time of ∼7 months (fitness effect of 115% per year). This double-mutant clone sweeps the blood when the person is in their late 70s and they are diagnosed with AML at age 80, 13 years after acquisition of the *IDH2* mutation. The sweep of the double mutant clone harbouring *SRSF2* and *IDH2* variants causes two low-frequency variants (in *CHEK2* and *IDH1*) to be outcompeted, highlighting that high-frequency clones provide important information about the behaviour of low-frequency clones and visa-versa.

We next considered cases where there was clear evidence of clonal interference between competing lineages (Fig. 3f, g). In C92_022 there has already been a full sweep of the blood by a clone harbouring three driver mutations (bi-alleleic loss of *TET2* and *SRSF2* p.P95H) by the time of the first sample, 8 years prior to diagnosis (Fig. 3f). On the background of this triple-mutant clone there is rapid evolution involving the acquisition of multiple further driver events. First, two independent clones caused by the acquisition of *JAK2* V617F and *MPL* W515L arise at age ∼ 55 and expand with doubling times of 3.9 and 4.6 months respectively (fitness effects of ∼212% and ∼181% per year). These are soon outcompeted by a substantially fitter clone carrying 19p CN-LOH which is acquired at ∼59 years of age and expands with a doubling time of just 2.2 months (fitness effect of ∼384% per year). Remarkably, however, this lineage does not sweep. Instead, the *JAK2* V617F lineage acquires two further events: a 9p CN-LOH and a subsequent event of unknown origin (“missing driver”) which grows with a doubling time of ∼1 month and sweeps the entire blood in under 2 years. Without invoking the presence of a highly fit missing driver (dashed trajectory, Fig. 3f) the variant frequencies measured at age 65 would be inconsistent with the behaviour of the observed clones. Thus the internal consistency between the variant frequency trajectories can be used to detect the presence of events missed by our sequencing panel (methods). C92_012 (Fig. 3g) highlights another example of clonal interference where the observed variant allele frequency trajectories suggest the presence of two missing drivers not captured by our sequencing panel. Despite these driver events being missed, the dynamics of the other observed driver events enable us to estimate their occurrence times and fitness effects.

Overall these findings demonstrate that, once established in the haematopoietic stem cell (HSC) pool, the dynamics of pre-leukaemic clones are surprisingly predictable. Whilst we cannot exclude the possibility of extrinsic effects altering some clones, one does not need to invoke more complex scenarios to explain the data: one can achieve remarkably close fits to variant trajectories by assuming clones have an intrinsic fitness and compete globally for one resource with minimal spatial structure.

### Acquisition age and fitness of driver mutations

Ap-plying our framework to all pre-AML cases enabled us to estimate the fitness and occurrence time of many of the key events driving the development of AML in these 50 cases (Fig. 4) and controls (Supplementary Figs. 23-30). In ∼80% of pre-AML cases, our data suggests that the acquisition of the key driver events in AML evolution plays out over decades. In half of individuals who develop AML, the first driver mutation has already been acquired by age 20 (Fig. 4, top panel). On average, these first driver events occur ∼50 years before AML diagnosis, but expand with relatively modest fitness advantages of ∼ 10% − 30% per year (doubling times of ∼ 3 − 10 years Fig. 5). A quarter of individuals have acquired their second driver mutation by age 50 (Fig. 4, top panel). On average, these second ‘hits’ occur ∼30 years before AML diagnosis and confer a substantially larger fitness advantage to cells, causing them to expand at rates between ∼ 20% − 100% per year (doubling times of ∼1-3 years, Fig. 5). Remarkably, this pattern continues as the pre-AML clone acquires further driver mutations. Clones with three driver mutations grow with fitness advantages of between ∼ 50% − 200% per year and quadruple mutants range from ∼ 100% − 1000% per year, doubling in a matter of weeks to months (Fig. 5). This remarkable increase in fitness with the number of driver mutations acquired is close to geometric: each driver mutation acquired confers a 2-4-fold increase in the growth rate. This finding strongly suggests that pre-leukaemic mutations combine synergistically.

**Fig. 4.**
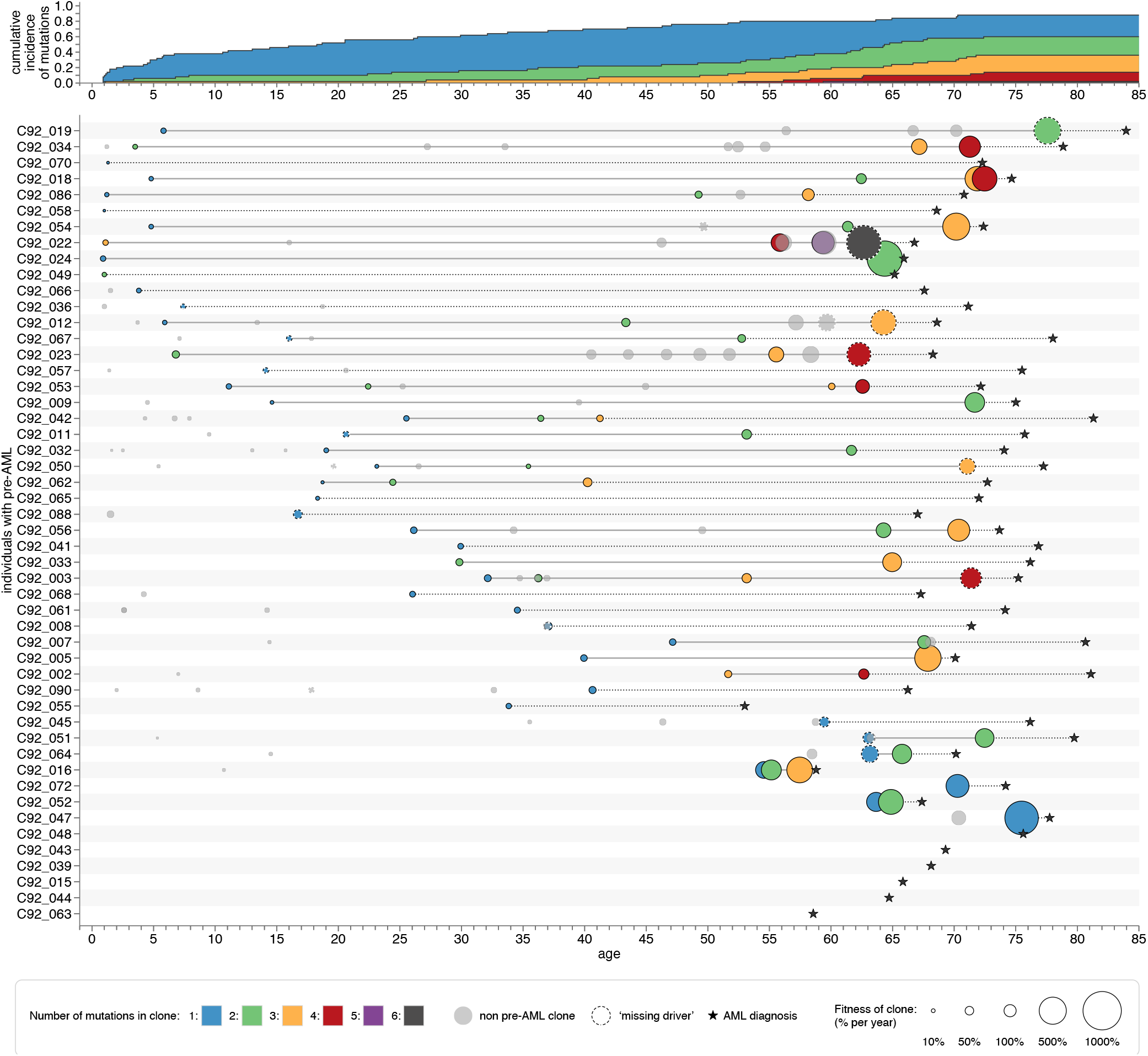
Fitness and occurrence time estimates of pre-leukaemic driver events. Occurrence time (x-axis) and fitness effect (data point size) estimates for clones arising in the 50 pre-AML cases. AML diagnosis is denoted by a black star. Solid lines connect the first driver event ancestral to the AML to the last detected driver event ancestral to the AML (an estimate for the detectable pre-leukaemic window). Greyed data points are clones on side branches (not ancestral to the AML). Dashed outlines of data points denotes a missing driver. Colour denotes the number of somatic driver mutations detected in the clone. Top panel: cumulative distributions of the occurrence time estimates for clones broken down by number of driver mutations in the clone (denoted by colour).

**Fig. 5.**
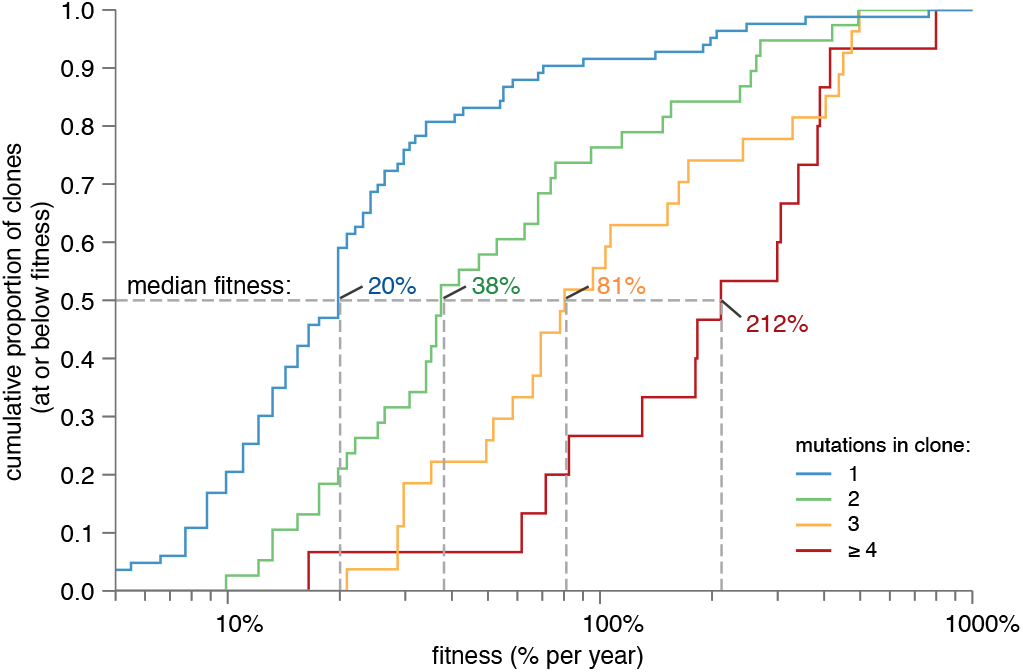
Fitness effects of driver mutations in pre-AML cases. Cumulative distributions of fitness effects for driver mutations occurring in clones harbouring one (blue), two (green), three (orange) and greater than three (red) driver mutations. The x-axis is on a log scale demonstrating the geometric progression of fitness with driver mutation number.

### A model of evolutionary dynamics in blood

The quantitative picture of the evolutionary dynamics afforded by our data highlight several important features. First, transformation usually involves the acquisition of multiple driver mutations in the same clone (range 2–6), with considerable inter-patient variation in when these driver events are acquired. Second, pre-leukaemic clones are subject to strong levels of positive selection with fitness advantages that vary by two orders of magnitude: ranging from 10% per year for the least fit single-mutant clones to 1000% per year for the fittest quadruple-mutant clones. Third, the variant frequency trajectories exhibit diverse qualitative behaviours suggesting strong clonal competition in some cases (e.g. C92_012 and C92_022) but a complete absence of competition in other cases (e.g. C92_005 and C92_007).

We hypothesised that many of these features may be emergent properties of a stochastic evolutionary model governed by a simple set of rules. To test this, we developed a model of the clonal dynamics inspired by previous models of microbial evolution ^*31,45,46*^, which we modified to incorporate key experimental observations (methods). We used this model to simulate the evolutionary dynamics in a”virtual” cohort of tens of thousands of simulated individuals for 85 years (Fig. 6). In this model, blood is maintained by a population of *N* cycling HSCs that divide symmetrically every *τ* years (Fig. 6a). HSCs acquire driver mutations at a rate *U* per year. Driver mutations vary in the fitness effects they confer to HSCs. A driver mutation arising in a wildtype HSC, creating a single mutant clone, confers fitness advantages drawn from a distribution with a median and range controlled by a single parameter *s* ∼ 10% per year (methods). A driver mutation arising in single-mutant clone confers a median fitness effect that is *r* = 3-fold larger (*s* ∼ 30% per year) creating a double-mutant clone. Triple-mutant and quadruple-mutant clones follow the same geometric pattern (with median fitness effects of *s* ∼ 100% and *s* ∼ 300% per year respectively). In the framework presented here, transformation to frank AML is defined by the acquisition of 4 driver mutations within the same clone that reaches *>* 50% cell fraction before the age of 85. Using established estimates for *Nτ* = 10^5 *37,38,40*^, *U* ∼ 10^−5^ per year ^*39*^ and *s* ∼ 10% per year for single-mutant fitness effects ^*38,41,50*^, we simulated the clonal dynamics in 25, 000 simulations and considered the evolutionary dynamics that is observed in the decades preceding the ∼ 100 virtual incidental AMLs (Fig. 6b-d).

**Fig. 6.**
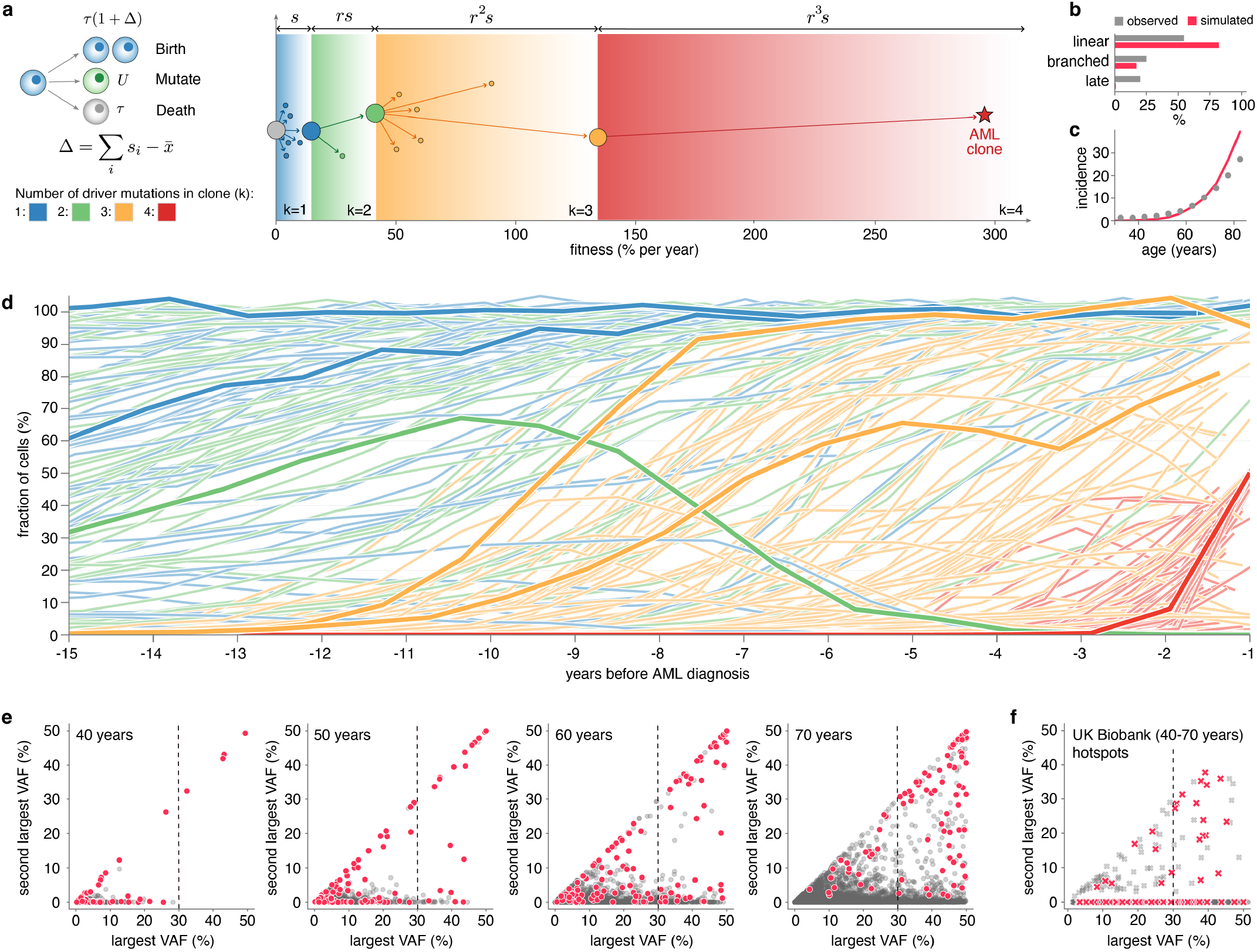
A unifying framework for pre-leukaemic clonal dynamics. **a**. A stochastic branching model of evolutionary dynamics (right-hand side) and schematic (left-hand side) of how clones (circles) give rise to daughter clones via the via the acquisition of driver mutations. Driver mutations confer increasingly large “jumps” in fitness (x-axis) from wild type HSCs (grey), to single- (blue), double- (green), triple- (orange), and quadruple-mutants clones (red). Simulated AMLs are defined by clones that acquire 4 driver mutations (red star). Growth rates of clones are determined by relative fitness (Δ) between the clone and the mean fitness of all other clones in the population 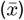. **b**. Fraction of virtual AMLs assigned to linear, branched or late evolutionary patterns for simulated AMLs (red) and observed cases (grey) **c**. Age-incidence (per 100,000 per year) of virtual AMLs (red) and clinically detected AMLs (grey, data from CRUK ^*49*^. **d**. Variant allele trajectories in 142 incidental simulated AMLs (from 25,000 simulations) coloured by the number of driver mutations in the clone. **e**. Joint frequency plots showing the VAFs of the largest and second largest variants detected in 25,000 simulated individuals. Future AML cases (red) and controls (grey). **f**. Equivalent plot to (**e**) for any variants at one of 20 CH hotspot positions (methods) detected in the UK Biobank. Future AML cases (red) and controls (grey).

Despite the simple nature of this model, it reproduces many of the qualitative and quantitative features observed in our data. Both linear and branched patterns of evolution are expected to emerge under this model with the linear pattern predicted to be more common (Fig. 6b). The model predicts a strongly increasing incidence of AML with age, in close quantitative agreement with the observed real-world incidence of AML (Fig. 6c). The variant frequency trajectories in the decades preceding virtual AMLs in this model also exhibit a similar spectrum of qualitative behaviours to the observed pre-AML cases including examples of selective sweeps, reversals of growth, multiple reversals of fortunes and late rapid expansions (Fig. 6d). Despite the simulated controls being governed by an identical evolutionary process, these behaviours are almost entirely absent from controls (Extended data Fig. 10a–d), demonstrating the profound effect that conditioning on a rare outcome can have on the observed dynamics.

We next explored whether this model could provide insights into features of the evolutionary dynamics that identify future AML cases. Simulated variant frequency trajectories plotted over a 50-year period revealed that most future AMLs in this model derive from instances where a single-mutant or double-mutant variant has already completely swept the blood more than a decade prior to virtual AML (Extended Data Fig. 10e). To check how predictive these somatic sweeps are of future AML we considered all simulated individuals that acquired at least one driver variant that could have been filtered out as “germline” in low depth sequencing data (VAF>30%) broken down by age (Fig. 6e). In our evolutionary model ∼95% of virtual AML cases have had at least one full somatic selective sweep >5 years prior to diagnosis and ∼75% have a had at least one sweep >10 years before diagnosis. The presence of a second somatic driver (VAF>10%) in addition to the first sweep identifies a group at high risk of future AML with PPV ∼100% (12/12) below the age of 50, ∼80% (46/61) at age 60, and ∼50% (86/177) at age 70.

The presence of somatic sweeps could be used to identify groups at high risk of future AML that have previously been missed due to the inability to distinguish somatic sweeps from germline variants ^*20,21*^. To demonstrate this we considered individuals in the UK biobank whose baseline blood sample contained somatic variants at VAFs high enough to be hard to distinguish from germline variants in low depth WES or WGS data (VAF>30%, Fig. 6f). Considering variants at well known somatic hotspot positions (methods), where the likelihood of germline variation is substantially lower, we find that individuals with at least one somatic sweep are at considerably increased risk of a future AML relative to those who do not (OR = 98.8). This finding suggests filters imposed for removing germline variants from somatic variant calls may in fact be removing complete somatic sweeps which are strongly associated with future AML risk.

## Discussion

These data have produced a quantitative picture of how AML evolves from precursor clones in the decades preceding diagnosis. Estimates for the acquisition ages and fitness effects for the key driving events in AML evolution highlight that in most cases the first steps of the multi-step path to AML occur early in life. In these early stages of pre-leukaemic evolution the blood is usually polyclonal with multiple single-mutant clones expanding in parallel from low cell fraction, a finding consistent with what is observed in CH ^*40*^. The fitness of these early pre-leukaemic clones is similar to those observed in CH ^*38,41,42*^, and similar to the fitness effects measured in our set of age-matched healthy controls. The dynamics of early driver variants are still dominated by positive selection rather than drift, but they expand slowly: taking years to double in size and decades to become detectable.

The acquisition of additional drivers on the background of these initial slow-growing clones appears to be a defining difference between controls and future-AML cases. In most pre-AML cases the initial slow-growing clone produces daughter clones carrying two, and later, three or four, driver mutations that grow far more rapidly than the single-mutant clones. In controls such multiple-mutant clones are rare: a double-mutant clone expanding to high VAF is expected to occur in less than 1 in 3,000 of 60-year olds (Fig. 6e). At these later stages of pre-leukaemic evolution, the blood is often oligoclonal: multiple double- and triple-mutant clones arise on independent lineages and, because of their larger fitness effects, expand and compete with each other, producing classic patterns of clonal interference ^*30,45,51*^. These patterns of clonal evolution emerge when the population size is large enough that fitness-increasing mutations are not rare, but not so large that they are very common (*θ* = *Nτµ* ∼ 1). Given existing estimates for *Nτ* ∼ 10^5^ years in humans ^*37,38,40*^, our data indicates that, at every point in the multi-stage path to AML, the rate at which driver mutations are acquired is on the order of ∼ 10^−5^ per cell per year, consistent with 1000s of sites in the genome offering mutational paths to increasing fitness. Whilst mutations that lead to increased fitness of clones are common at every stage of the evolution, the probability that the full complement of drivers required for transformation are acquired in a given person is a small, albeit strongly increasing function of age. This conceptual framework may therefore partially reconcile the fact that cancer drivers are common in healthy tissues ^*12*–*16*^, whilst still predicting that cancer itself would be relatively rare ^*49*^. We note that this evolutionary framework may also provide a natural way for incorporating recent findings on environmental risk factors ^*52*^. Exposures that increase the fitness of mutant clones will lead to more rapid clonal expansions that would potentially promote oncogenesis while exposures that suppress the growth advantage of mutant clones would be protective.

Stochasticity plays a crucial role in these dynamics. The acquisition of highly fit double- and triple-mutant clones early in life appear to be key determinants of future AML risk. This is exemplified by early complete sweeps of the blood in which all blood cells derive from a single HSC ancestor. We observe this in at least a third of the 50 pre-AML cases – a conservative estimate as we know our panel misses some driver events. Identifying early somatic sweeps thus might be a fruitful avenue for improving early detection and risk prediction for AML. However, identifying such events in large-scale cohorts such as UK Biobank is challenging due to filtering of germline variants at high VAF. Despite the large odds ratios associated with somatic sweeps it is unlikely that somatic variants that have been misclassified as germline would have been detected in GWAS studies as they would be expected to be extremely rare. We speculate that risk prediction for future AML may be able to be substantially improved by being able to identify somatic sweeps e.g. by removing germline contaminants using a second tissue sample such as buccal swabs. The fact that the 2nd and 3rd “hits” in this multi-stage process typically occur more than a decade prior to diagnosis, provides a long window of opportunity for potential intervention. The *IDH2* R140Q double-mutant in C92_007 is a case in point. It expands exponentially for 14 years with a fitness effect of ∼115% before diagnosis (Fig. 3e). Were the growth of this clone to be reduced (e.g. using an IDH inhibitor ^*23*^ or a neoantigen vaccine ^*24*^ it is tempting to speculate whether AML could have been prevented. Due to the exponential nature of the clonal growth, even modest reductions in growth rate — compounded over decades — can have a profound effect on the size of pre-leukaemic clones and thus on the probability of them acquiring further transforming mutations. We therefore speculate that low toxicity interventions producing even modest reductions in growth rate to high-risk clones may be able to substantially reduce the risk of transformation.

A surprising finding in our data was how predictable and deterministic the dynamics of established clones are. Variant trajectories are remarkably consistent with what would be expected under the simplest model of clonal competition: i.e that clones compete only via a global interaction term requiring that their frequencies sum to one. This suggests that, by the time clones have grown to be >0.2% cell fraction in the blood, the effects of spatial structure in the bone marrow are averaged out due to mixing and re-seeding of multiple niches ^*53,54*^. Whilst we cannot exclude the possibility that changing extrinsic factors shape some of the clonal dynamics observed, it is notable how well the variant frequency trajectories can be quantitatively explained without invoking these more complex scenarios.

Our ability to measure how somatic evolution shapes, and is shaped by, disease risk is a new frontier of human genetics ^*55*^. A quantitative understanding of this somatic evolution is an important part of deciphering how and why disease risk changes as we age. The haematopoietic system is an ideal model for developing such a quantitative understanding due to its ease of sampling, its relative lack of spatial structure and the existing quantity of large data sets. Our work is a step towards a complete understanding AML development at a quantitative level. Further work is needed to understand whether and to what extent these principles translate to other cancers.

## Methods

### Longitudinal peripheral blood DNA samples

Longitudinal peripheral blood DNA samples were obtained from blood serum samples biobanked as part of the UK Collaborative Trial of Ovarian Cancer Screening (UKCTOCS) (ISRCTN22488978; ClinicalTrials.gov NTC00058032). UKCTOCS is a multi-centre randomised controlled trial which recruited 202,638 postmenopausal women aged 50-74 years, from England, Wales and Northern Ireland between 2001 and 2005, to assess the impact of screening on ovarian cancer mortality ^*47*^. UKCTOCS eligibility criteria and trial details have been described previously ^*56*^. UKCTOCS was reviewed and approved by the North West Multi-centre Research Ethics Committee (MREC) (REC reference 00/8/34) and participants provided written consent for use of their samples and data in ethically approved secondary studies. Our study, using UKCTOCS samples and data, was reviewed and approved by the South Central Berkshire B Research Ethics Committee (REC reference 18/SC/0481).

At UKCTOCS enrolment 50,640 participants were randomised to a ‘multimodal screening arm’, which involved annual blood serum samples for up to 11 years up until 31st December 2011. All samples were collected using a standardised protocol. Blood was collected in gel tubes (8mL gel separation serum tubes; Greiner Bio-One 455071) at the trial centres and transported daily at room temperature to a central labratory, where the samples were centrifuged (1500g for 10 min) and the separated serum aliquoted into 10 x 500*µ*L straws using a semi-automated MAPI platform (IMV). The straws were stored in liquid nitrogen tanks at the central laboratory which, when full, were transported to a HTA licensed commercial cryofacility (Fisher Bioservices, UK until March 2018, NIHR Biocentre April 2018-July 2023 and BioDock since then). The median time from sample collection to centrifugation (∼22.1 hours) resulted in leukocyte DNA leaking into the serum ^*57*^. UKCTOCS participants’ health outcomes were followed, via linkage to national cancer, death and administrative registries. By 2017, there were 50 women in the ‘multimodal screening arm’ that had incidentally developed AML (ICD-10 C92.0) and had sufficient serum sample available from *geq*2 timepoints prior to AML diagnosis. These 50 women were diagnosed with AML at a median age of 72 (range 53-83 years, interquartile range 8 years) and had serum samples biobanked from a median of 5 timepoints (range 2-10, interquartile range 4.75) prior to AML diagnosis (Extended Data Fig. 1a, b). We obtained 1ml serum from each of the yearly timepoints available (262 samples) from these 50 women (‘pre-AML cases’), as well as from timepoint-matched serial samples from 50 age-matched UKCTOCS participants who remained blood cancer free (‘controls’). DNA was extracted by LGC Genomics, using an adapted Kleargene™ method with mag beads on KingFisher™ 96, and eluted in 10 mM Tris-Cl pH 8.0. The DNA was quantified using PicoGreenTM, yielding an average of 273 ng/ml DNA (range 11-814 ng/ml serum) in the pre-AML samples and 269 ng/ml DNA (range 2-1313 ng/ml serum) in the control samples (Extended Data Fig. 1c).

### TETRIS-seq custom panel design

To ensure we could track the evolution of a comprehensive array of clonal haematopoiesis- and AML-associated genomic changes in the longitudinal pre-AML samples and control samples, we developed 2 separate targeted panels, to be used on the same input DNA sample, in an integrated library preparation approach: a small panel (∼ 58kb) specifically for SNVs and indels and a large panel (∼1.6 MB) for mCA, chromosomal rearrangement and KMT2A partial tandem duplication (KMT2A-PTD) detection (Supplementary Note 1). For the SNV and indel panel, we designed a custom set of 120-nt oligonucleotide probes (TWIST Biosciences) targeting commonly mutated exons within 32 genes commonly mutated in clonal haematopoiesis and AML (Supplementary Figs. 1-4, Supplementary Table 1): *ASXL1, BCOR, BCORL1, CBL, CEBPA, CHEK2, CSF3R, DDX41, DNMT3A, EZH2, FLT3, GATA2, GNAS, GNB1, IDH1, IDH2, JAK2, KIT, KRAS, MPL, NPM1, NRAS, PPM1D, PTPN11, RAD21, RUNX1, SF3B1, SRSF2, STAG2, TET2, TP53, U2AF1, WT1, ZRSR2*.

For the mCA, chromosomal rearrangement and KMT2A-PTD panel, we designed a custom set of 120-nt oligonucleotide probes (TWIST Biosciences) targeting a total of 10,326 common SNPs (minor allele frequency (MAF) > 0.01) spaced every ∼ 280 kb across the genome to allow the detection of mCAs (Supplementary Table 2). Using this approach, deviations in heterozygous SNP B-allele frequencies (BAFs) can be detected which, when combined with read depth information, enables detection of a gain, loss or CN-LOH event. For detection of chromosomal rearrangements, a custom set of 120-nt oligonucleotide baits (TWIST biosciences) was designed to target the known breakpoint regions of each of the rearranged partner chromosomes ^*58*–*61*^ of 7 chromosomal rearrangements that define specific subcategories of AML in the World Health Organisation (WHO) AML classification ^*62*^: t(6;9) *DEK*::*NUP214*, t(8;21) *RUNX1*::*RUNX1T1*, t(9;11) *KMT2A*::*MLLT3*, t(9;22)

*BCR*::*ABL*, t(15;17) *PML*::*RARA*, t(16;16) *CBFB*::*MYH11*, inv(16) *CBFB*::*MYH11* (Supplementary Fig. 5, Supplementary Table 3). *KMT2A* is renowned for having numerous possible breakpoint partners and so additional common breakpoint regions in *KMT2A* were also targeted ^*63*^. *KMT2A*-PTD most commonly involve exons 2 or 3 and span through exon 9 to 11 ^*64*^. They can be detected by observing a relative increase in read depth, starting from exon 2 or 3, compared to an exon that is never involved in *KMT3A*-PTD (e.g. exon 27) ^*65*^. A custom set of 120-nt oligonucleotide baits (TWIST biosciences) was therefore designed to target *KMT2A* exons 2-27 (Supplementary Table 3).

### TETRIS-seq error-corrected library preparation/ sequencing

TETRIS-seq library preparation was performed as per the ‘TWIST custom panel enrichment workflow’ (TWIST Biosciences), but with adaptations to allow for incorporation of duplex unique molecular identifiers (UMIs), for error-corrected sequencing, and to allow for the use of two targeted panels on the same input DNA. DNA yield varied between UKCTOCS samples (Extended Data Figure 1), but for each timepoint the same amount of input DNA was used for both the pre-AML sample and matched control (mean 43 ng, median 50 ng, range 4.5-80 ng).

Briefly, DNA was enzymatically fragmented (using Twist Biosciences’ proprietary enzyme mix), end-repaired and dA-tailed at 32° C for 21-24 min (Supplementary Table 4). IDT xGen™ CS adapters (5.5 ul of 10 *µ*M), containing 3 bp duplex UMI sequences, were then ligated to each end of the DNA fragments at 20° C for 15 min, followed by bead purification using 0.8X DNA Purification beads (Twist Biosciences). The duplex UMI-tagged fragments were PCR amplified in a 50*µ*l reaction volume (25*µ*l KAPA® HiFi HotStart ReadyMix (Roche Diagnostics), 10*µ*l IDT xGen™ unique dual index (UDI) primers (10*µ*M), 15*µ*l duplex UMI-tagged fragments). The following conditions were used for PCR amplification: 98° C for 45s; 11-12 cycles of 98° C for 15s (see Supplementary Table 5), 60° C for 30s, 72° C for 30s; 72° C for 1 min. Following bead purification, using 1X DNA Purification beads (Twist Biosciences), the amplified UMI-tagged dual index-labelled fragments were eluted into 22*µ*l ddH_2_O and then visualised and quantified using the Agilent 2200 TapeStation.

Following quantification, each sample was divided in two; one half for the SNV/ indel panel and the other half for the mCA/ chromosomal rearrangement panel. Samples were pooled together in groups of 8 (187.5 ng of each indexed sample), with separate pools for each of the two panels. The pooled samples were concentrated, using 1.5X Agencourt AMPure XP beads (Beckman Coulter™) and then hybridised to the custom panel probes (SNV/ indel panel probes or mCA/ chromosomal rearrangement panel probes) in a thermocycler at 95° C for 5 min followed by 70° C for 16 hours. Following Streptavidin bead capture (30 min at room temperature), the captured DNA was PCR amplified in a 50*µ*l reaction volume (25*µ*l KAPA® HiFi HotStart ReadyMix (Roche Diagnostics), 2.5*µ*l TWIST amplification primers, 22.5*µ*l captured DNA) under the following conditions: 98° C for 45s; 12 cycles (mCA/ chromosomal rearrangement panel) or 17 cycles (SNV/ indel panel) of 98° C for 15s, 60° C for 30s, 72° C for 30s; 72° C for 1 min. Following bead purification, using 1X DNA purification beads (Twist Biosciences), samples were eluted into 32*µ*l ddH_2_O and then visualised and quantified using the Agilent 2200 TapeStation. Samples were then diluted to 10 nM and submitted for sequencing. Libraries were sequenced on the Illumina NovaSeq 6000 S4 (CRUK Cambridge Institute Genomics Core Facility) using the XP workflow, with 150 bp paired ends reads, 8 bp reads in index 1 and 8 bp reads in index 2. 40 samples were sequenced per lane with 10% PhiX control DNA spiked into each lane. SNV/indel and mCA/ chromosomal rearrangement panels were sequenced in different lanes and, with only a few exceptions, timepoints from the same individual were sequenced on different lanes. Each sequencing run contained ∼ 3 billion reads per lane (∼ 68 million reads per sample). For the SNV/ indel panel (∼ 58 kb, 589 probes), this equated to a raw depth (pre-consensus calling) of ∼ 50,000X. For the mCA/ chromosomal rearrangement panel (∼ 1.6 MB, 13114 probes), this equated to a raw depth of ∼ 1000X.

A custom computational workflow was written for the processing of the sequencing data. This workflow used a number of software packages, including Picard ^*66*^, Fulcrum Genomics fgio package ^*67*^, Burrows-Wheeler Aligner (BWA) ^*68*^, the Genome Analysis Toolkit (GATK) ^*69*^, SAMtools ^*70*^, VarDictJava ^*71*^, Pindel ^*72*^ and ANNOVAR ^*73*^, as well as several custom Python scripts. The workflow consisted of four main steps (Supplementary Note 1C): i) UMI extraction and initial alignment; ii) Single strand consensus sequence (SSCS) calling; iii) Duplex consensus sequence (DCS) calling; and iv) Putative variant detection. For the SNV/ indel panel, variants were called from DCS files. The same depth accuracy was not required for the mCA, *KMT2A PTD* and chromosomal rearrangement panel and so only the SSCS were used for these.

### TETRIS-seq *in silico* noise correction model for SNV calling

The theoretical error rate of duplex sequencing is *<* 10^−9^ errors per bp, which is simply the probability of two complementary errors occurring at the same nucleotide position on both DNA strands, either spontaneously or during the 1st PCR cycle ^*43*^. In reality, however, library preparation artefacts e.g. due to sonication, end-repair and mapping errors ^*44*^ have all been shown to slip through the duplex error correction. Therefore in order to be able to reliably call variants at ultra-low frequency, we developed an *in silico* noise correction method for SNV calling (Supplementary Note 1). Our noise correction model is based on the null hypothesis that each base-change at a position in the panel has a specific error rate (*ϵ*) and the number of variant reads (*k*) observed in a sample at that position would be consistent with beta-binomial sampling at that specific error rate. A custom Python script was written to infer the error rate (*ϵ*) and dispersion (*δ*) values for each possible single nucleotide base-change at each position in the panel (hereafter referred to as simply ‘position’) and samples were called as ‘real’ variants if their variant read count was inconsistent with this distribution of errors. First, samples with a VAF >10% at the position were automatically called as real and excluded from the subsequent analysis. Then, the remaining samples at the position were used to fit a beta-binomial distribution (or a binomial distribution if there were ≤ 3 samples remaining with ≥ 1 variant reads). The beta-binomial (or binomial) *p*-values for all the samples with ≥1 variant reads were calculated and a sample was called as a real variant if it’s *p*-value was less than a *p*-value threshold (6 *×* 10^−6^ which gave a false discovery rate (FDR) of <5% (Supplementary Note 1). When fitting a binomial or beta-binomial error distribution to a position only once, a problem arises if more than one sample at the position has a ‘real variant’. The problem is that the lower VAF variants will not be called as they will be fitted as ‘errors’ within a falsely over-dispersed beta-binomial distribution or a binomial distribution with a falsely elevated error rate. We therefore chose to use an iterative approach for positions observed in haematopoietic and lymphoid tissues in COSMIC v92 ^*58*^ (∼ 2% of sites across our custom panel), where we might expect to see more than one sample with a real variant in our pre-AML cohort. Similarly, if more than one timepoint sample from the same individual was sequenced on the same flow-cell, an iterative approach was used for positions at which one of those samples had been called as a real variant. The iterative approach at the position was continued until no further samples from the individual with multi-timepoint samples were called as real. Our *in silico* noise correction model was applied to all the samples sequenced on the same sequencing lane (∼40 samples) and then additional post-processing filters were applied as detailed in Supplementary Note 1F.

### TETRIS-seq indel, FLT3-ITD, chromosomal rearrangements

Indels were called from DCS files using VarDictJava ^*71*^ and post-processing filters were applied as detailed in Supplementary Note 1. FLT3-ITD mutations were called using Pindel ^*72*^ and were called as real if they were detected in both the SSCS and DCS files. Chromosomal rearrangements were called from SSCS files using Manta ^*74*^.

### TETRIS-seq mCA and *KMT2A*-PTD calling

A custom Python script was written to enable detection of mCAs from the SSCS reads using the SNP ‘B-allele frequencies’ (BAF) and read depths (log_2_R ratios) in the mCA/ chromosomal rearrangement panel (see Supplementary Note 1G). Loss and gain events result in deviations in both LRR and BAF whereas CN-LOH events result in BAF deviations without a change in LRR (because there is no change in the amount of genetic material) (Supplementary Fig. 12). The proportion of cells (‘cell fraction’) harbouring the mCA was calculated from the heterozygous BAFs as detailed in Supplementary Table S1. If an mCA was detected at 100% cell fraction in both the latest and earliest timepoint sample from an individual, then it was deemed likely to be germline. One of the benefits afforded by having longitudinal samples is the ability to use large BAF deviations from higher-cell fraction mCAs, detected at timepoints closer to AML diagnosis, to identify which SNPs lie on the same chromosome in the affected region of interest. This phasing information can then be applied to the same individual’s samples from earlier timepoints, allowing a much higher sensitivity for detection of the mCA when it is at lower cell fraction (Supplementary Fig. 13). Using this approach we were able to call mCAs at cell fractions as low as 0.1%.

To detect KMT2A-PTD events a custom Python script was written (using pysam ^*75*^ pileup) to calculate the mean read depth across each targeted KMT2A exon. These mean depths were then normalised to the mean read depth across KMT2A exon 27 and the exon 3:27 ratio (*R*) calculated. The fraction of cells harbouring the KMT2A-PTD can then be calculated as: cell fraction = 2(R - 1) (Supplementary Note 1H).

### Inferring clonal architecture

A hierarchical step-wise process was used to infer whether a variant was likely the only driver mutation in a clone (i.e a ‘single-mutant clone’) or whether it was more likely in a clone with one or more of the other variants detected in that individual (see Supplementary Table 6). First we looked at the largest cell fractions for each of the variants detected in an individual: using the pigeon-hole principle, if the sum of the cell fractions for two variants was >100% at any timepoint then these two variants were inferred to be in the same clone. Second, we looked at the cell fraction dynamics of the variant trajectories: if the cell fraction trajectories crossed-over at any timepoint (i.e. one variant’s cell fraction increased to above the cell fraction of another) then these two variants were inferred to be in separate clones. Third, we looked at the correlation in cell fraction dynamics between variants: if the trajectories of two variants showed similar growth dynamics (e.g. increasing or decreasing in cell fraction at the same timepoints) then these variants were inferred to be in the same clone. Next, if a larger clone was already present when a new variant was acquired, we inferred that this new variant was more likely to have arisen within the existing larger clone if the larger clone’s cell fraction was >50% when the new variant was acquired. Finally, in particular when considering variants that were only detected at timepoints closest to time of AML diagnosis, but which were still present at low cell fraction, we looked at the growth rate of the variant and it’s timing relative to AML diagnosis: the higher the growth rate, and the closer to AML diagnosis the variant was first detected, the more likely we were to infer that the variant was associated with a multiple-mutant clone.

### Inferring mutation acquisition age and fitness effect

We developed a mathematical framework for inferring the acquisition age and fitness effects of clones. In this framework each clone (defined by its most recently acquired mutation) has an ‘establishment time’ (*t*_0_) and fitness effect (*s*), which determine the clone’s size over time, such that at a particular time (*t*_1_), the clone size is given by the equation:

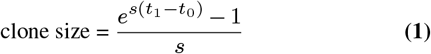

The frequencies of all clones in the population (including non-mutated HSCs) sums to 1, such that at a particular time (*t*_1_), the clone frequency is given by the equation:

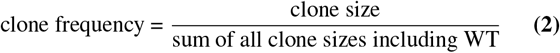

The dynamics of a specific *variant*, which is directly measured via the variant’s VAF, are determined by summing the frequency of the original parental clone (i.e. the first clone to acquire the variant) and all daughter clones which inherited the variant. Using this framework, together with inferences for the clonal architecture of each sample (as described above), we developed a maximum likelihood approach to infer fitness and establishment time estimates for each clone in the population by comparing the predicted VAF trajectories to the observed VAF trajectories in all pre-AML cases and controls. In order to assign a likelihood to the observed data, given the predicted trajectories, we developed a likelihood function as follows: for each variant at each timepoint, we assigned a likelihood to the observed number of variant reads using a binomial probability with mean given by the predicted frequency and sample size given by the total depth at that position in the sample. The log(likelihoods) were summed across all the variant timepoints for all trajectories and our maximum likelihood approach minimised the sum of all the total log(likelihoods).

Our maximum likelihood optimisation approach started with an ‘initial estimate’ for the establishment time and fitness of each clone. Depending on the number of VAF timepoints, this was typically estimated from the gradient of a straight line drawn through the log(VAF) trajectory (with or without ‘bounded VAFs’ for undetectable VAF timepoints) or was based on the number of mutations in the clone (see Supplementary Note 3). Once the ‘initial estimate’ for the establishment time and fitness of each clone had been set, a 3-stage maximum likelihood process was employed. Within each stage ∼1000-5000 optimisations were performed, with adjustment of the establishment time and fitness with each optimisation in order to find the lowest total log(likelihood) across all trajectories in that sample. Each of the 3 stages involved progressively finer adjustments in the predicted establishment time and fitness of the clone with each optimisation (similar to simulated annealing). Penalties were applied to the log(likelihoods) in certain situations, e.g. to account for timepoints at which a variant was undetectable (see Supplementary Note 3). For each sample, the 3-stage maximum likelihood process was repeated for 25 ‘seeds’ and the results of the seed that produced the lowest total log(likelihood) were taken as the most likely establishment times and fitnesses for the variants in that sample.

### A simulated model of evolutionary dynamics in blood

In this model, blood is maintained by a population if *N* cycling HSCs that divide symmetrically every *τ* years, modelled as a continuous time birth death process ^*38*^. HSCs acquire driver mutations at a rate *U* per year. Driver mutations vary in the fitness effects they confer to HSCs. The fitness effects are drawn from a probability density distribution which is modelled using a stretched exponential as described previously ^*39*^

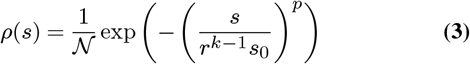

where 𝒩 is a normalisation constant, *k* is the number of driver mutations in the clone and where we have set *p* = 3, *r* = 3. A driver mutation arising in a wildtype HSC, creating a single mutant clone, confers fitness advantages drawn from a distribution with a median and range controlled by a single parameter *s*_0_ =11% per year. A driver mutation arising in single-mutant clone confers a median fitness effect that is *r* = 3-fold larger (typical fitness effects *s* = 33% per year) creating a double-mutant clone. Triple and quadruple-mutant clones follow the same geometric pattern (with median fitness effects of *s* ∼100% and *s* ∼ 300% per year respectively). Fitness advantages of driver mutations combine additively in this model, such that the growth rate of a clone carrying *k* driver mutations is given by

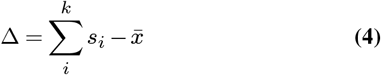

where 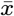 is the mean fitness of all clones in the population weighted by their clone size and where *s*_*i*_ are the fitness advantages of each of the driver mutations in the clone. Transformation to frank AML is defined by the acquisition of 4 driver mutations within the same clone that reaches >50% cell fraction before the age of 85. Using established estimates for *Nτ* = 10^5^, *U* ∼ 10^−5^ per year and *s* ∼ 11% per year for single-mutant fitness effects, we simulated clonal dynamics in 25,000 simulations and considered the evolutionary dynamics that is observed in the decades preceding the ∼ 100 virtual incidental AMLs.

### High VAF variants in UK Biobank

Clonal haematopoiesis variants were called in a random sample of 335,090 individuals from UK Biobank, excluding any individuals with a history of cancer before the UK Biobank baseline assessment (including self-reported cancer, excluding basal cell carcinoma, pre-cancer of the cervix and rodent ulcer). The North West Multi-centre Research Ethics Committee (MREC) reviewed and approved the UK Biobank scientific protocol and operational procedures (REC 21/NW/0157) and all participants provided signed informed consent at enrolment. Variants were called at 23 genomic positions previously identified as the locations of clonal haematopoiesis driver variants ^*38*^ from the UK Biobank exome CRAM files using *samtools mpileup* on the DNANexus Research Analysis Platform (UK Biobank Application Number 28126): *DNMT3A* R320C, R326C, R598*, R729W, Y735C, R736C/H, R771*, R882H/C/S, W860R, P904L; *GNB1* K57E; *IDH1* R132H; *IDH2* R140Q, R172K; *JAK2* V617F; *KIT* D816F/H/V/Y; *KRAS* G12C/D/V; *MPL* W515L/*; *NPM1* W288fs; *NRAS* G12C/D/V; *SF3B1* K666N, K700E; *SRSF2* P95H/R. Variants with <3 supporting reads were discarded. 410 individuals were identified who were diagnosed with AML after the UK Biobank baseline assessment (‘prospective cases’). Individuals who developed other haematological malignancies after UK Biobank baseline assessment were excluded (*n* = 3,164). 3,678 individuals carried at least one of the clonal haematopoiesis variants analysed, of whom 83 were prospective AML cases. 72 individuals carried two CH driver variants, of whom 14 were prospective AML cases.

### Statistics & Reproducibility

No statistical methods were used to predetermine sample size: all individuals in the ‘multimodal arm’ of UKCTOCS who had been diagnosed with AML by 2018, and who had sufficient DNA in their serum for analysis, were included.

## Data availability

### Code availability

All code used in this study will be made available on the Blundell lab Github page:

#### Acknowledgements

We thank Daniel Fisher, Ivana Cvijovic, Sarah Seton-Rogers, Alex Frankell, Siddhartha Kar, Andrew King, George Vassiliou, Inigo Martincorena and all members of the Blundell lab for helpful comments on this work. C.J.W and J.R.B are supported by the Early Cancer Institute, the CRUK Cambridge Centre and the NIHR Biomedical Research Centre. J.R.B. is supported by a UKRI Future Leaders Fellowship (MR/S031782/1). C.J.W. was supported by a CRUK Clinical Research Fellowship and is currently supported by a Wellcome Early Career Award (226929/Z/23/Z). H.A.J.M. is supported by the International Alliance for Cancer Early Detection, an alliance between Cancer Research UK (C14478/A29329), Canary Center at Stanford University, the University of Cambridge, OHSU Knight Cancer Institute, University College London and the University of Manchester. U.M., S.A. and A.G-M. are supported by the NIHR UCL Hospitals Biomedical Research Centre and by Medical Research Council Clinical Trials Unit at UCL core funding (MR_UU_12023). We thank the UKCTOCS participants, without whom this study would not have been possible, and everyone involved in the conduct and oversight of UKCTOCS. UKCTOCS was funded by Medical Research Council (G9901012 and G0801228), Cancer Research UK (C1479/A2884), and the UK Department of Health, with additional support from The Eve Appeal. The long-term follow-up UKCTOCS was supported by National Institute for Health Research (NIHR HTA grant 16/46/01), Cancer Research UK, and The Eve Appeal. The analysis of UK Biobank data was conducted using the UK Biobank Resource under Application Number 28126. None of the funders had a role in data analysis, decision to publish or preparation of the manuscript.

## Author contributions

J.R.B conceived of the study. J.R.B. and C.J.W designed the study with input from U.M, A.G-M. and S.A. J.R.B and C.J.W developed the sequencing strategy (TETRIS-seq). C.J.W generated the libraries and developed the bioinformatic processing pipelines with input from J.R.B. J.R.B and C.J.W developed the evolutionary framework for estimating fitness and occurrence time with input from Y.P.G.P. Simulations and models were developed by J.R.B with input from Y.P.G.P and C.J.W. UK Biobank analysis was performed by H.A.J.M with input from J.R.B and C.J.W. Manuscript was written by J.R.B and C.J.W with input from all authors.

## Extended Data Figures

**Extended Data Fig. 1.**
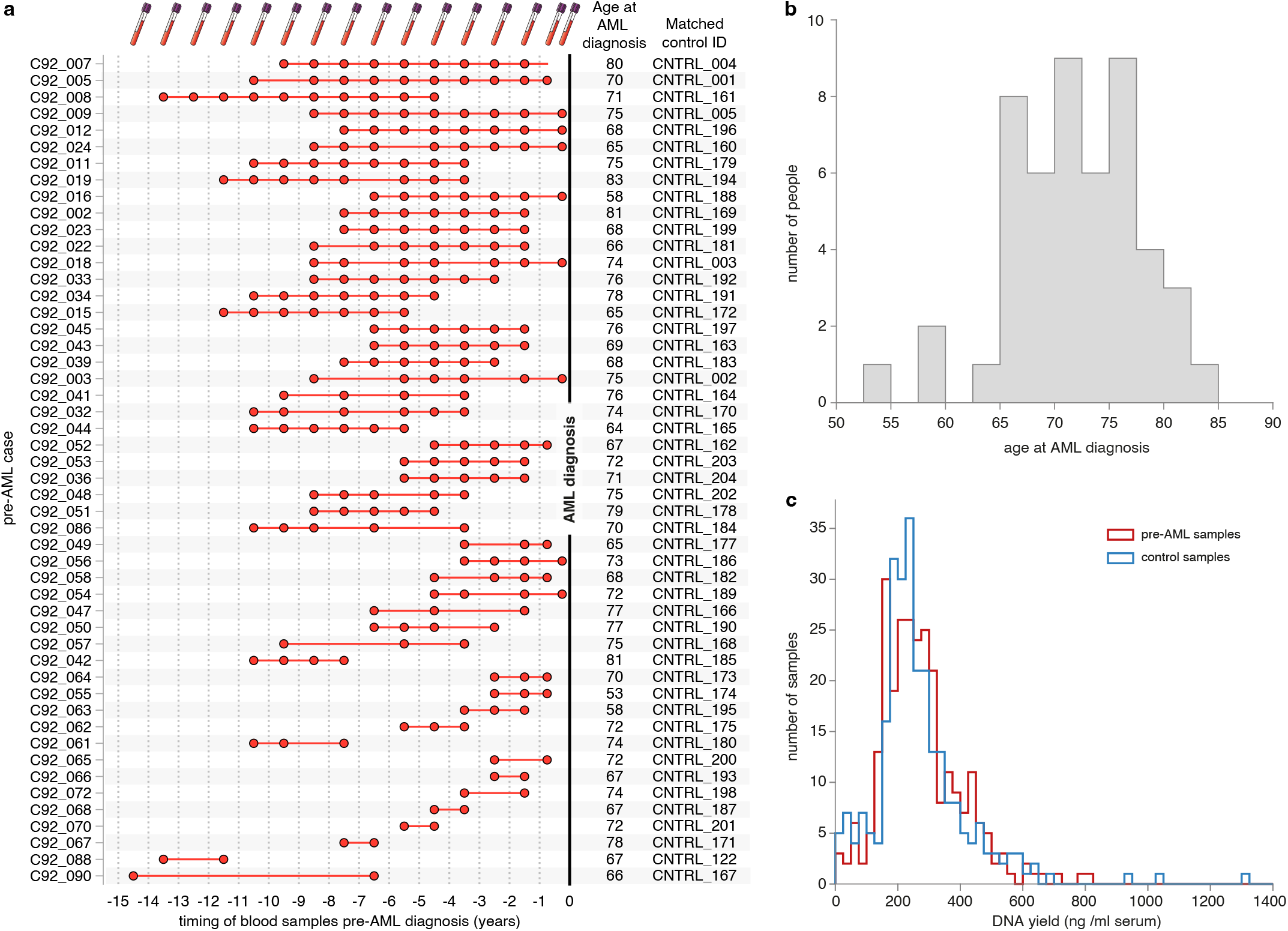
Longitudinal blood samples pre-AML diagnosis. **a**. Timings of blood samples in relation to AML diagnosis for all individuals in UKCTOCS who had multiple blood samples collected pre-AML diagnosis. **b**. Distribution of age at AML diagnosis for individuals in UKCTOCS: mean age 71 years, range 53-83 years, s.d. 6 years. **c**. Distribution of DNA yields from pre-AML serum samples (red histogram) and control serum samples (blue histogram). Mean DNA yield: 273 ng/ml in pre-AMLs, 269 ng/ml in controls.

**Extended Data Fig. 2.**
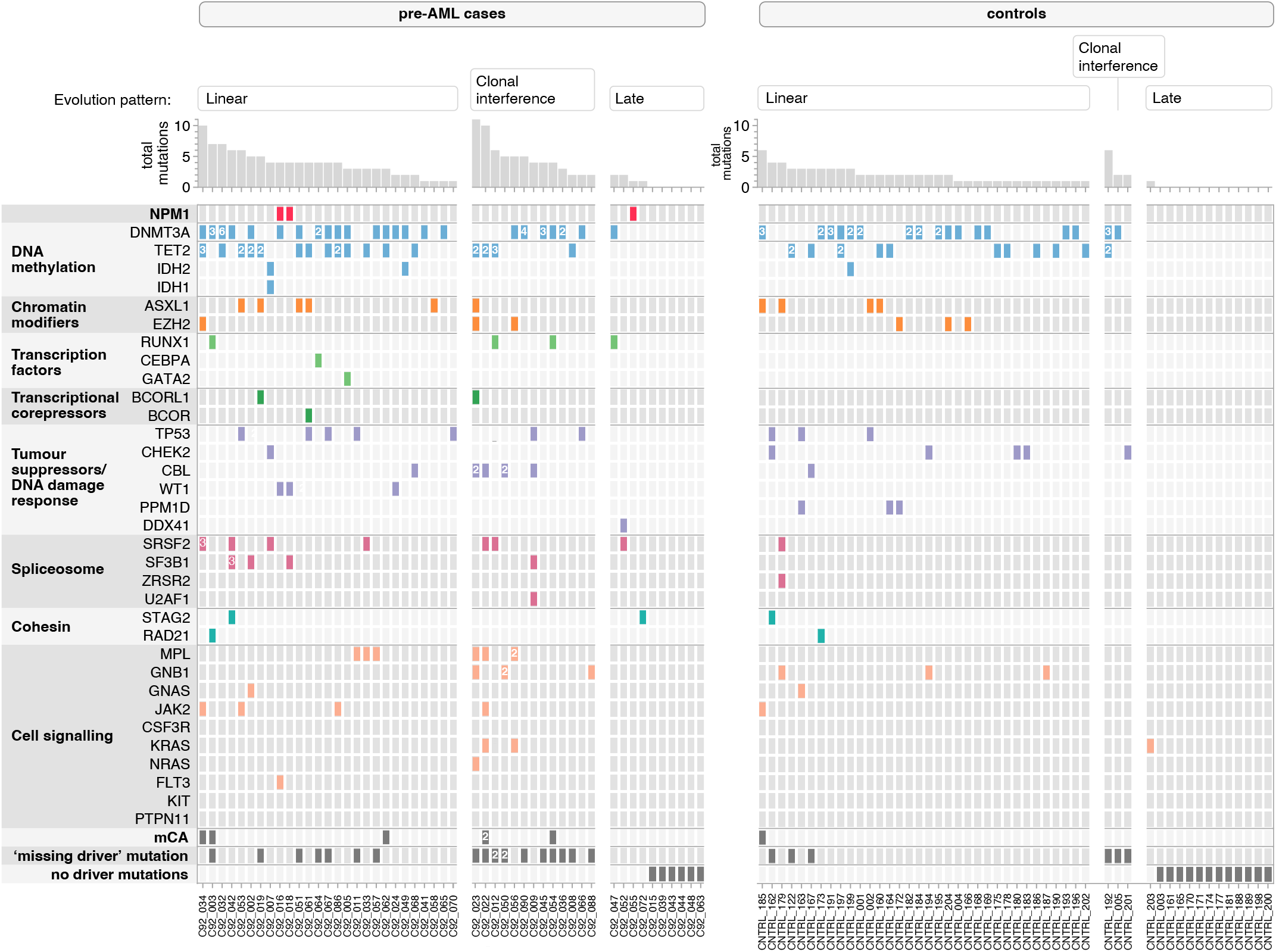
Classes of mutations detected in pre-AML and control samples. Each column represents a pre-AML case or a control. Coloured boxes indicate the mutations detected in each individual. If multiple mutations were detected within the same gene within an individual the number detected is written within the coloured box.

**Extended Data Fig. 3.**
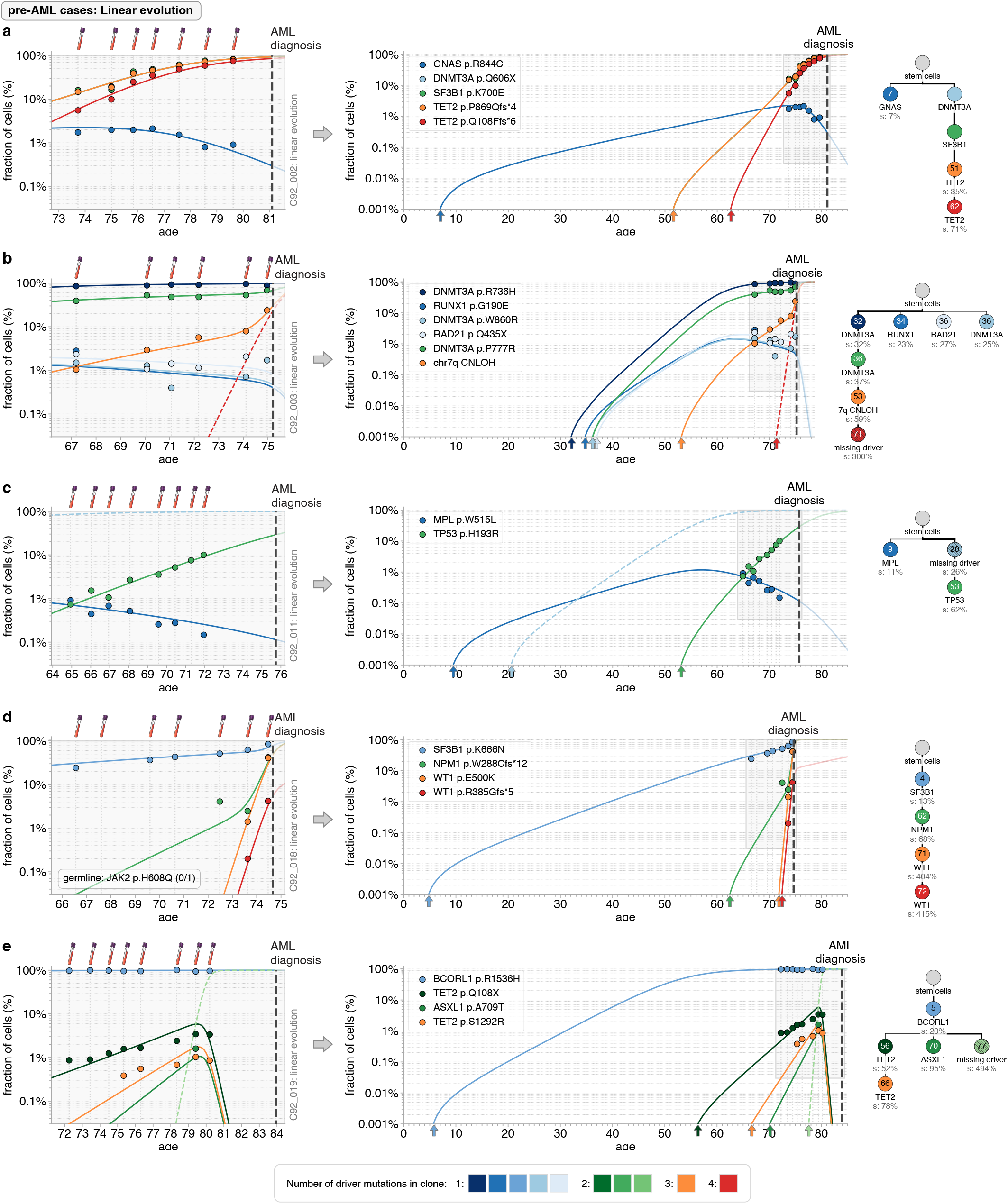
Quantitative dynamics of driver mutations in the decades before AML diagnosis: pre-AML cases with linear evolution part 1. **a-e**. Observed variant frequency trajectories (data points) compared with predicted variant frequency trajectories (coloured lines) due to the most likely fitness and occurrence time estimates of clones, across the period of UKCTOCS blood sampling (left-hand-side plots) and since the birth of the individual (right-hand-side plots). Trajectories of any inferred ‘missing drivers’ are indicated by dashed coloured lines. Grey vertical dashed lines indicate timing of blood samples. Thick black vertical dashed line indicates time of AML diagnosis. Phylogenetic trees (right-hand-side) show the inferred clonal composition of the population, occurrence times of each clone (number in circles, years) and their fitness effect (% per year). The number of driver mutations in the clone is indicated by its colour (see legend).

**Extended Data Fig. 4.**
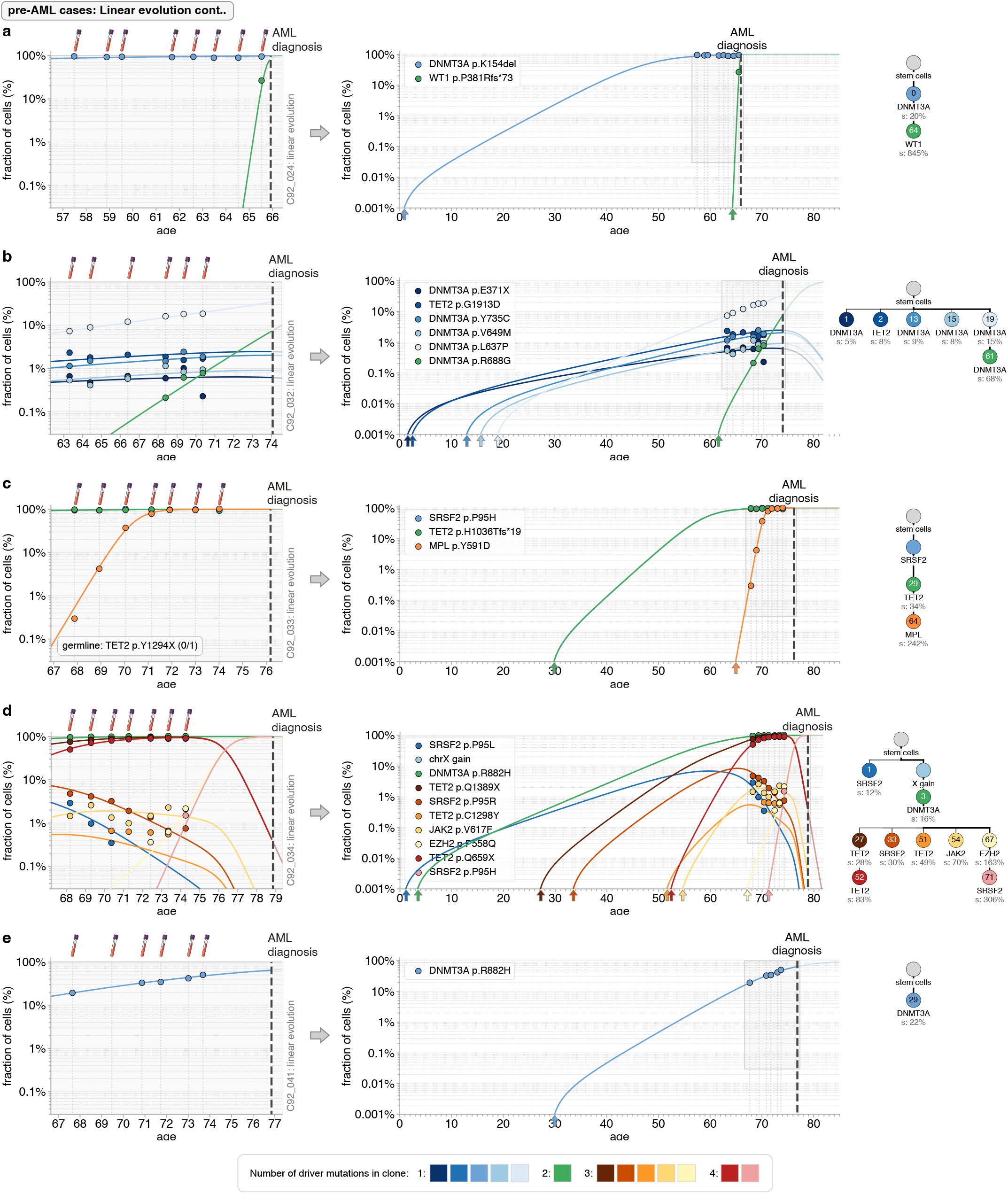
Quantitative dynamics of driver mutations in the decades before AML diagnosis: pre-AML cases with linear evolution part 2. **a-e**. Observed variant frequency trajectories (data points) compared with predicted variant frequency trajectories (coloured lines) due to the most likely fitness and occurrence time estimates of clones, across the period of UKCTOCS blood sampling (left-hand-side plots) and since the birth of the individual (right-hand-side plots). Trajectories of any inferred ‘missing drivers’ are indicated by dashed coloured lines. Grey vertical dashed lines indicate timing of blood samples. Thick black vertical dashed line indicates time of AML diagnosis. Phylogenetic trees (right-hand-side) show the inferred clonal composition of the population, occurrence times of each clone (number in circles, years) and their fitness effect (% per year). The number of driver mutations in the clone is indicated by its colour (see legend).

**Extended Data Fig. 5.**
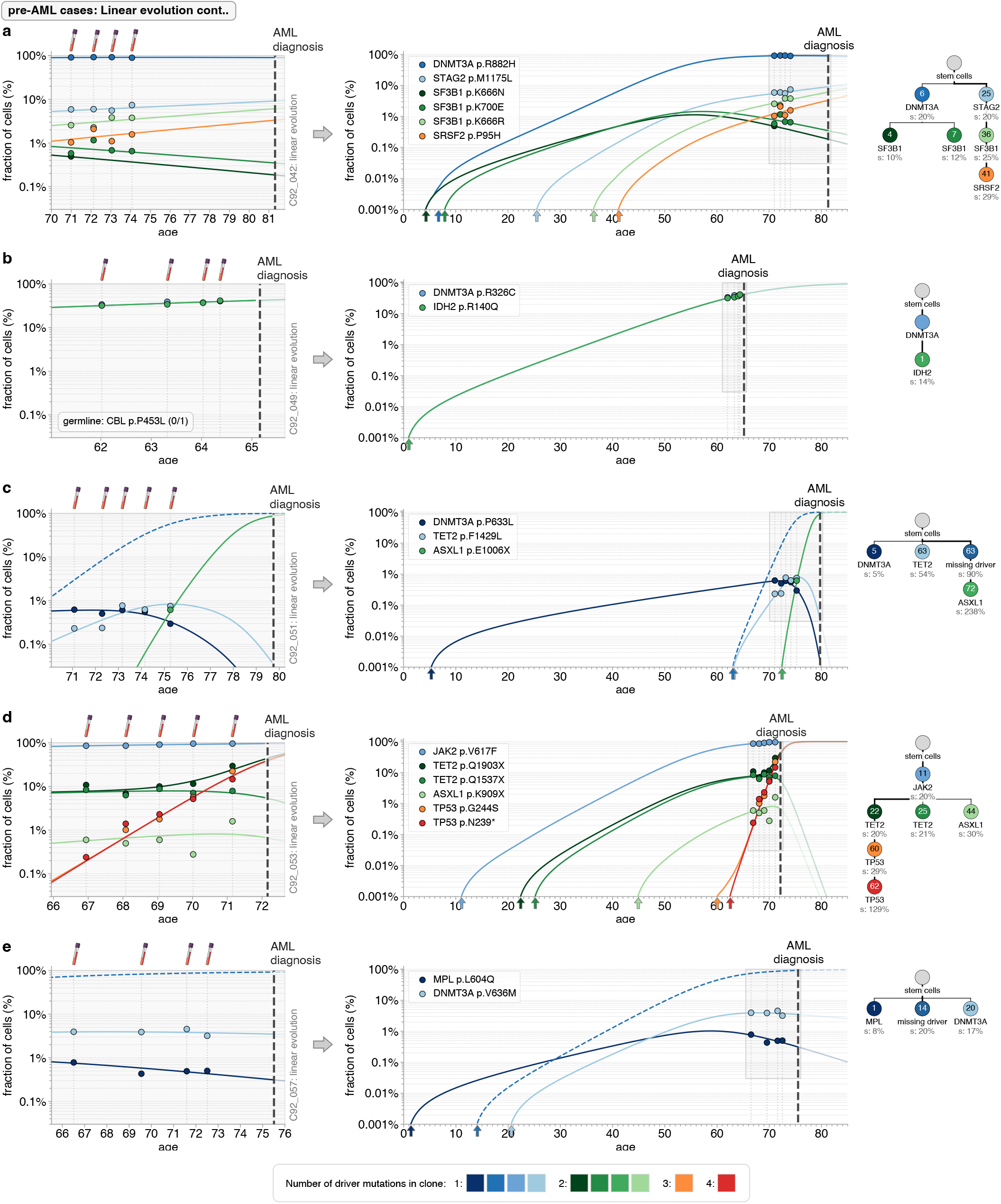
Quantitative dynamics of driver mutations in the decades before AML diagnosis: pre-AML cases with linear evolution part 3. **a-e**. Observed variant frequency trajectories (data points) compared with predicted variant frequency trajectories (coloured lines) due to the most likely fitness and occurrence time estimates of clones, across the period of UKCTOCS blood sampling (left-hand-side plots) and since the birth of the individual (right-hand-side plots). Trajectories of any inferred ‘missing drivers’ are indicated by dashed coloured lines. Grey vertical dashed lines indicate timing of blood samples. Thick black vertical dashed line indicates time of AML diagnosis. Phylogenetic trees (right-hand-side) show the inferred clonal composition of the population, occurrence times of each clone (number in circles, years) and their fitness effect (% per year). The number of driver mutations in the clone is indicated by its colour (see legend).

**Extended Data Fig. 6.**
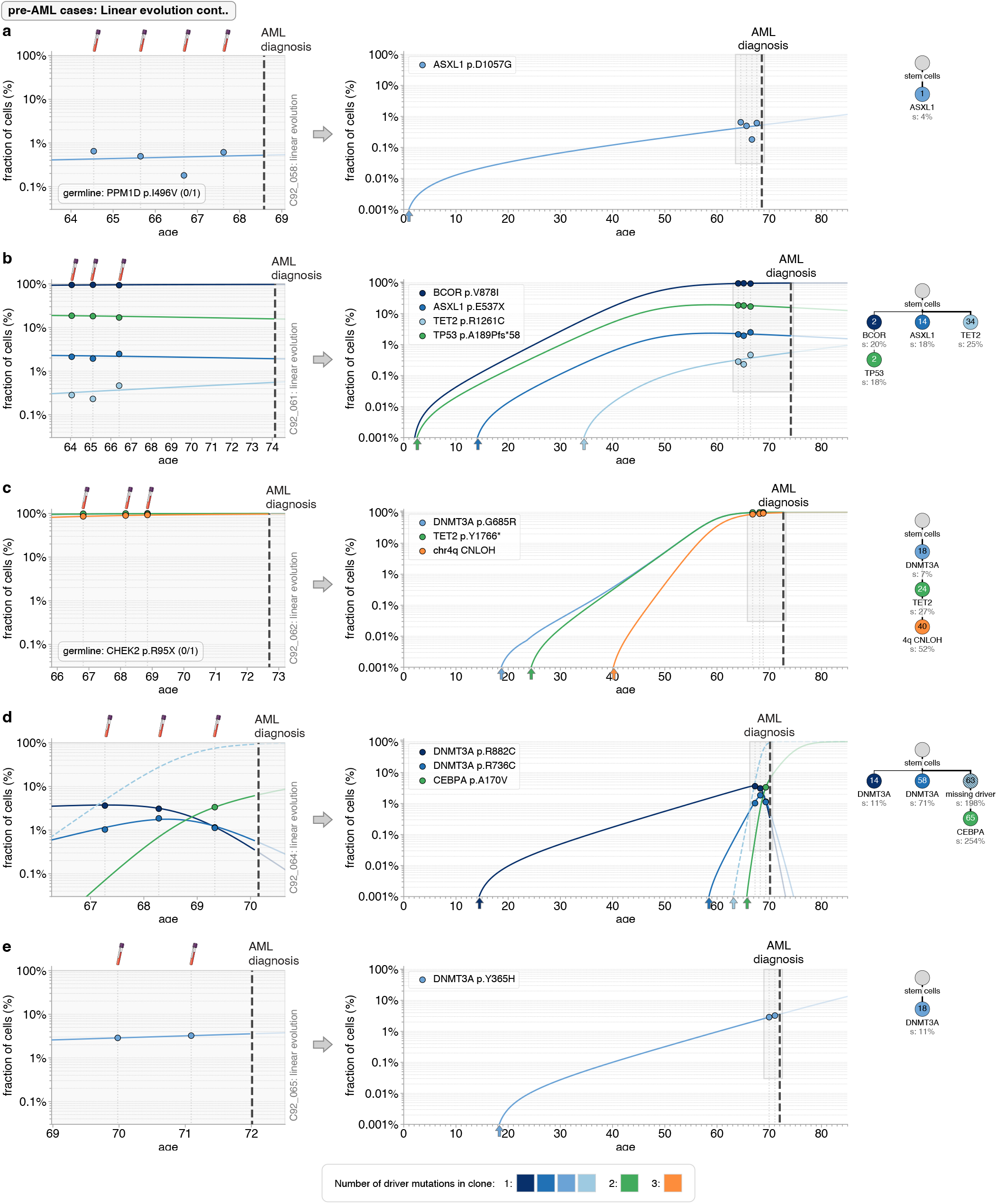
Quantitative dynamics of driver mutations in the decades before AML diagnosis: pre-AML cases with linear evolution part 4. **a-e**. Observed variant frequency trajectories (data points) compared with predicted variant frequency trajectories (coloured lines) due to the most likely fitness and occurrence time estimates of clones, across the period of UKCTOCS blood sampling (left-hand-side plots) and since the birth of the individual (right-hand-side plots). Trajectories of any inferred ‘missing drivers’ are indicated by dashed coloured lines. Grey vertical dashed lines indicate timing of blood samples. Thick black vertical dashed line indicates time of AML diagnosis. Phylogenetic trees (right-hand-side) show the inferred clonal composition of the population, occurrence times of each clone (number in circles, years) and their fitness effect (% per year). The number of driver mutations in the clone is indicated by its colour (see legend).

**Extended Data Fig. 7.**
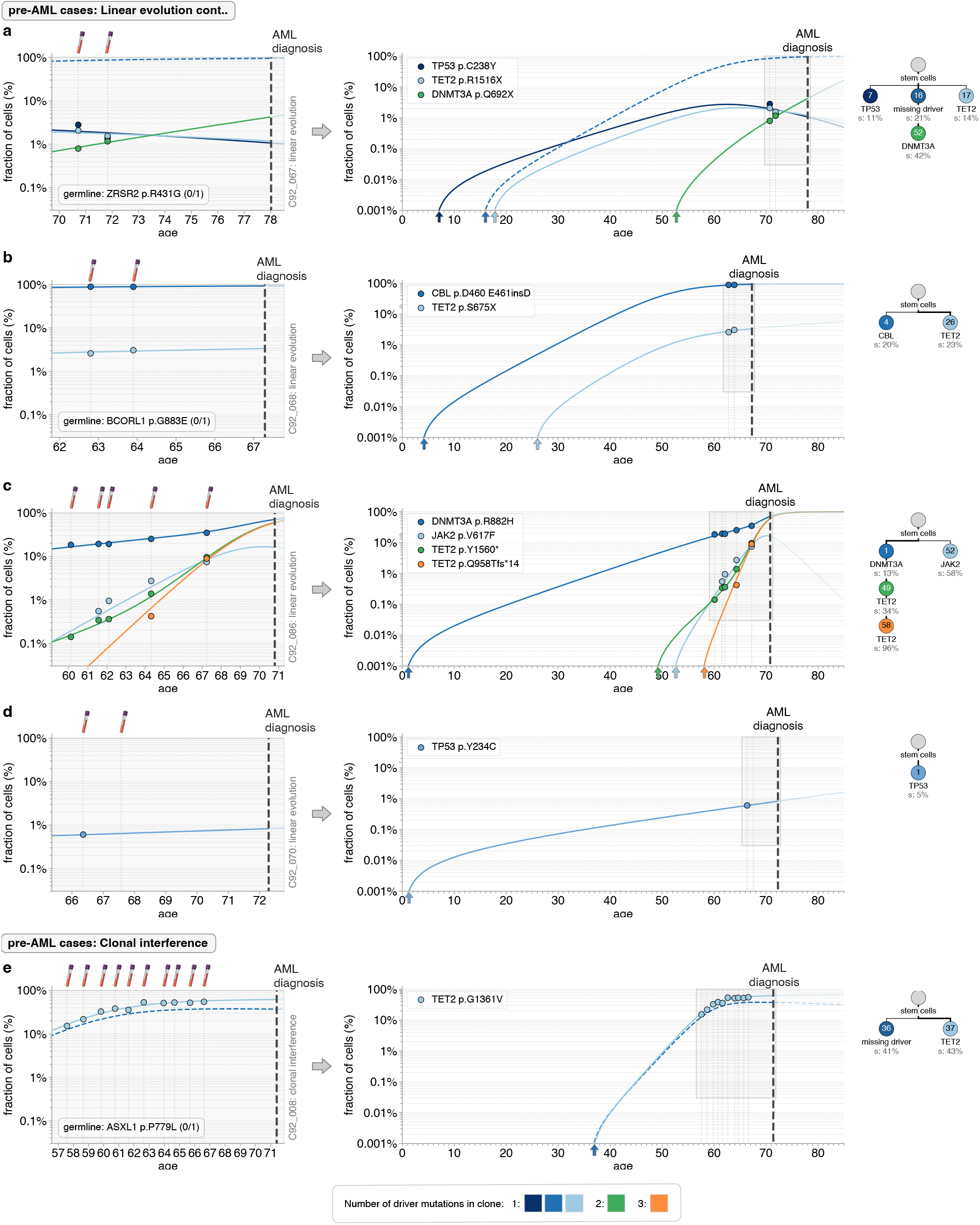
Quantitative dynamics of driver mutations in the decades before AML diagnosis: pre-AML cases with linear evolution part 5 and Clonal interference part 1. **a-e**. Observed variant frequency trajectories (data points) compared with predicted variant frequency trajectories (coloured lines) due to the most likely fitness and occurrence time estimates of clones, across the period of UKCTOCS blood sampling (left-hand-side plots) and since the birth of the individual (right-hand-side plots). Trajectories of any inferred ‘missing drivers’ are indicated by dashed coloured lines. Grey vertical dashed lines indicate timing of blood samples. Thick black vertical dashed line indicates time of AML diagnosis. Phylogenetic trees (right-hand-side) show the inferred clonal composition of the population, occurrence times of each clone (number in circles, years) and their fitness effect (% per year). The number of driver mutations in the clone is indicated by its colour (see legend).

**Extended Data Fig. 8.**
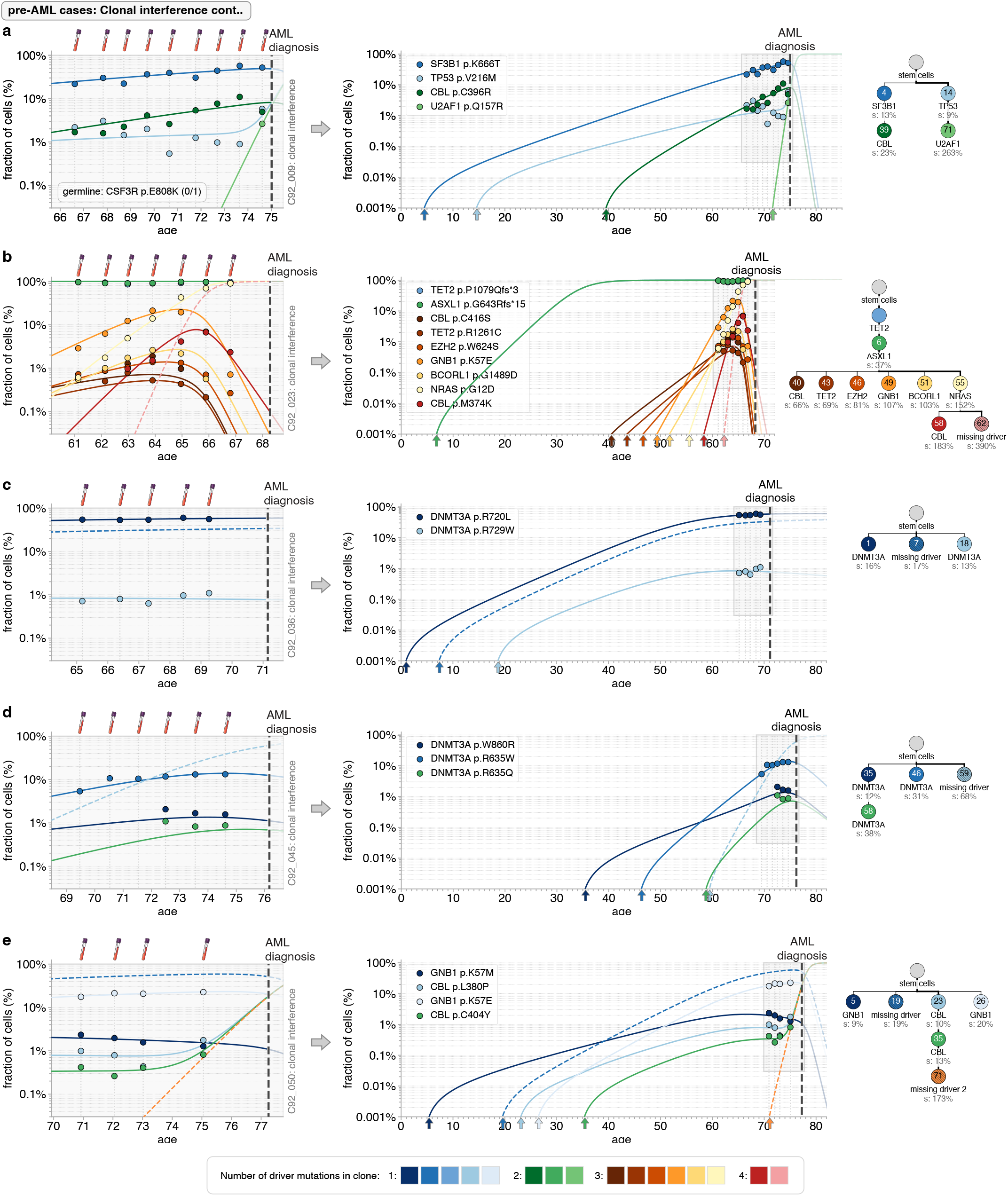
Quantitative dynamics of driver mutations in the decades before AML diagnosis: pre-AML cases with clonal interference part 2. **a-e**. Observed variant frequency trajectories (data points) compared with predicted variant frequency trajectories (coloured lines) due to the most likely fitness and occurrence time estimates of clones, across the period of UKCTOCS blood sampling (left-hand-side plots) and since the birth of the individual (right-hand-side plots). Trajectories of any inferred ‘missing drivers’ are indicated by dashed coloured lines. Grey vertical dashed lines indicate timing of blood samples. Thick black vertical dashed line indicates time of AML diagnosis. Phylogenetic trees (right-hand-side) show the inferred clonal composition of the population, occurrence times of each clone (number in circles, years) and their fitness effect (% per year). The number of driver mutations in the clone is indicated by its colour (see legend).

**Extended Data Fig. 9.**
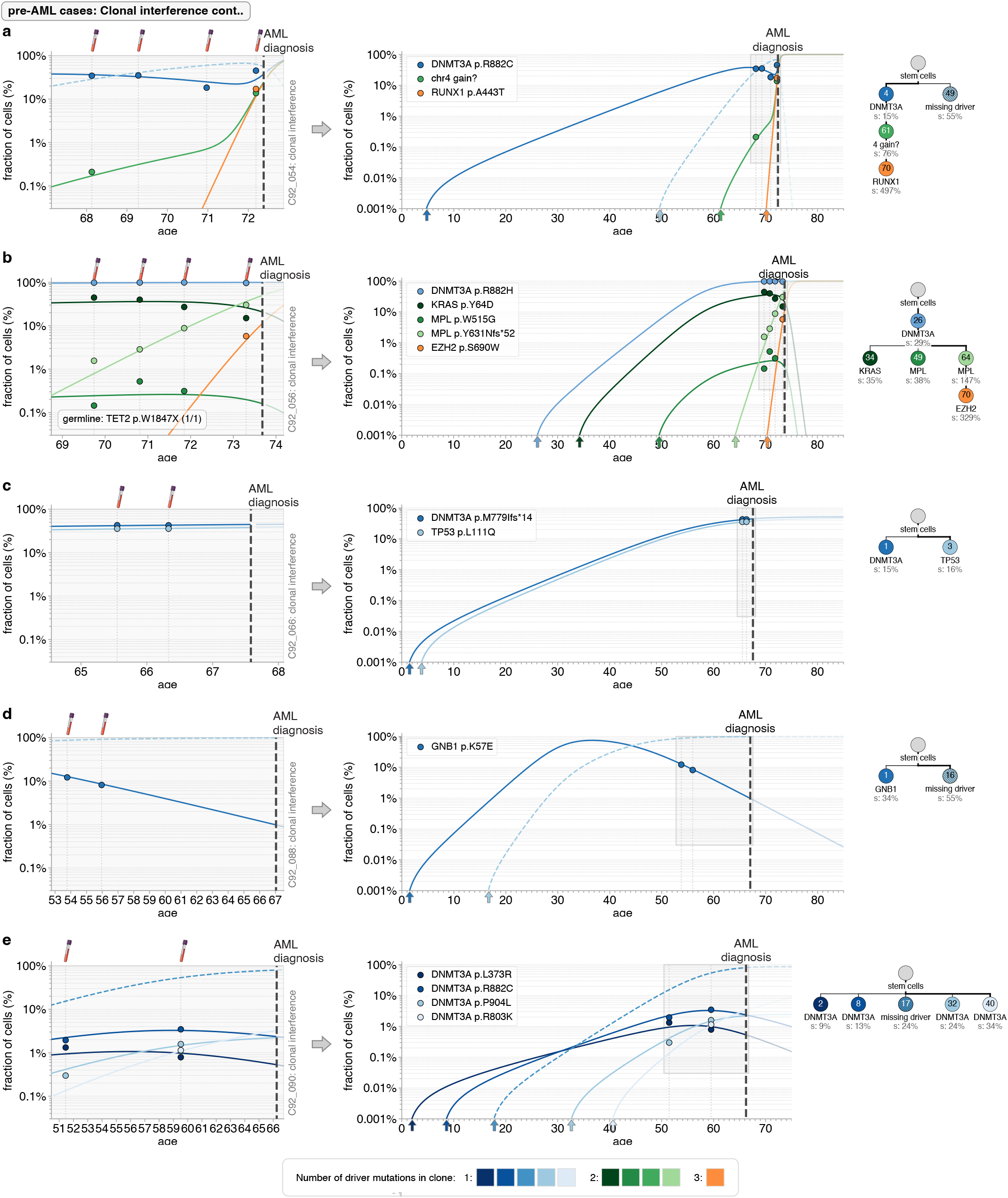
Quantitative dynamics of driver mutations in the decades before AML diagnosis: pre-AML cases with clonal interference part 3. **a-e**. Observed variant frequency trajectories (data points) compared with predicted variant frequency trajectories (coloured lines) due to the most likely fitness and occurrence time estimates of clones, across the period of UKCTOCS blood sampling (left-hand-side plots) and since the birth of the individual (right-hand-side plots). Trajectories of any inferred ‘missing drivers’ are indicated by dashed coloured lines. Grey vertical dashed lines indicate timing of blood samples. Thick black vertical dashed line indicates time of AML diagnosis. Phylogenetic trees (right-hand-side) show the inferred clonal composition of the population, occurrence times of each clone (number in circles, years) and their fitness effect (% per year). The number of driver mutations in the clone is indicated by its colour (see legend).

**Extended Data Fig. 10.**
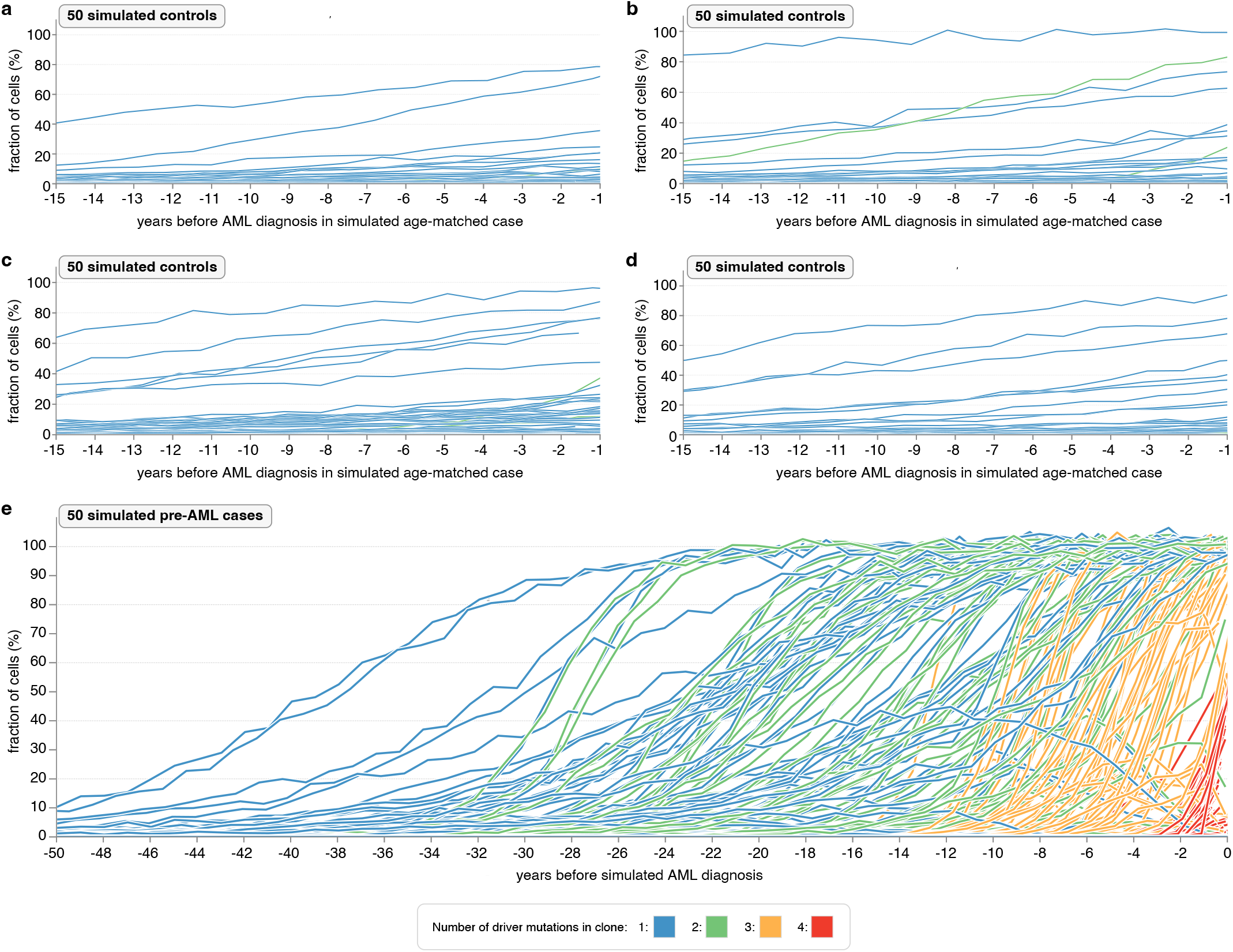
(**a–d**). Cell fraction trajectories from 4 different sets of 50-age matched simulated controls in the 15 years prior to diagnosis of AML in age- and timepoint-matched simulated individuals who were subsequently diagnosed with AML. In age-matched controls large single-mutant clones are not uncommon, but large double-mutant clones are rare. **e**. Cell fraction trajectories from 142 virtual AMLs from 25,000 simulated individuals in the 50 years prior to simulated AML diagnosis. Driver variants that arise in multiple-mutant clones are denoted by colour: single- (blue), double- (green), triple- (orange) and quadruple-mutants (red). Early single and double mutant sweeps are a common feature of future virtual AMLs.

## Supplementary Notes

### 1. TETRIS-seq: Comprehensive error-corrected sequencing with *in silico* noise correction

#### A. Targeted SNV/ indel panel

To design a custom panel for targeting gene mutations, we chose the most commonly mutated genes from 9 different clonal haematopoiesis studies ^*18,19,21,36,76*–*80*^, as well as genes recurrently mutated in AML (e.g. NPM1, FLT3) ^*81*^ and DDX41, in which both inherited and somatic variants can be associated with an increased risk of AML ^*82*^. We then looked at the distribution of variants across these genes, in both clonal haematopoiesis and AML, and specifically targeted the exons where variants were most commonly found (Supplementary Figs. 1-4, Supplementary Table 1).

#### B. Targeted panel for mCAs, KMT2A-PTD and chromosomal rearrangements

##### Mosaic chromosomal alterations (mCAs)

To detect mCAs, we used an approach analogous to SNP microarrays, involving the targeting of common SNPs evenly spaced across the genome (a ‘SNP backbone’). This method allows heterozygous SNP B-allele frequency (BAF) deviations to be detected which, when combined with read depth information, enables detection of gain, loss or CN-LOH events. In order to design a custom set of 120-nt oligonucleotide baits for the ‘SNP backbone’ (TWIST Biosciences), common SNPs (minor allele frequency (MAF) > 0.01) were downloaded from UCSC dbSNP (release 153, hg19). A custom Python script was then written to filter out SNPs that were likely to be uninformative as follows:

- Heterozygous SNPs are essential for BAF deviations and so, to maximise their number, SNPs were filtered to retain only those whose MAF was 0.40 - 0.45 in 1000 genomes ^*83*^.
- Previous work has found significant between-sample variation in normalised read depths if regions with low GC (≤30%) or high GC (≥60%) are targeted ^*84*^, which can make interpretation of copy number change difficult. The GC content of the region +/-60bp of each of the SNPs was therefore calculated using Pybedtools ^*85*^ and SNPs were excluded if the GC content was not between 35-55%.
- Mapping artefacts can result in inaccurate BAF and read depth measurements and so, to reduce the risk of this, SNPs were excluded if their surrounding region (+/-60bp) mapped to more than one location (using Bowtie2 ^*86*^ or overlapped with highly repetitive regions in Repeatmasker (hg19).

A minimum of 5 SNPs are typically needed to call an mCA and so the smaller the gap between targeted SNPs, the higher the resolution for detecting shorter mCAs ^*84*^. With better length resolution comes greater panel size, however, which will result in lower depth and therefore poorer resolution for small BAF deviations associated with low cell fraction mCAs. To strike a balance between the two, we targeted a total of 10,326 SNPs which were spaced every ∼ 280 kb across the genome (Supplementary Table 2), allowing us to detect mCAs as short as ∼ 1.5 MB.

##### KMT2A partial tandem duplications (KMT2A-PTD)

Partial tandem duplications in *KMT2A* (*KMT2A*-PTD) are found in 5-10% of adult *de novo* AML and most commonly involve exons 2 or 3 and span through exon 9 to 11 ^*64*^. They can be detected by observing a relative increase in read depth, starting from exon 2 or 3, compared to an exon that is never involved in *KMT2A*-PTD (e.g. exon 27) ^*65*^. A custom set of 120-nt oligonucleotide baits (TWIST biosciences) was therefore designed to target *KMT2A* exons 2-27 (Supplementary Table 3).

##### Chromosomal rearrangements

We targeted 7 chromosomal rearrangements that define specific subcategories of AML in the World Health Organisation (WHO) AML classification ^*62*^: t(6;9) DEK::NUP214, t(8;21) RUNX1::RUNX1T1, t(9;11) KMT2A::MLLT3, t(9;22) BCR::ABL, t(15;17) PML::RARA, t(16;16) CBFB::MYH11, inv(16) CBFB::MYH11. A custom set of 120-nt oligonucleotide baits (TWIST biosciences) was designed to target the known breakpoint regions of each of the rearranged partner chromosomes ^*58*–*61*^. KMT2A is renowned for having numerous possible breakpoint partners and so additional common breakpoint regions in KMT2A were also targeted ^*63*^. Overall, coverage of the intended target regions was good, although the presence of highly repetitive sequences meant there were coverage gaps in some target regions (Supplementary Fig. 5, Supplementary Table 3). This should not be an issue as long as the rearranged partner’s breakpoint is covered by the panel because the non-targeted breakpoint region will be present as part of a ‘chimeric sequence’, which will be captured by the baits targeting the partner chromosome. It should therefore be possible to identify both breakpoint partners, even if only one was targeted by the panel.

The total size of the custom panel targeting mCAs, KMTA-PTD and chromosomal rearrangements was 1,631,472 bp (∼ 1.6 MB), which was covered by 13,114 probes.

**Supplementary Fig. 1.**
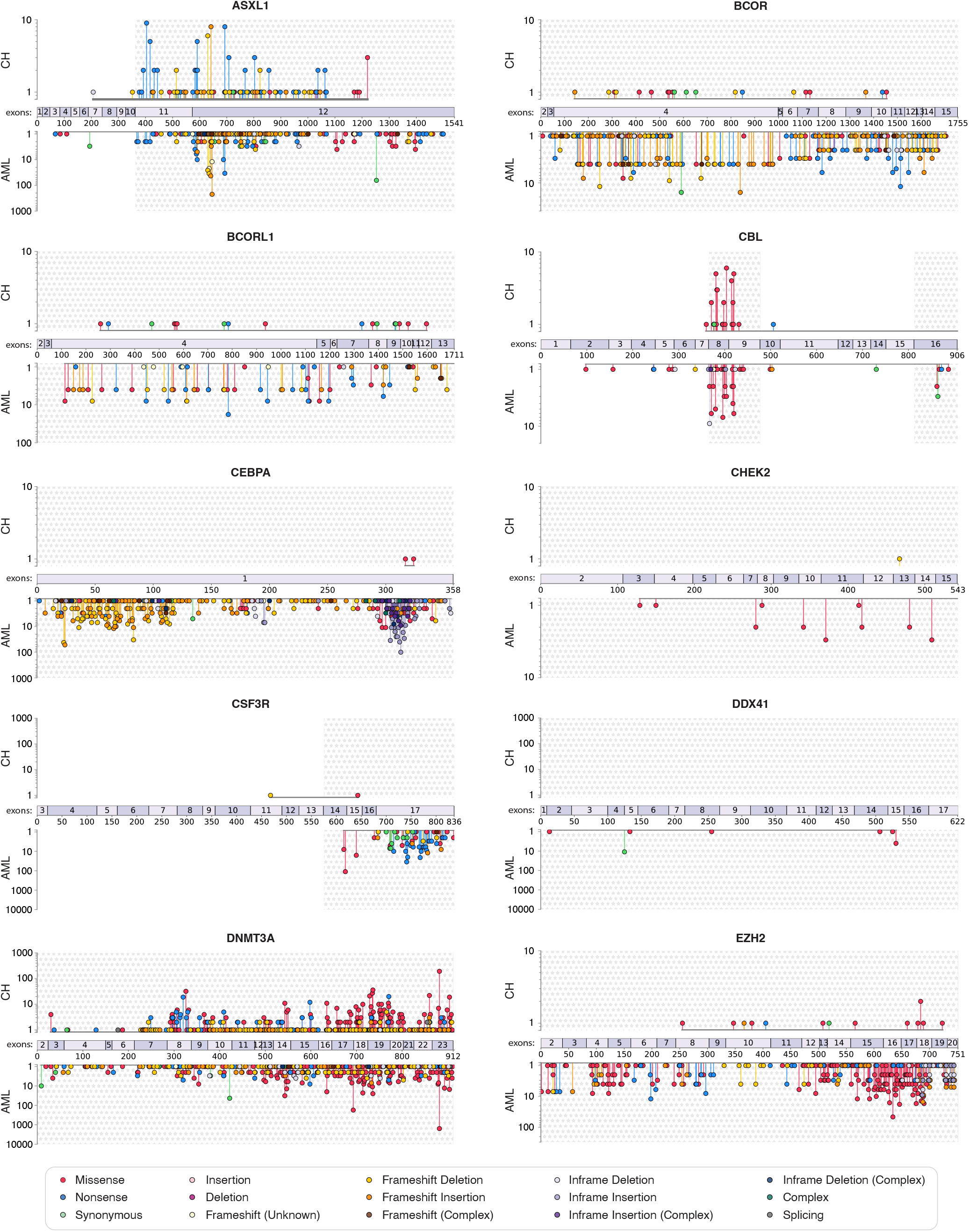
Gene regions tarted by TETRIS-seq SNV/ indel panel: part 1. The distribution of variants detected across 9 different clonal haematopoiesis studies ^*18,19,21,36,76*–*80*^ are shown in the upper part of the plot (‘CH’). The distribution of variants detected in individuals with AML in COSMIC ^*58*^ are shown in the lower part of the plot (‘AML’). Regions chosen for the TETRIS-seq custom SNV/ indel panel are highlighted with stars.

**Supplementary Fig. 2.**
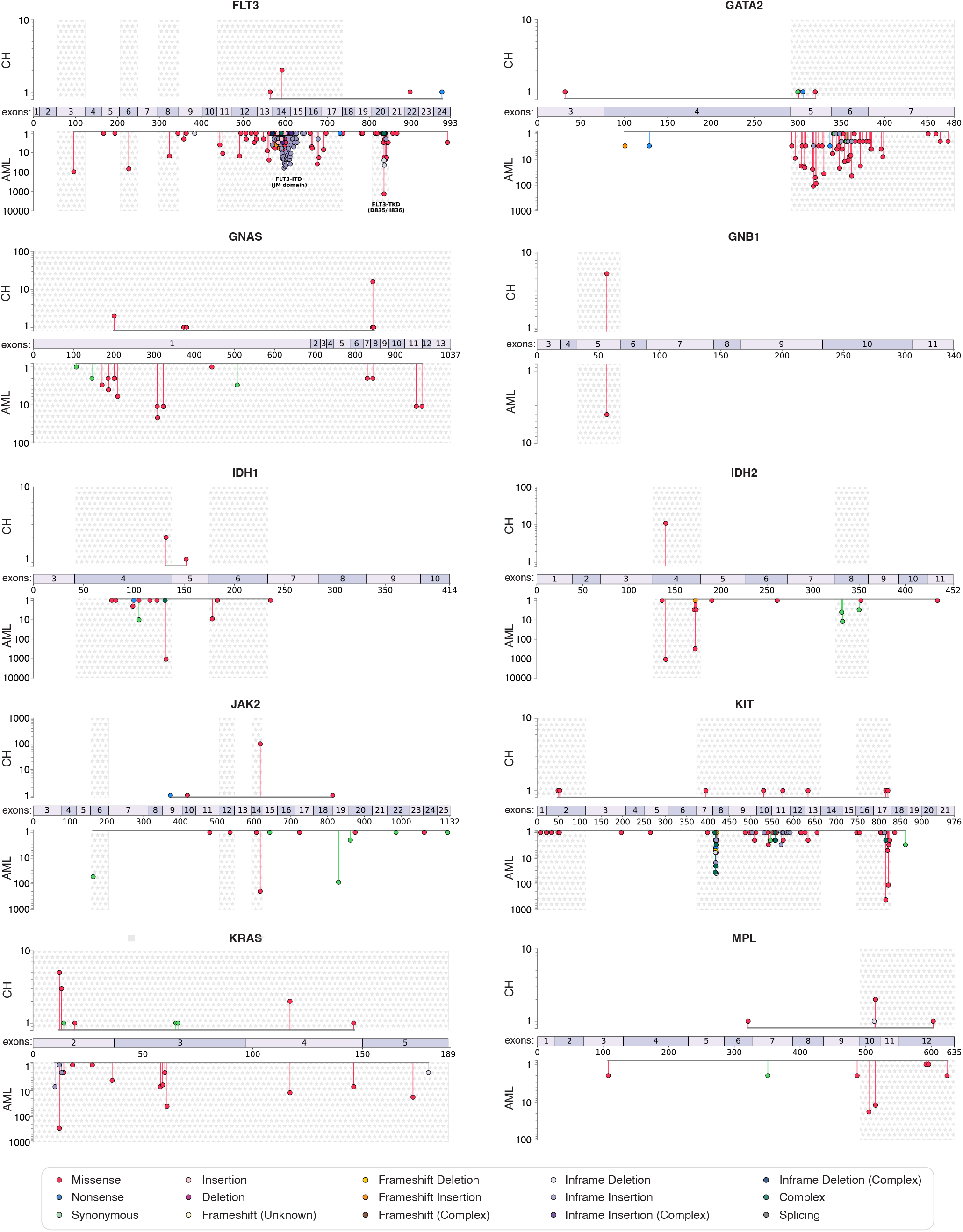
Gene regions tarted by TETRIS-seq SNV/ indel panel: part 2. The distribution of variants detected across 9 different clonal haematopoiesis studies ^*18,19,21,36,76*–*80*^ are shown in the upper part of the plot (‘CH’). The distribution of variants detected in individuals with AML in COSMIC ^*58*^ are shown in the lower part of the plot (‘AML’). Regions chosen for the TETRIS-seq custom SNV/ indel panel are highlighted with stars.

**Supplementary Fig. 3.**
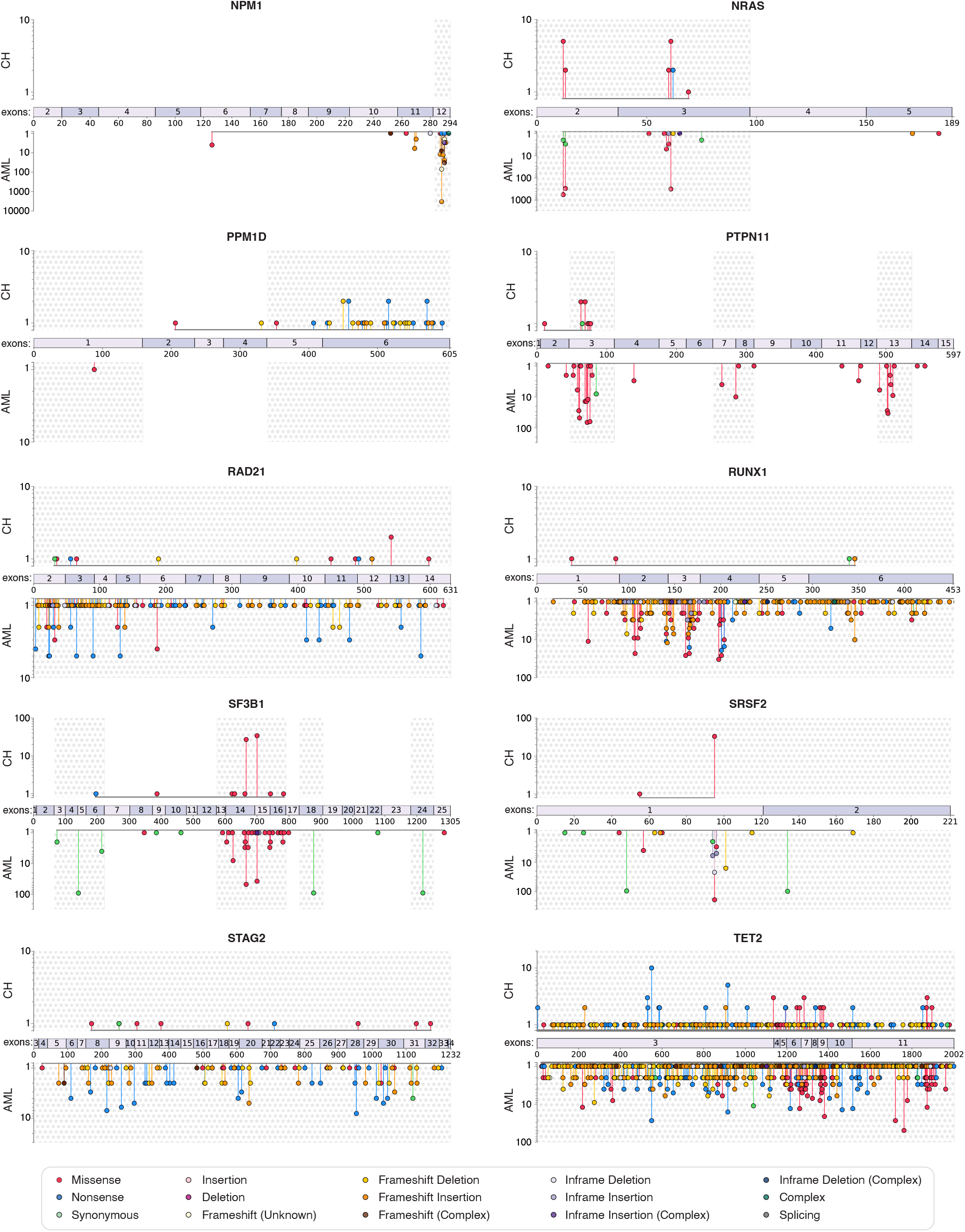
Gene regions tarted by TETRIS-seq SNV/ indel panel: part 3. The distribution of variants detected across 9 different clonal haematopoiesis studies ^*18,19,21,36,76*–*80*^ are shown in the upper part of the plot (‘CH’). The distribution of variants detected in individuals with AML in COSMIC ^*58*^ are shown in the lower part of the plot (‘AML’). Regions chosen for the TETRIS-seq custom SNV/ indel panel are highlighted with stars.

**Supplementary Fig. 4.**
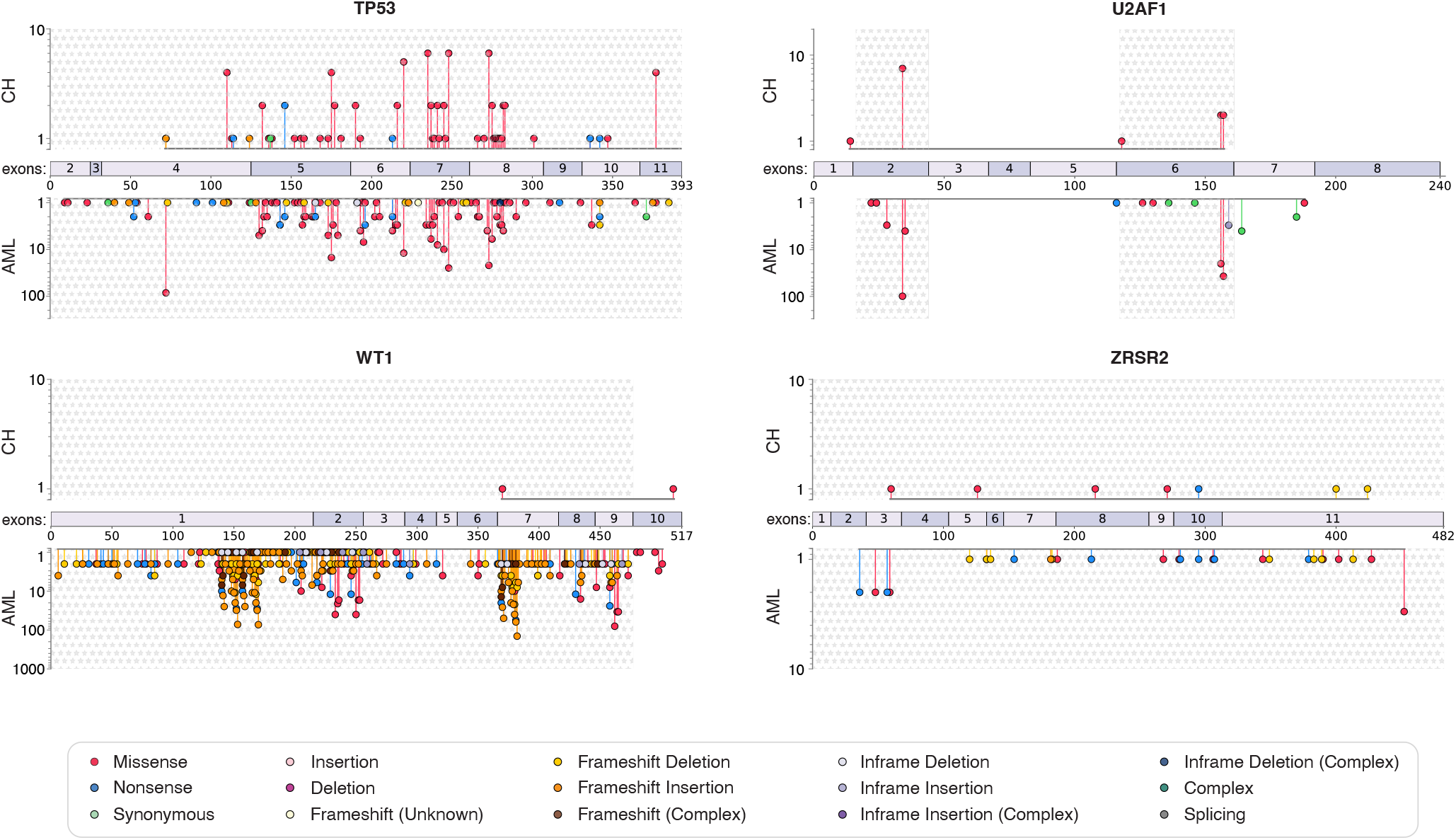
Gene regions tarted by TETRIS-seq SNV/ indel panel: part 4. The distribution of variants detected across 9 different clonal haematopoiesis studies ^*18,19,21,36,76*–*80*^ are shown in the upper part of the plot (‘CH’). The distribution of variants detected in individuals with AML in COSMIC ^*58*^ are shown in the lower part of the plot (‘AML’). Regions chosen for the TETRIS-seq custom SNV/ indel panel are highlighted with stars.

**Supplementary Fig. 5.**
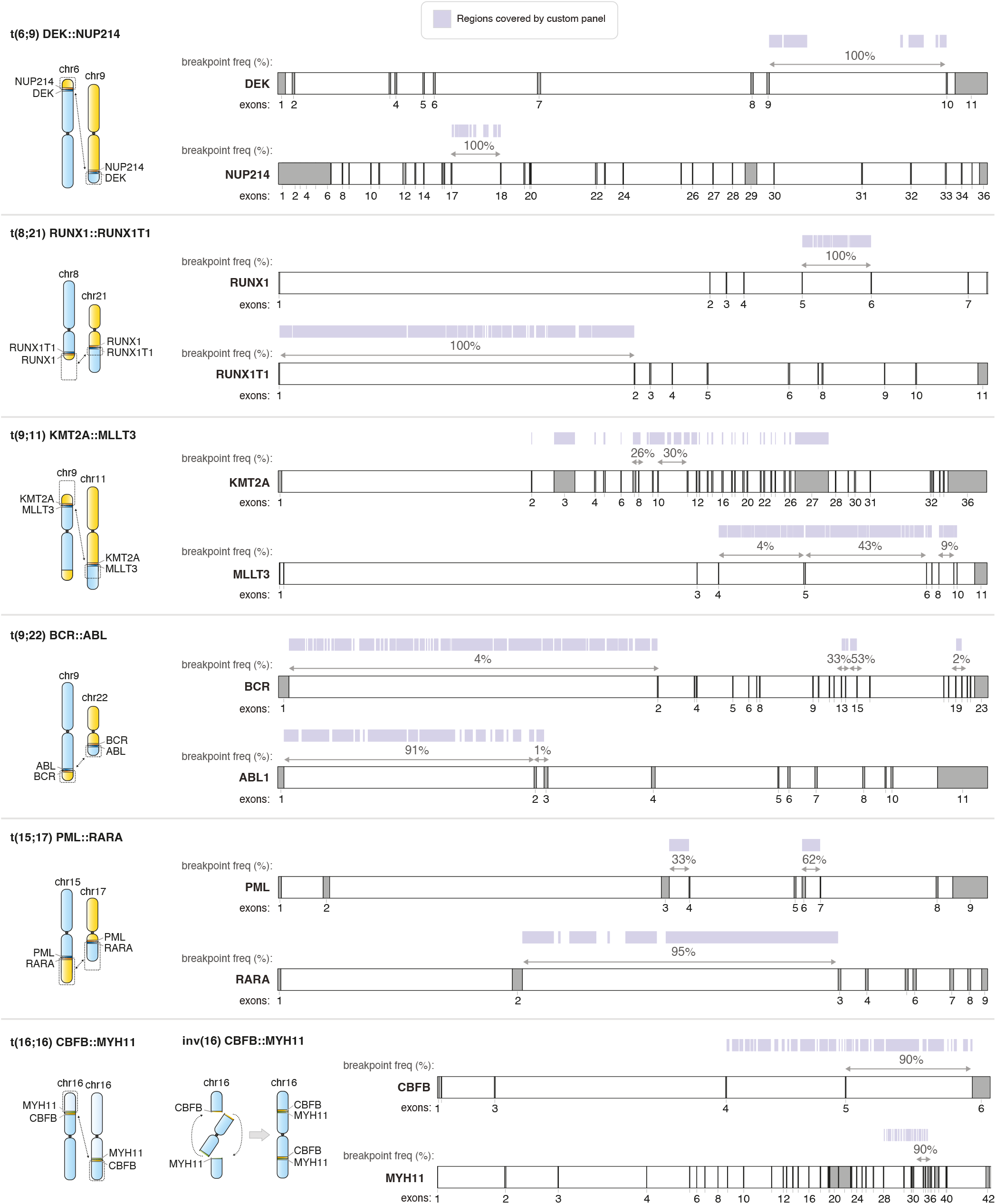
Custom panel coverage of chromosomal rearrangement breakpoint regions. The regions covered by the 120-nt oligonucleotide baits are shown in purple above the region. The frequency with which particular regions are the location of the breakpoint are written immediately above the region (indicated by arrows) ^*58*–*61*^.

#### C. Computational workflow for processing of error-corrected sequencing data

A custom computational workflow was written for the processing of the sequencing data, which consisted of four main steps: i) UMI extraction and initial alignment; ii) Single strand consensus sequence (SSCS) calling; iii) Duplex consensus sequence (DCS) calling; and iv) Putative variant detection.

##### i) UMI extraction and initial alignment

Sequenced reads were demultiplexed using their sample-specific dual indexes and the demultiplexed fastq files were converted to unmapped BAM files using Picard ^*66*^ *FastqToSam*. The inline UMIs were extracted from each read and stored in the ‘RX’ tag of the unmapped BAM file using fgbio *ExtractUmisFromBam*. Illumina adapter sequences were marked using Picard *MarkIlluminaAdapters* and the adapter-marked BAM was converted to a fastq using Picard *SamToFastq*, which also clipped the adapter sequences. The fastq was aligned to the The Broad Institute’s b37/hg19 reference genome ^*69*^ using BWA mem ^*68*^ to create a mapped BAM. Picard *MergeBamAlignment* was used to transfer the BAM tags from the unmapped adapter-marked BAM file to the mapped BAM file to ensure that the UMI information for each read was retained.

##### ii) Single strand consensus sequence (SSCS) calling

A custom Python script was written to generate single strand consensus sequences (SSCS) from the mapped BAM file in a stepwise manner. First, reads that had a mapping quality >20 and shared the same UMI tag, genomic coordinates and template length were grouped together to form ‘UMI families’. For the SNV/ indel panel, UMI families were discarded if they contained <3 reads. For the mCA/ chromosomal rearrangement panel, no minimum UMI family size was required. The reads in a UMI family were then compared at each sequence position, if their base quality score was >20, and a consensus nucleotide was called if there was at least 90% agreement between the reads. If there was <90% agreement, then an ‘N’ was called as the consensus nucleotide at that position. This meant, for UMI families containing <10 reads, there had to be 100% agreement between the reads at a position for the base to not be called as ‘N’. The resulting SSCS reads were written to an ‘unmapped’ SSCS BAM file. Reads that did not have a 99, 163, 147, 83 flag (i.e. were not ‘mapped in proper pairs’) were discarded, except for the generation of SSCS BAM files for FLT3-ITD calling and chromosomal translocation calling.

##### iii) Duplex consensus sequence (DCS) calling

A custom Python script was written to generate duplex consensus sequences (DCS) from the ‘unmapped’ SSCS BAM file in a stepwise manner. First, SSCS reads corresponding to a pair of the initial DNA strands were identified and grouped together. The UMI tag associated with each read consists of two 3-nucleotide sequences and the UMI tag of the read from its partner strand is a transposition of this, e.g. if a ‘read 1 forward’ sequence had the UMI tag ‘ATG-CAT’, it would be grouped with a ‘read 2 forward’ sequence that had the UMI tag ‘CAT-ATG’, the same genomic coordinates and the same template length. The paired SSCS sequences were then compared at each sequence position and a consensus nucleotide was called if the bases matched. If one or both of the bases had a quality score <20, or if the bases did not match, then an ‘N’ was called as the consensus nucleotide at that position. The resulting DCS reads were written to an ‘unmapped’ DCS BAM file with ‘forward strand’ DCSs becoming ‘read 1’ reads and ‘reverse strand’ DCSs becoming ‘read 2’ reads.

##### iV) Putative variant detection

For SSCS or DCS variant detection, the ‘unmapped’ SSCS or DCS BAM file was processed as follows: To identify any adapter sequences that may have been missed pre-consensus calling, Illumina adapter sequences were again marked using Picard *MarkIlluminaAdapters* and the adapter-marked SSCS or DCS BAM was converted to a fastq using Picard *SamToFastq*, which also clipped any remaining adapter sequences. The fastq was realigned to The Broad Institute’s b37/hg19 reference genome ^*69*^ using BWA-MEM ^*68*^ to create a mapped BAM file. Picard *MergeBamAlignment* was used to transfer the BAM tags, containing SSCS or DCS calling metrics, from the unmapped adapter-marked BAM file to the mapped BAM file. Overlapping reads and 3 nucleotides from the end of each read were clipped and then sequences were realigned with GATK’s Indel Realigner ^*69*^. The aligned sequences were processed with SAMtools ^*70*^ mpileup using the parameters -BOa -Q0 -d 1,000,000 to ensure all the pileups were returned without any filtering. A custom Python script was then written to generate a VCF file which contained SNV and indel information as well as the variant depth and total read depth at every position in the panel. All positions with a variant depth >0 were annotated with ANNOVAR ^*73*^. Indels were also called from the realigned BAM file using VarDictJava ^*71*^ and FLT3-ITD variants were called using Pindel ^*72*^.

#### D. Error-corrected sequencing metrics

In order to achieve as low a VAF limit of detection as possible, it is important to maximise the number of DCSs formed. Forming a DCS requires an SSCS to be generated from both DNA strands and so it is important that there are sufficient sequencing reads sharing the same UMI tag sequence to do this (i.e. sufficient UMI tag family size). Excluding the UMI family size peak at 1 (most likely due to sequencing errors in the UMI tags ^*87*^, an optimal peak family size of 6 is recommended to maximise the efficiency of duplex sequencing ^*87*^, striking a balance between optimal PCR input and sequencing depth/ cost. Nearly 75% of our samples had a peak UMI tag family size of 2 (Figure 6a), but our family size distributions were broad, with an average mean UMI tag family size of 8 and maximum UMI tag family size of ∼175 for both pre-AML and control samples (Supplementary Fig. 6b). This meant, even when we required a minimum UMI tag family size of 3 for an SSCS to be called, we still retained ∼92% of reads (Supplementary Fig. 6c), which represented ∼64% of UMI tag families (Supplementary Fig. 6d). Our mean SSCS:DCS ratio was 5-6 (Supplementary Fig. 6e). Across all samples, for the SNV/ indel panel, mean SSCS depth was ∼5500 and mean DCS depth was ∼1800. With ∼50ng DNA input (∼15,000 haploid genomes), this equates to an efficiency of ∼7.5% for SSCS and ∼6% for DCS.

**Supplementary Fig. 6.**
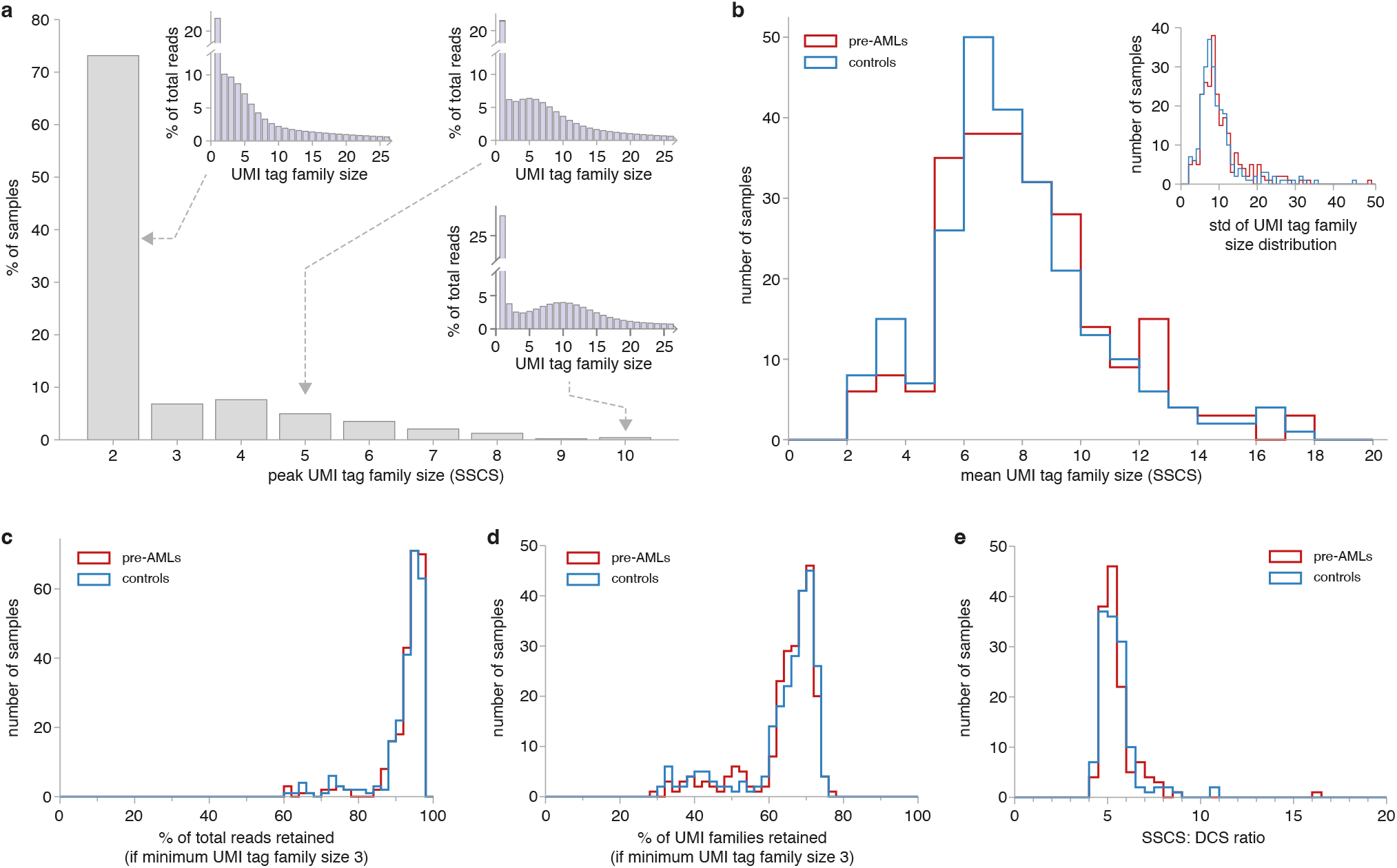
Error-corrected sequencing metrics (SNV/ indel panel). **a**. Distribution of peak UMI tag family sizes across all pre-AML and control samples. Insets show example UMI tag family size distributions with peaks of 2, 5 and 10. Example distributions have been capped at a UMI tag family size of 25. Average maximum UMI tag family size was 179 (range 60-1044) for pre-AML cases and 169 (range 57-875) for controls. **b**. Distribution of mean UMI tag family sizes across all pre-AML samples (red) and control samples (blue). Average mean UMI tag family size was 8 (range 2-17) for both pre-AML cases and controls. Inset shows the distribution of standard deviations (std) for the UMI tag family size distributions. **c**. Distribution of the proportion of total reads retained if those forming UMI tag family sizes of <3 are discarded. Mean proportion was 93% (range 61-98%) for pre-AML samples (red) and 92% (range 61-98%) for controls (blue). **d**. Distribution of the proportion of UMI tag families retained if those containing <3 reads are discarded. Mean proportion was 64% (range 29-77%) for pre-AML samples (red) and 64% (range 30-75%) for controls (blue). **e**. Distribution of ratio of SSCS to DCS reads across pre-AML samples and control samples. Mean SSCS:DCS ratio was 5.5 (range 4.3-16.2) for pre-AML cases (red) and 5.5 (range 4.4 to 10.9) for controls (blue).

##### How many variants do we expect to see?

In previous work we showed that the distribution of clone sizes is consistent with a simple branching process model of haematopoietic stem cell (HSC) divisions ^*38*^, such that the probability density as a function of VAF (*f*) is given by:

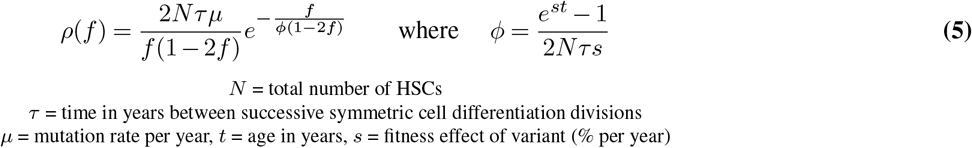

Single-cell derived sequencing work ^*37*^ provides us with HSC mutation rate estimates of ∼ 2.7 *×* 10^−9^ /bp /year, which, multiplied by the size of our SNV/ indel panel (∼ 58 kb), gives a mutation rate (*µ*) estimate of ∼ 1.6 *×* 10^−4^ per year. Our previous work has provided estimates for *Nτ* of ∼ 100, 000 as well as the distribution of fitness effects (*s*) of variants across 10 of the most commonly mutated clonal haematopoiesis genes ^*38*^. Using these parameters and integrating the expected density of clones (eq. 5) over the distribution of fitness effects and then over a range of VAFs (e.g. from 0.1% to 50%), allows us to estimate the number of variants we should expect to observe above a certain VAF in an individual of a particular age (Supplementary Fig. 7a). We can estimate that we should expect to see ∼ 8 variants between 0.1% and 50% VAF in a 65 year old individual, of which ∼ 5 will be at VAFs < 0.5%. This is much less than the 252 putative variants between 0.1% and 50% VAF (240 at VAFs <0.5%) that we observed in our test DNA sample from a 65 year old individual (Supplementary Fig. 7b). The theoretical error rate of duplex sequencing is often quoted as *<* 10^−9^ error per bp, which is simply the probability of two complementary errors occurring at the same nucleotide position on both DNA strands, either spontaneously or during the 1st PCR cycle ^*43,88*^. In reality, however, library preparation artefacts e.g. due to sonication, end-repair and mapping errors ^*44*^ have all been shown to slip through the duplex error correction and likely explains why we see far more ‘variants’ than we expect. Therefore, in order to reliably call ‘true variants’ <0.5% VAF we developed an *in silico* noise correction method for SNV calling, which involved developing a null model for the errors, so that inconsistencies from the model would be identified as ‘true variants’.

**Supplementary Fig. 7.**
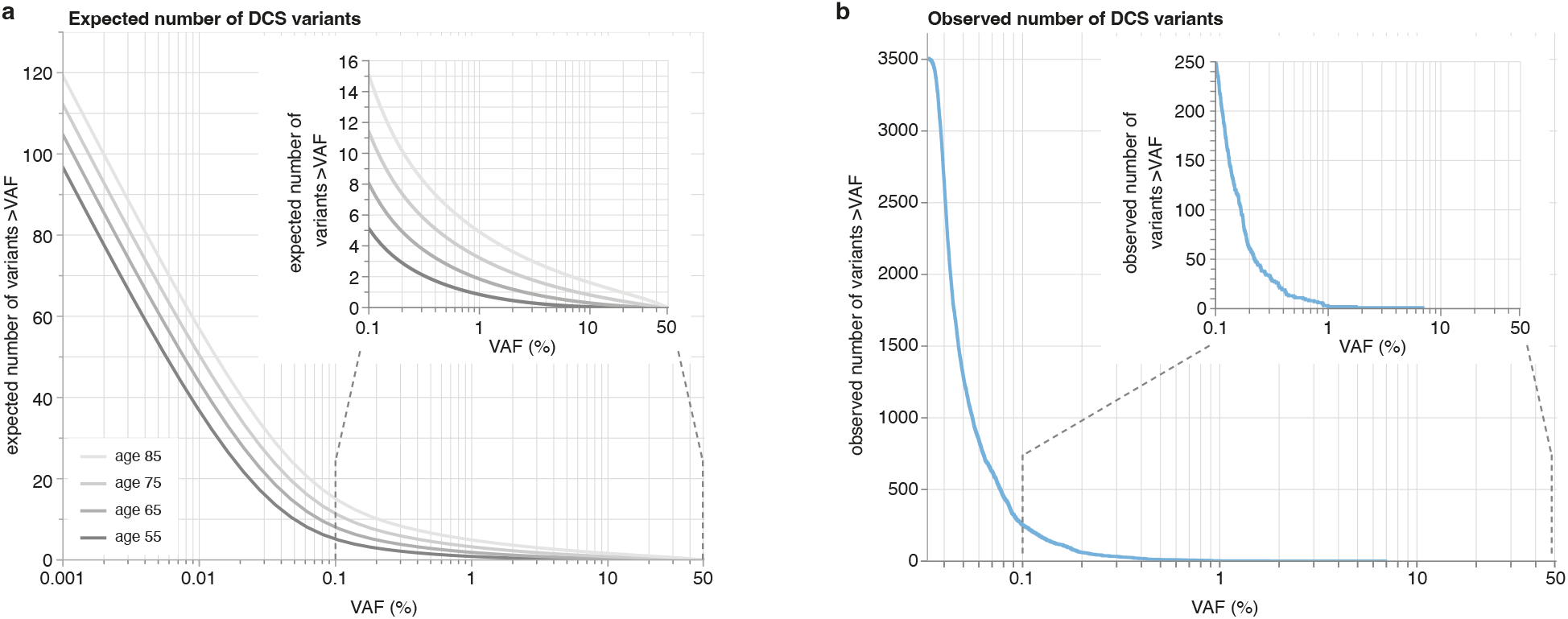
Expected and observed number of DCS variants. **a**. Expected number of variants detectable above a given VAF, using the custom SNV/ indel panel, for individuals aged 55, 65, 75 and 85 years. **b**. Number of DCS ‘variants’ observed above a given VAF, using the custom SNV/ indel panel, in DNA sample from a 65 year old individual.

#### E. Development and validation of an *in silico* noise correction model for SNV calling

To develop an *in silico* noise correction method for SNV calling we used sequencing data from 40 UKCTOCS samples (20 pre-AMLs and 20 controls), which had all been sequenced on the same Illumina NovoSeq S4 lane. The sequencing data was processed using the computational workflow described above and the DCS output files, containing variant depth and total depth at all positions in the SNV/indel panel, were used for error model development. The null hypothesis for our error-model was that each base-change at a position in the panel had a specific error rate (*ϵ*) and the number of variant reads (*k*) observed in a sample at that position would be consistent with beta-binomial sampling at that specific error rate. When considering many samples, the variation in total read depths between samples makes the distribution of variant reads difficult to interpret and so the distribution of ‘VAFs’ (variant reads/ total depth) across samples at a position base-change was visualised instead. We started by assuming that all ‘variants’ with the same position base-change could be real if their DCS ‘VAF’ was >1% and then calculated the position’s mean error rate, for that base change, from the ‘VAF’s of all the other samples. Looking at some example position-specific base-changes from across the panel (Supplementary Fig. 8a), we can see that whilst the majority appear consistent with a binomial position specific error model there are some that appear slightly over-dispersed, hence the reason for considering a beta-binomial position base-change specific error model, rather than a simple binomial model (Supplementary Fig. 8b). A beta-binomial position base-change specific error model allows for a wider variation in the sample errors at a given position base-change, controlled by a parameter *δ*:

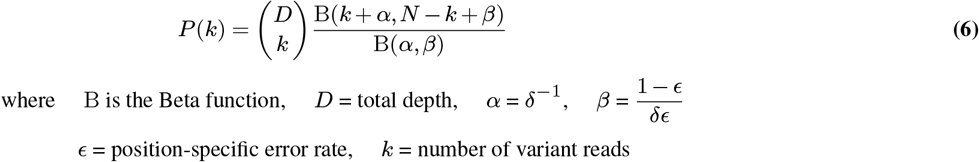

The binomial distribution naturally emerges from the beta-binomial distribution in the limit of *δϵ* ≪ 1 and so an advantage of a beta-binomial model is that it is also able to capture the behaviour of positions that are consistent with a simple binomial model.

**Supplementary Fig. 8.**
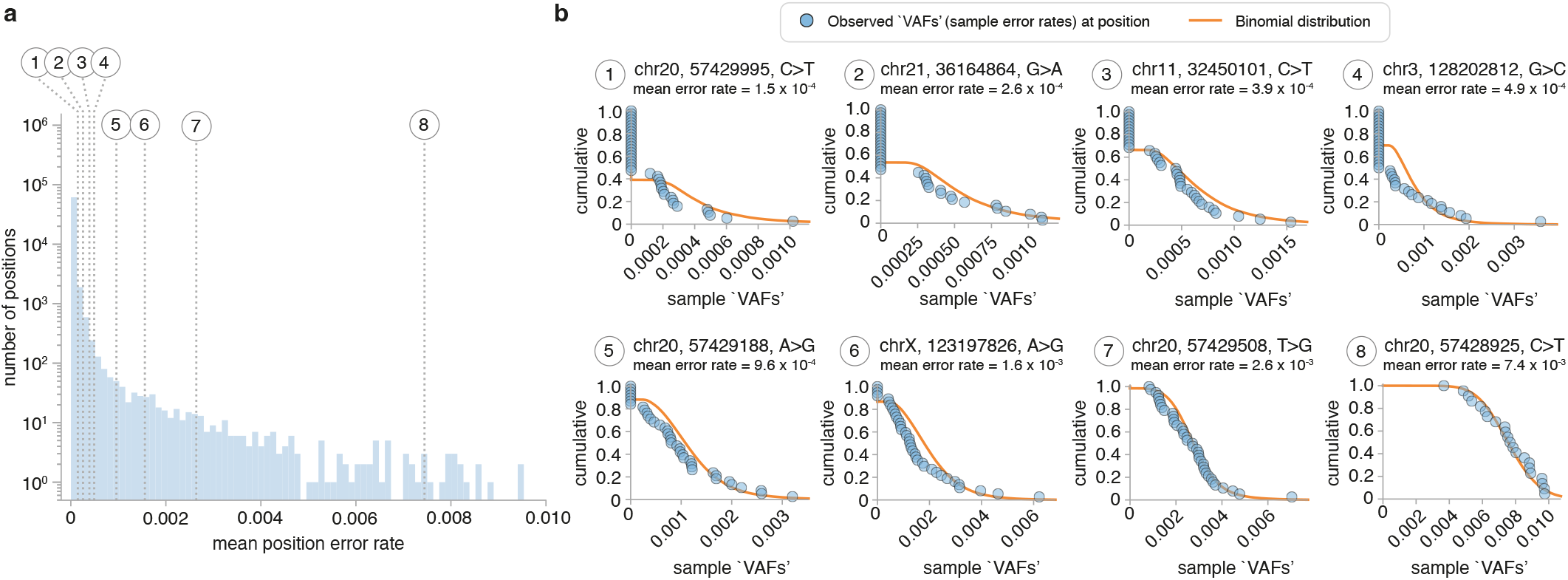
Distribution of sample DCS ‘VAFs’ (error-rates) at example positions. **a**. Distribution of mean position error rates across the SNV/indel panel. Samples with a position depth <500 were excluded from the error rate calculation and positions that had 0 variant reads across all samples at the position base-change are not shown in the distribution. Example positions across the distribution were chosen (numbered) to examine in more detail. **b**. 8 positions, that had at least 1 variant read detected in at least 16 samples with minimum sample depth of 500 and ‘VAF’ <1% were chosen from the distribution of mean position base-change error rates across the SNV/indel panel (in a.). Each sample’s ‘VAF’ (error rate) for the base-change at that position is shown in blue (cumulative distribution). If no variant reads were detected for that base-change in the sample then the error rate was recorded as 0. The orange line represents a binomial distribution, summed across all samples at the position where *n* = sample depth and *p* = mean position base-change error rate.

##### Inferring *ϵ* and *δ* parameters for identification of real variants

A custom Python script was written to infer the error rate (*c*) and dispersion (*δ*) values for each possible single nucleotide base-change at each position in the panel (hereafter referred to as simply ‘position’) and samples were called as ‘real’ variants if their variant read count was inconsistent with this distribution of errors. First, samples with a VAF >10% at the position were automatically called as real and excluded from the subsequent analysis. Then, the remaining samples at the position were used to fit either a beta-binomial distribution or a binomial distribution, depending on the number of samples at the position that had ≥ 1 variant reads:

If there were ≤ 3 samples remaining with ≥ 1 variant reads, there was insufficient data to fit a beta-binomial distribution and so the distribution of errors was assumed to be binomially distributed with *ϵ* = ∑ variant reads/ ∑ depth. The binomial *p*-values for all the samples with ≥1 variant reads were calculated and a sample was called as real if its *p*-value was less than a chosen *p*-value threshold.

If there were > 3 samples with ≥ 1 variants reads, a beta-binomial distribution was fitted to all the samples at the position. First, *ϵ* was estimated as ∑ variant reads/ ∑ depth across all the samples to be included in the fit and *δ* was estimated using the method-of-moments estimator (see Box 1). These *δ* and *ϵ* estimates were then used to initialise a maximum likelihood approach which re-inferred *δ* and *ϵ* by minimising the negative log likelihood of the model (see Box 2). If the method-of-moments estimate for *δ* was <1, suggesting the data was either undispersed or underdispersed relative to the binomial, then *δ* was initialised as 10^−4^ in the maximum-likelihood approach. A lower bound for *δ* was set to limit *δ/ ϵ* to >10^−8^ as the distribution had already become binomial at *δ* _*ϵ*_ <10^−6^ and numerical issues occurred below this level due to large *α* and *β* values. The beta-binomial *p*-values for all the samples with ≥1 variant reads were calculated and a sample was called as a real variant if it’s *p*-value was less than a chosen *p*-value threshold.

###### Box 1

**Method of moments estimator for *δ***

The method of moments estimator for *δ* can be calculated by noting the 1st and 2nd moments of the beta-binomial…

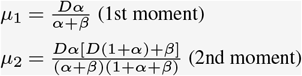

where *D* = mean total depth, 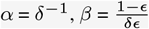 and ϵ = position error rate

Setting these raw moments equal to the 1st and 2nd raw sample moments…

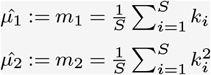

where *S* = number of samples and *k* = number of variant reads in a sample and solving for *α* and *β* we get…

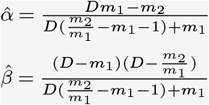

Given *δ* = 1/*α*, we can use this estimate for *α* to estimate *δ*.

###### Box 2

**Maximum likelihood approach for inferring *c* and *δ* parameters**

For each possible base change at a position, testing different position error rates (*ϵ*) and beta-binomial dispersion values (*δ*)…

1. For each sample, calculate the beta-binomial likelihood of measuring that number of variant reads for that base change at that position (*k*), given the sample depth (*D*), position specific base change error rate (*ϵ*) and dispersion value (*δ*):

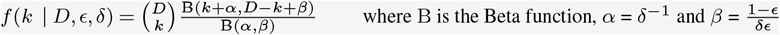
2. Calculate the likelihood of the model by multiplying across all the sample Beta-binomial likelihoods:

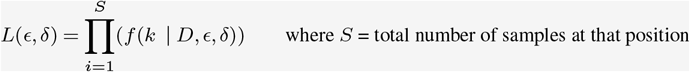 This creates very small numbers and so an alternative is to sum across all the sample log(Beta-binomial likelihoods):

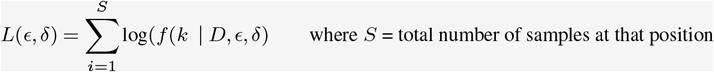
3. Find the values of *c* and *δ* that maximise the likelihood of the model (or minimize the negative likelihood of the model).

##### An iterative approach at potential ‘hotspot’ sites

When fitting a binomial or beta-binomial error distribution to a position only once, a problem arises if more than one sample at the position has a ‘real variant’. The problem is that the lower VAF variants will not be called as they will be fitted as ‘errors’ within a falsely over-dispersed beta-binomial distribution or a binomial distribution with a falsely elevated error rate. We therefore chose to use an iterative approach for positions observed in haematopoietic and lymphoid tissues in COSMIC v92 ^*58*^ (∼ 2% of sites across our custom panel), where we might expect to see more than one sample with a real variant in our pre-AML cohort.

- For positions with ≤ 3 samples with ≥ 1 variant reads, the iterative approach involved fitting a binomial distribution to all the samples, calling variants as real if their *p*-value was less than a chosen threshold, removing the real variants and then fitting the binomial again. This was then continued at the position until no further real variants were called.
- For positions with > 3 samples with ≥ 1 variants reads, the iterative approach was similar except at each iteration the sample with the highest VAF was excluded from the beta-binomial fit. This is because our prior probability of there being at least one real variant at these potential ‘hotspot’ positions is higher and if it were at high VAF it would result in a falsely over-dispersed beta-binomial distribution being fitted and real variants being missed. Beta-binomial *p*-values were then calculated for all samples, including the highest VAF sample, and variants were called as real if their *p*-value was less than a chosen threshold. This iterative approach (excluding the highest VAF sample from the beta-binomial fit each time) was then continued at the position until no further real variants were called.

##### An iterative approach for longitudinal samples

In the majority of cases, different timepoint samples from the same individual were sequenced on different flow-cells. When there was more than one timepoint sample from the same individual sequenced on the same flow-cell, an iterative approach was used for positions at which one of those samples had been called as a real variant. This is because, for these positions, there is a much higher chance that one of the other timepoints also contained the same variant, but at a different VAF. Without an iterative approach, it is likely the sample with the lower VAF would not be called as real. The iterative approach at the position was continued until no further samples from the individual with multi-timepoint samples were called as real.

##### Choosing an appropriate p-value threshold for calling real variants

To calculate an appropriate *p*-value threshold, we examined the false discovery rate (FDR) as a function of *p*-value threshold. Correctly estimating the FDR relies on the assumption that the underlying null model is correct, such that the number of false positives reliably increases as the *p*-value threshold increases. To check this, a custom Python script was written to simulate 40 samples across ∼575 positions, with the number of variant reads in each sample chosen from a beta-binomial distribution with a position specific *ϵ* and *δ* (i.e. all ‘variants’ were errors). Each position had a different *ϵ* and *δ*, covering regularly distributed combinations of *ϵ* between 10^−3^ and 10^−1^ and *δ* between 10^−2^ and 1. A beta-binomial distribution (or binomial if ≤ 3 samples with ≥ 1 variant reads) was fitted at each position (non-iteratively), *p*-values for each sample calculated and ‘real’ variants called if the sample’s beta-binomial (or binomial) *p*-value was less than the *p*-value being tested. The number of variants called across all positions was plotted as a function of *p*-value threshold (Supplementary Fig. 9a). This showed that the false positive rate was indeed consistent with what we would expect for most *p*-values, although there was a slight decrease at very low *p*-values. This may be a reflection of the fitted beta-binomial distribution slightly over-estimating the dispersion (*δ*) parameter and provides further support for our decision to exclude the highest VAF sample when fitting the beta-binomial distribution at ‘hotspot’ sites.

When we tested a range of different *p*-value thresholds on our real samples (∼40 UKCTOCS samples from 1 NovaSeq 6000 S4 lane), the false positive rate appeared consistently much lower than expected (Supplementary Fig. 9c). Initial analysis of the position error-rates had shown that most position error rates were very low, at < 10^−4^ (Supplementary Fig. 8), and so we repeated the simulated false positive analysis (as described above), but for positions where the error rate was much lower (*c*: 10^−4^ to 10^−3^, *δ*: 10^−3^ to 1). Interestingly, we saw a similar pattern to the real samples, where the false positive rate was consistently lower than expected (Supplementary Fig. 9b). We reasoned that this may be a consequence of the analysis being across only ∼40 samples, which means at very low *ϵ* there aren’t any samples that have ≥ 1 variant reads and so no false positive variants are called. Therefore, to chose a *p*-value threshold, we chose a *p*-value that gave an FDR of ∼ 5% (*p*-value: 6 *×* 10^−6^, with the understanding that the actual FDR will likely be even lower than this (i.e. FDR <5%) and that it is difficult to accurately estimate the FDR for this number of samples when the average error-rate is so low.

##### Position specific distributions of errors

Once the ‘real’ variants had been called at each position, a final beta-binomial or binomial distribution was fitted to the remaining samples at the position to ascertain each position’s distribution of errors. Across a pilot set of 20 pre-AML cases and 20 controls, real variants were called at 197 positions out of 174,366 positions (Supplementary Fig. 10). Of the positions where no variants were called, ∼60% contained only samples with 0 variant reads (Supplementary Fig. 10: A positions) and ∼30% of positions were fitted using a binomial distribution because there were ≤3 samples with ≥1 variant reads (Supplementary Fig. 10: B, C positions). The fitted beta-binomial distribution was consistent with a binomial distribution in ∼5% of positions (Supplementary Fig. 10: D-F positions) and a beta-binomial distribution in ∼ 3% of positions (Supplementary Fig. 10: G-L positions). Of the positions where 1 or more real variants were called (197 positions), ∼40% had only samples with 0 variant reads remaining and ∼ 35% were fitted using a binomial distribution because there were ≤3 samples remaining with ≥ 1 variant reads after the real variants were called. The fitted beta-binomial distribution was consistent with a binomial distribution in ∼11% of positions (Supplementary Fig. 10: P, Q positions) and was consistent with a beta-binomial distribution in ∼10% of positions (Supplementary Fig. 10: R-T positions). Overall, across all positions, 61% contained only samples with 0 variant reads, 36% were consistent with a binomial distribution of errors, and 3% were consistent with a beta-binomial distribution of errors. Looking at example positions from across the range of error distributions, we can see that the model appears to be performing well, appropriately fitting a beta-binomial distribution to positions whose distribution of errors appears over-dispersed relative to a binomial (Supplementary Fig. 10 subplots).

**Supplementary Fig. 9.**
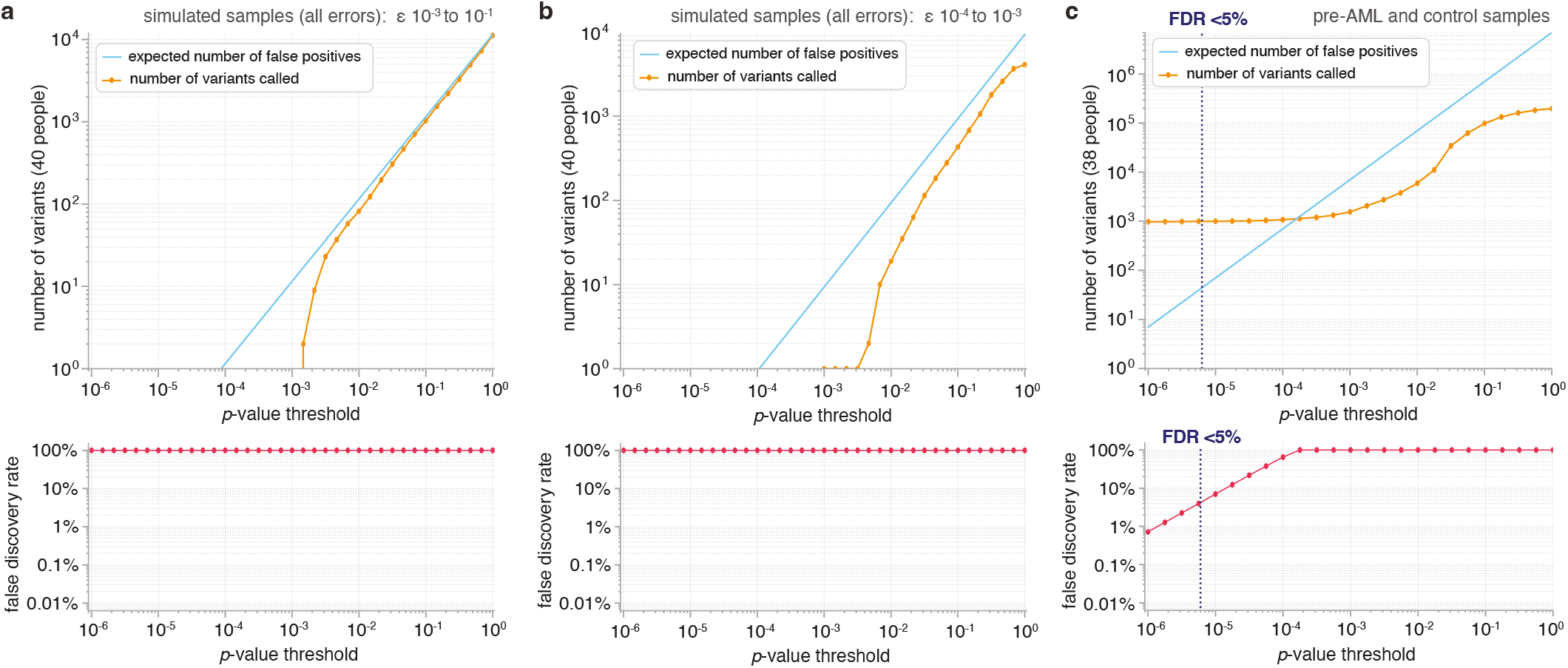
Choosing a *p*-value threshold for calling real variants. The point at which the read data (orange line) starts to deviate from the blue line, represents the *p*-value threshold where there are equal numbers of false positives and true positives. If we lower the *p*-value threshold further, the proportion of true positives increases, but we also increase the false negative rate.

**Supplementary Fig. 10.**
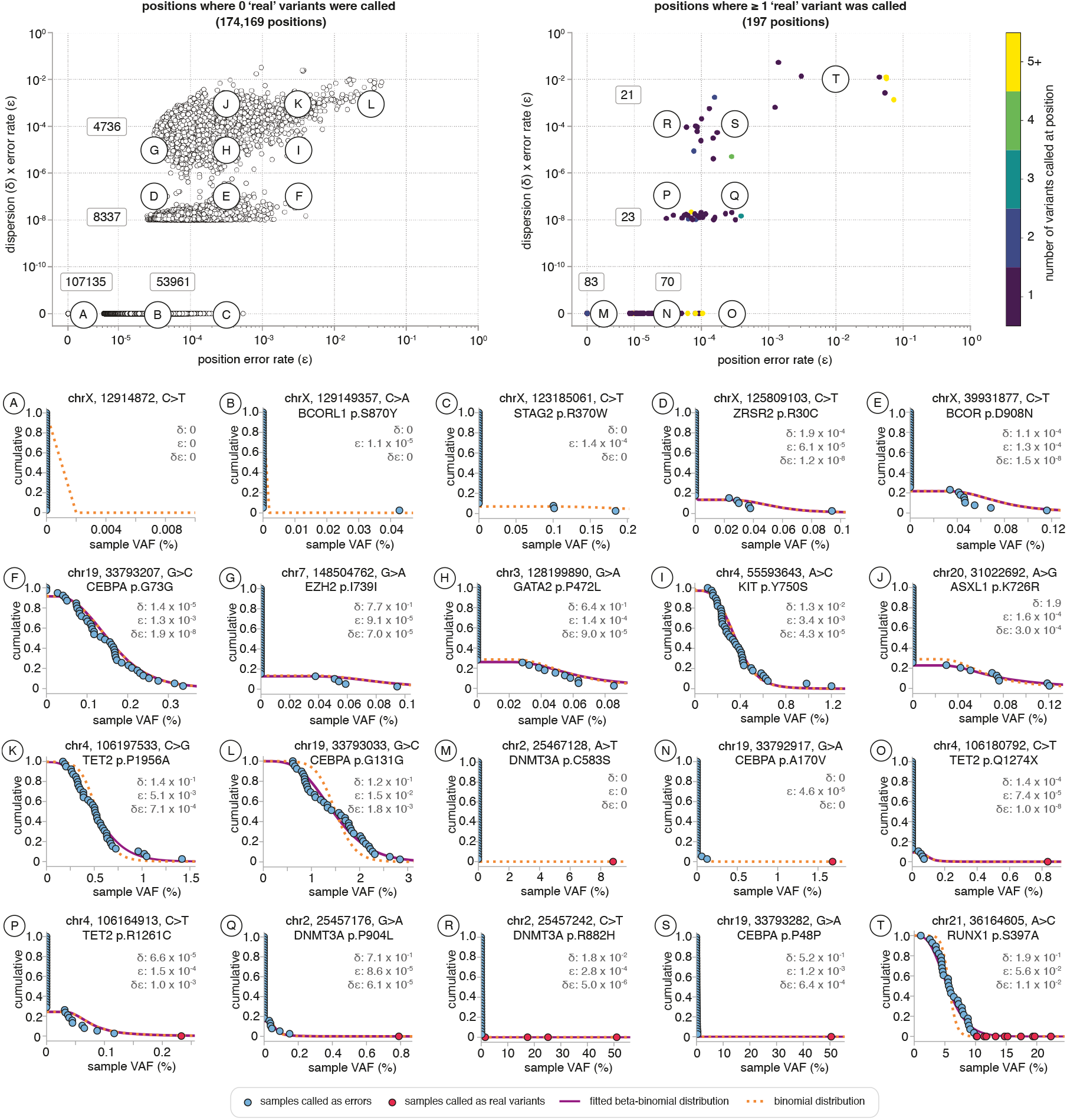
Position specific error distributions across 20 pre-AML and 20 control UKCTOCs samples. The relationship between the final position error rate (*ϵ*) and *δ* _*ϵ*_ is shown for positions where no variants were called as real (top left plot) and positions where 1 or more variants were called as real (top right plot). The number of positions in each area is shown on the plot. Example positions from different *δ* and *δ* _*ϵ*_ regions on the plot (indicated by letters A to T) are shown below. The final fitted error distribution is plotted as a purple solid line (beta-binomial) or orange dashed line (binomial). The binomial distribution is also shown for beta-binomially distributed positions for comparison. Samples called as real variants are shown in red. Samples called as errors are shown in blue.

**Supplementary Fig. 11.**
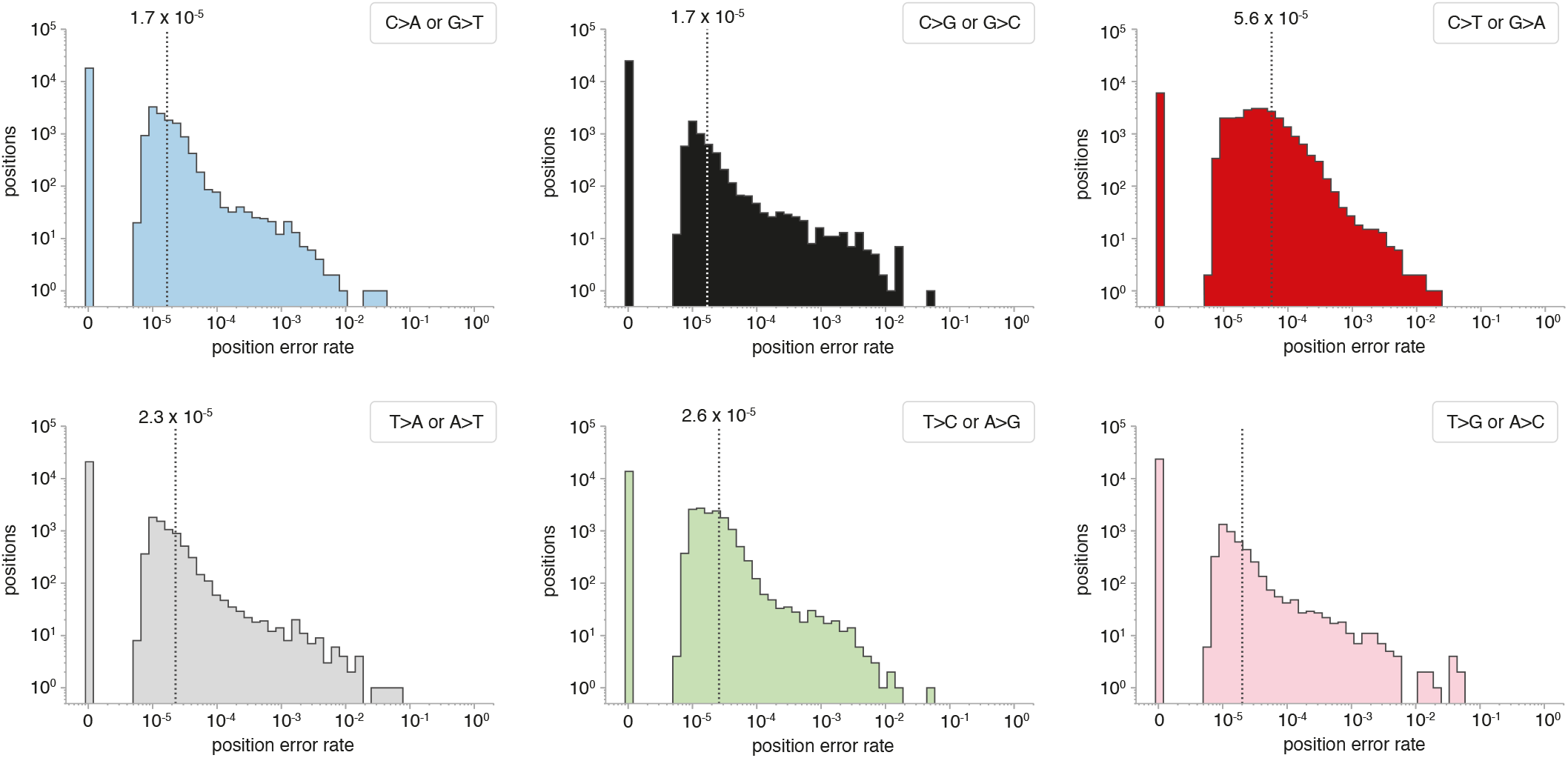
Distribution of final position-specific error rates, grouped by base change. The overall base-change error-rate is shown (dashed line) and was calculated as Σvariant reads/Σ depth for the samples that were not called as ‘real variants’ at all the positions with the particular base change.

##### Overall panel error rate

The overall error rate across the panel can be calculated as: Σ variant reads/Σ depth for the samples that were not called as real variants, and was 2.6 *×* 10^−5^ across an initial pilot of 20 pre-AML cases and 20 controls (one sequencing lane). This means, on average, we would be unable to call variants at frequencies lower than this (i.e. VAF detection limit > 0.026%). Categorising the positions by their base change, we can see that C>T and G>A variants tend to have an error rate ∼5x higher than other base changes, with error rates as high as ∼10^−2^ (Supplementary Fig. 11). The distribution of position error rates is quite broad, with some positions having error-rates as high as nearly 10^−1^, meaning there are some positions where we would be unlikely to call real variants unless they had a VAF > 10%.

#### F. Post-processing of variant calls

The error model was applied to all the samples sequenced on the same sequencing lane (∼ 40 samples) and then additional post-processing filters were applied:

- Variants called as ‘errors’ were ‘rescued’ and called as ‘real’ if their *p*-value was <0.1 and they had been called as ‘real’ (using the position-specific beta-binomial error model) with a *p*-value *<* 10^−10^ at any of the other timepoints in that individual.
  - Variants were filtered as ‘likely germline’ if any of the following applied:
  - Frequency in ExAC >0.001
  - The variant has an RSID
  - The variant had a mean VAF >40% (across all timepoints) in >5% controls and was observed in COSMIC v92 ^*58*^ ≤5 times.
  - The variant’s VAF at every timepoint was consistent with sequencing error of a heterozygous or homozygous germline variant.
- Variants were excluded if:
  - Seen in ≥5% controls, unless the variant had been observed in haematopoietic/ lymphoid tissues in COSMIC v92 ^*58*^ ≥1 time.
  - Detected in only 1 variant read (unless FLT3-ITD)
  - Mean DCS depth across the panel for that sample was <100
  - Total read depth at the position was <500 or >2 standard deviations below the mean panel read depth for that sample.
  - Synonymous variant
  - Mean position in read <8
  - Strand bias phred score >60
  - None of the timepoints for the variant had a *p*-value <0.1

Having longitudinal samples provided us with additional power for identifying errors, as we were able to use information from variants that were called, or not called, at multiple timepoints. There were variants that were not detected at some timepoints, but were detected at the timepoints before and after. The expected growth rate was calculated from the VAFs at the two detected timepoints (VAF_1_ at *t*_1_ and VAF_3_ at *t*_3_) to determine how many reads (*k*) the variant should have been expected to be seen in at the missing timepoint (*t*_2_), taking in to account the read depth of the missing sample:

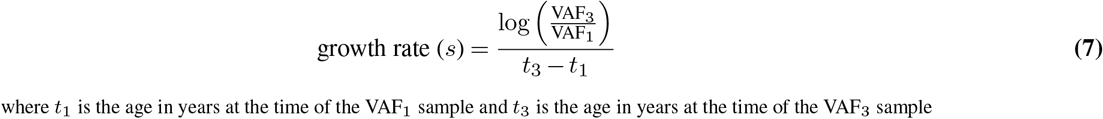

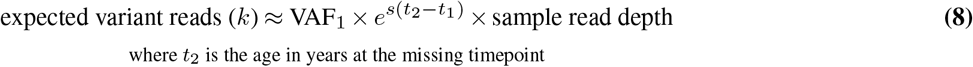

The chance of not detecting the variant at *t*_2_ was then calculated:

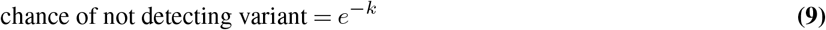

If the chance of not detecting the variant was <10%, then the variants from the preceding timepoints were excluded, as it was deemed likely to also be an error. If a detected variant was missing from more than 1/3 of the timepoints, the chance of not detecting the variant was calculated at all the missing timepoints and if the combined chance of missing all of them was <10%, then the variant was excluded from all timepoints.

Indels were excluded if they were detected in <4 DCS reads, unless they were a known NPM1 exon 12 frameshift hotspot mutation. Indels were also excluded if they were seen in ≥10% of people, unless the variant had been observed in haematopoietic and lymphoid tissues in COSMIC v92 ^*58*^. Using information from multiple timepoints, indels were excluded if they were not detected at >4% VAF at ≥1 timepoint and indels that were detected at some timepoints, but not others, were filtered using the same method as for SNVs (eqs. 7-9).

#### G. Detecting mosaic chromosomal alterations (mCAs)

A custom Python script was written to enable detection of mCAs from the SSCS reads using the SNP ‘B-allele frequencies’ (BAF) and read depths (log_2_R ratios) in the mCA/ chromosomal rearrangement panel.

##### Calculating log_2_ R ratios (LRR) and B-allele frequencies (BAF)

For each sample, the average read depth across each 120 bp targeted region (SNP region) was calculated. The average read depth of each SNP region was then normalised to the average read depth across all targeted regions in the sample, to create ‘sample normalised read depths’ for each SNP region. To account for inter-region variation in read depth coverage, due to variability in capture efficiency, the ∼20 control samples from the same sequencing lane were used as a ‘panel of normals’ to calculate the average normalised read depth for each SNP region. The mean coefficient of variation (CV) in read depths was calculated across the ‘panel of normals’, for each chromosome, and controls were excluded from the ‘panel of normals’ if their read depth CV for ≥ 2 chromosomes was more than 1.5 standard deviations above the mean CV. This helped to prevent any samples that had a significant copy number alteration (i.e. gain or loss event) being included in the ‘panel of normals’. Log_2_R ratios (LRR) were then calculated for each SNP region by comparing the region’s ‘sample normalised read depth’ to the average normalised read depth for that region from the ‘panel of normals’. The VAF of all targeted SNPs, as well as any SNPs >1% MAF in 1000 genomes ^*83*^ that were also covered by the panel, were calculated (from read depth/total depth) and were plotted across the chromosomes to visualise the B-allele frequency (BAF).

##### Identifying mCAs as either gain, loss or CN-LOH events

Once BAFs and log_2_R ratios (LRR) had been calculated, it was possible to identify regions in which either copy number alterations (i.e loss or gain events) or copy-neutral loss of heterozygosity (CN-LOH) events had occurred. Loss and gain events result in deviations in both LRR and BAF whereas CN-LOH events result in BAF deviations without a change in LRR (because there is no change in the amount of genetic material) (Supplementary Fig. 12).

**Supplementary Fig. 12.**
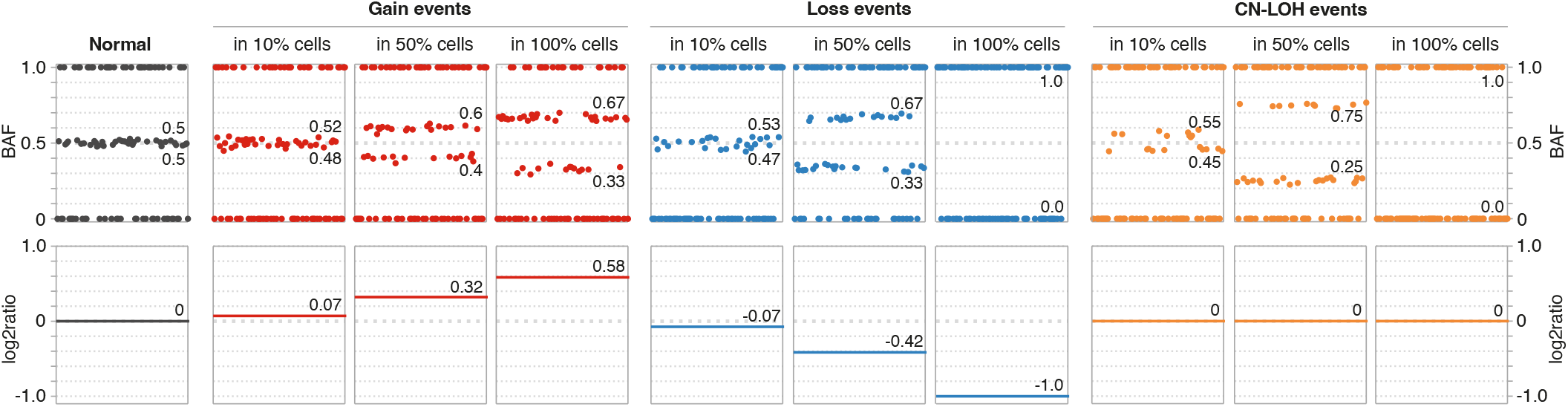
LRR and BAF deviations for mCA detection. Normal regions (black plot) contain SNPs with B-allele frequencies (BAFs) of 0 (homozygous AA alleles), 0.5 (heterozygous AB alleles) and 1.0 (homozygous BB alleles) and no change in read depth (log_2_ ratio of 0). Regions with gain events (red plots) show deviations in BAF, up to a maximum of *±*0.17 if a duplication event is in 100% of cells. Gain events result in an increase in read depth (log_2_ ratio > 0), due to the extra genetic material. Regions with loss events (blue plots) or CN-LOH events (orange plots) also show deviations in BAF and can both have full loss of heterozygosity if the loss or CN-LOH affects 100% of cells. Loss and CN-LOH events can be distinguished by the read depths, which are decreased in loss events (log_2_ ratio < 0) due to loss of genetic material, but are unchanged in CN-LOH events (log_2_ ratio = 0). Data generated for schematic using simulated samples (100 SNPs per region with mean read depth across ‘panel’ of 1000 reads).

##### Determining length and cell fraction of mCAs

Because BAF deviations could occur due to inadvertent somatic variants at common SNP sites or variable bait capture, we required BAF deviations (or lack of heterozygosity) in ≥5 consecutive SNPs for an mCA to be called. With SNPs spaced ∼280kb apart, this meant the smallest mCA we could expect to detect would be ∼1.5 MB. The ‘start’ of the mCA was taken as the coordinate of the first BAF-deviated SNP and the ‘end’ coordinate was taken as the coordinate of the last BAF-deviated SNP in the affected region. The proportion of cells (‘cell fraction’) harbouring the mCA was calculated from the heterozygous BAFs as detailed in Supplementary Table S1. If an mCA was detected at 100% cell fraction in both the latest and earliest timepoint sample from an individual, then it was deemed likely to be germline.

**Table S1.** Expected LRR, heterozygous BAFs and cell fractions for autosomal somatic mCAs. mCA cell fraction (*p*) can be calculated directly from the heterozygous BAFs (*µ*_1_ and *µ*_2_). Adapted from Jacobs et al. ^*89*^

| Somatic mCA (autosomal) | LRR | Heterozygous BAFs ($\mu_1, \mu_2$) | mCA cell fraction ($p$) |
| --- | --- | --- | --- |
| Gain | $\log_2\left(\frac{2+p}{2}\right)$ | $0.5 \pm \frac{p}{2(2+p)}$ | $\frac{2(\mu_2 - \mu_1)}{1 - (\mu_2 - \mu_1)}$ |
| Loss | $\log_2\left(\frac{2-p}{2}\right)$ | $0.5 \pm \frac{p}{2(2-p)}$ | $\frac{2(\mu_2 - \mu_1)}{1 + \mu_2 - \mu_1}$ |
| CN-LOH | 0 | $0.5 \pm \frac{p}{2}$ | $\mu_2 - \mu_1$ |

##### Using phasing information to call mCAs at low cell fractions

Because mCA calling relies on the ability to detect deviations in heterozygous BAFs, the noisier the BAF measurements (e.g. due to low sequencing read depth), the harder it will be to detect subtle BAF deviations associated with low cell fraction mCAs (e.g. gain event in 10% cells in Supplementary Fig. 12). Haplotype phasing, which involves identifying SNPs which lie on the same chromosome and therefore have the same direction of BAF deviation, can be used to improve the sensitivity of low cell fraction mCA calling. This approach has been used to call mCAs at cell fractions as low as 1% in UK Biobank participants ^*90,91*^ by utilising long-range phase information generated from long identical-by-descent (IBD) tracts shared among distantly related individuals ^*92*^. This information is unfortunately not available for the UKCTOCs participants. However, one of the benefits afforded by having longitudinal samples is the ability to use large BAF deviations from higher-cell fraction mCAs, detected at timepoints closer to AML diagnosis, to identify which SNPs lie on the same chromosome in the affected region of interest. This phasing information can then be applied to the same individual’s samples from earlier timepoints, allowing a much higher sensitivity for detection of the mCA when it is at lower cell fraction (Supplementary Fig. 13). Using this approach we were able to call mCAs at cell fractions as low as 0.1%.

#### H. Detecting *KMT2A* partial tandem duplications (*KMT2A*-PTD)

Partial tandem duplications in *KMT2A* (*KMT2A*-PTD) most commonly involve exons 2 or 3 and span through exon 9 to 11 ^*64*^. Exon 27, which is the largest exon in *KMT2A*, is characteristically not involved. Therefore to detect *KMT2A*-PTD events a custom Python script was written (using pysam ^*75*^ pileup) to calculate the mean read depth across each targeted KMT2A exon. These mean depths were then normalised to the mean read depth across *KMT2A* exon 27 and the exon 3:27 ratio (*R*) calculated.

**Supplementary Fig. 13.**
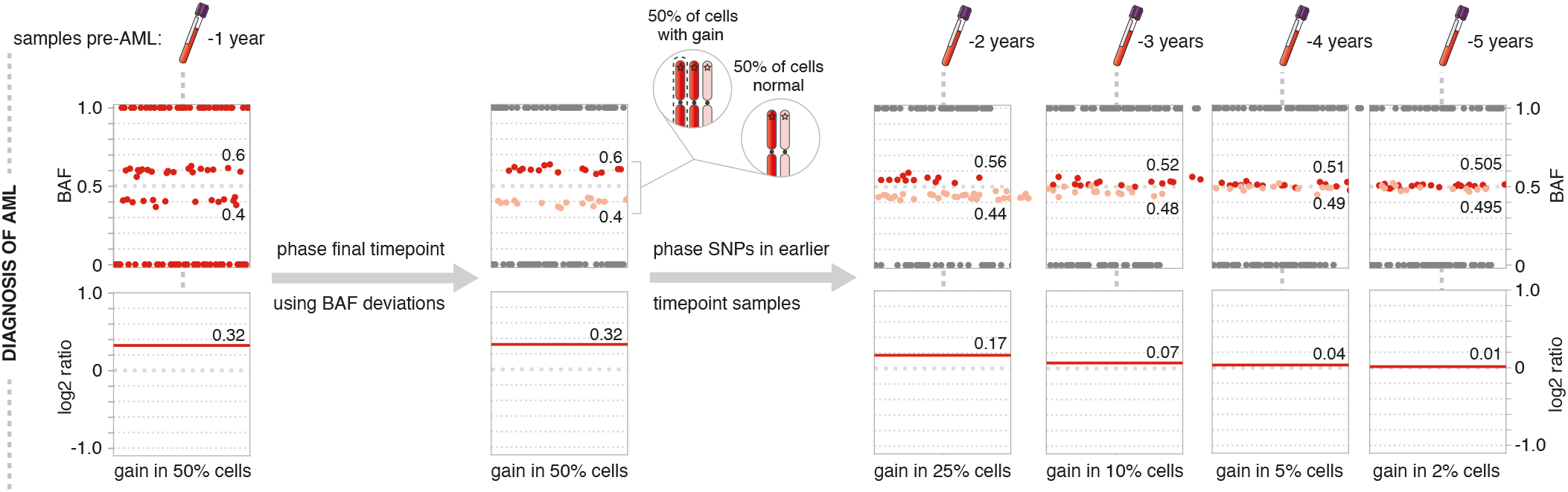
Phasing SNPs for detection of low cell fraction mCAs. mCAs detected at high enough cell fraction to allow differentiation of the heterozygous SNPs that have deviated above and below 0.5 BAF enable SNPs to be ‘phased’ (e.g. in the final timepoint ‘sample’ pre-AML diagnosis shown on the left). This phasing information can then be applied to earlier timepoint samples (shown on the right) and provides higher sensitivity for detection of smaller BAF deviations and therefore higher sensitivity for detection of mCAs at lower cell fraction. Data generated for schematic using a simulated sample (100 SNPs per region with mean read depth across ‘panel’ of 1000 reads).

The fraction of cells harbouring the *KMT2A*-PTD can then be calculated as:

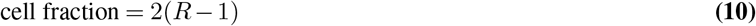

A *KMT2A*-PTD affecting one *KMT2A* allele in 100% of cells should be easy to detect, with a mean read depth ratio (*R*) of 1.5 (Supplementary Fig. 14a). At lower cell fractions, however, the ratio becomes quite small and may become harder to detect (Supplementary Fig. 14b), particularly if the read depths within each exon are highly variable. This is a limitation of our *KMT2A*-PTD detection method and means we may not detect *KMT2A*-PTD events pre-AML diagnosis if they are at low cell fraction. *KMT2A*-PTD are effectively small gain events and so SNP BAFs could be used to improve sensitivity, akin to their use in mCA detection. There are, however, only 10 SNPs at >1% MAF in 1000 genomes ^*83*^ across our targeted *KMT2A* regions and none of these are in exon 3.

**Supplementary Fig. 14.**
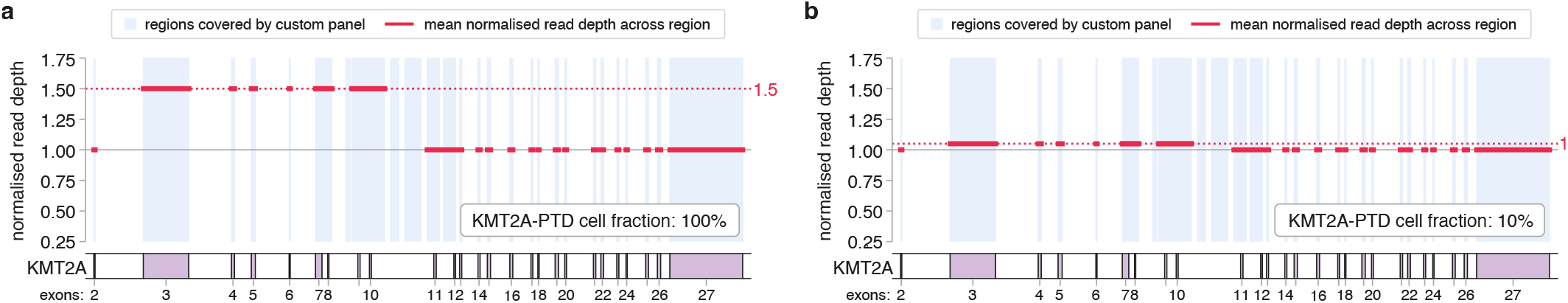
Schematic showing the effect of *KMT2A*-PTD on exon 3: exon 27 read depth ratios. **a**. *KMT2A*-PTD involving exons 3-10 in 100% of cells results in an exon 3: exon 27 read depth ratio of 1.5. **b**. *KMT2A*-PTD involving exons 3-10 in 10% of cells results in an exon 3: exon 27 read depth ratio of 1.05.

### 2. Reconstruction of clonal evolutionary histories in pre-AML and control samples

**Supplementary Fig. 15.**
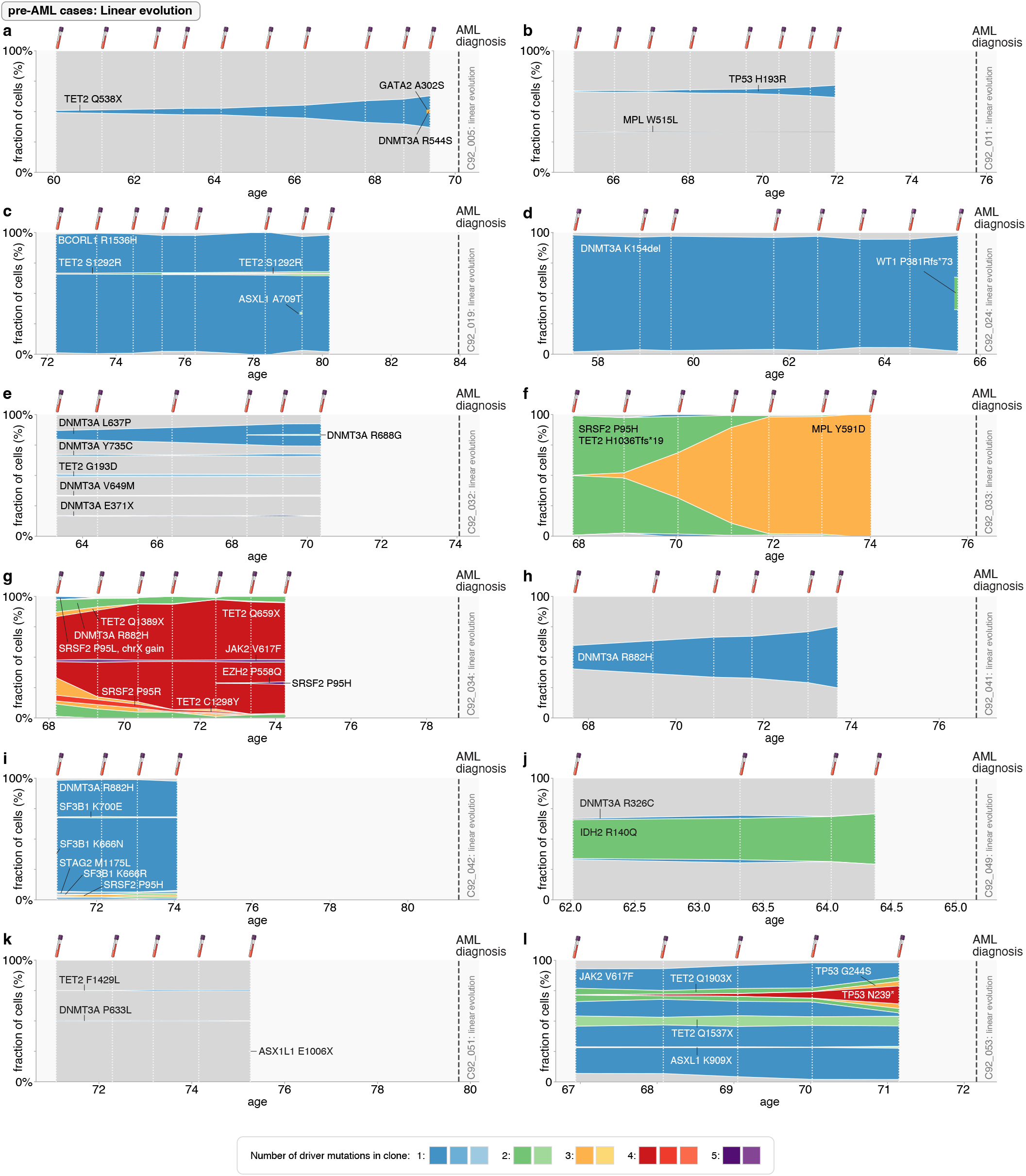
Reconstruction of clonal evolutionary histories: pre-AML cases with linear evolution part 1. Muller plots exhibiting clonal dynamics in pre-AML cases showing “linear evolution”. Colouring denotes the number of observed driver mutations in each clone (see legend). White vertical dashed lines indicate timing of UKCTOCS blood samples. Thick grey vertical dashed line indicates time of AML diagnosis.

**Supplementary Fig. 16.**
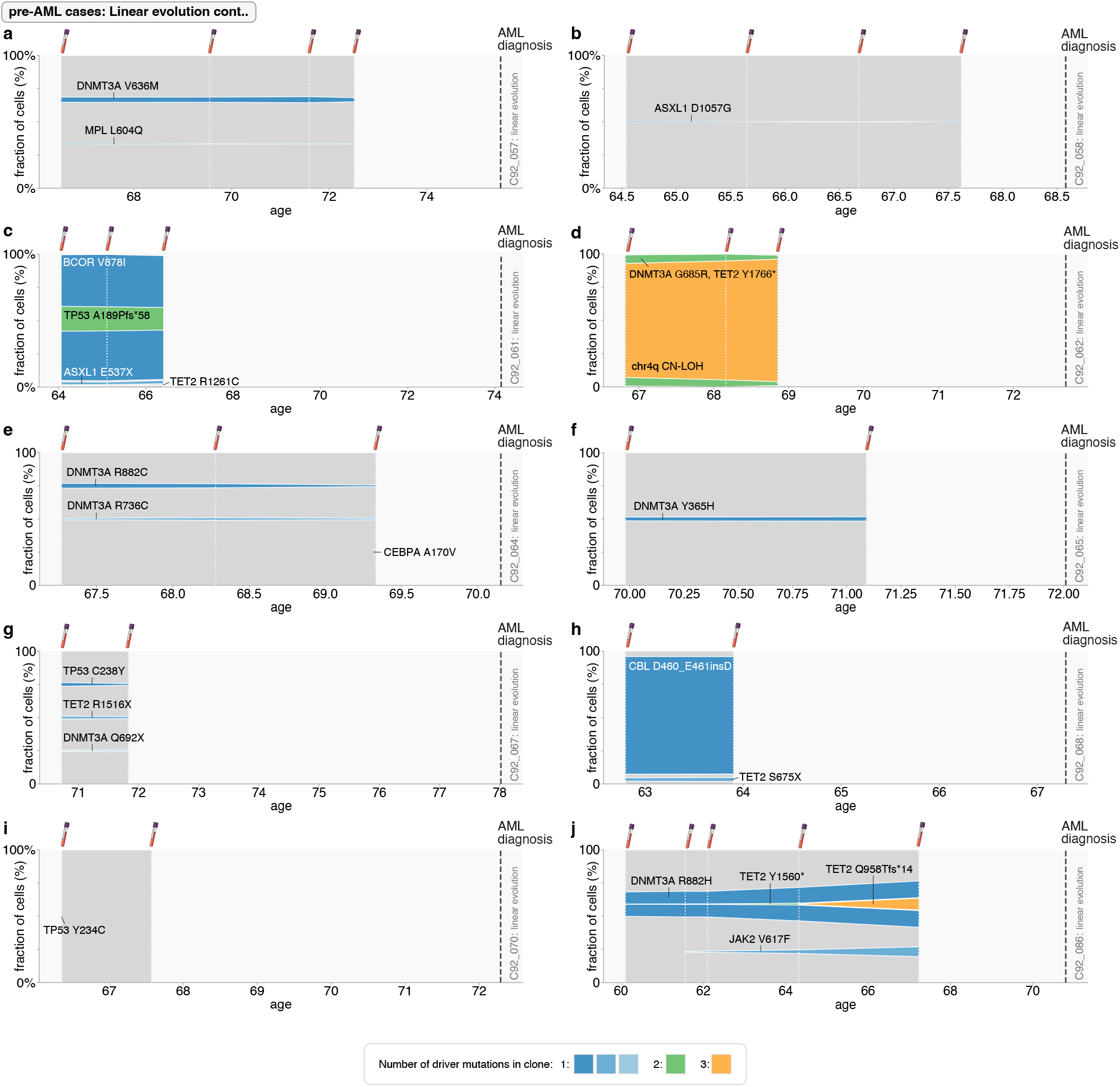
Reconstruction of clonal evolutionary histories: pre-AML cases with linear evolution part 2. Muller plots exhibiting clonal dynamics in pre-AML cases showing “linear evolution”. Colouring denotes the number of observed driver mutations in each clone (see legend). White vertical dashed lines indicate timing of UKCTOCS blood samples. Thick grey vertical dashed line indicates time of AML diagnosis.

**Supplementary Fig. 17.**
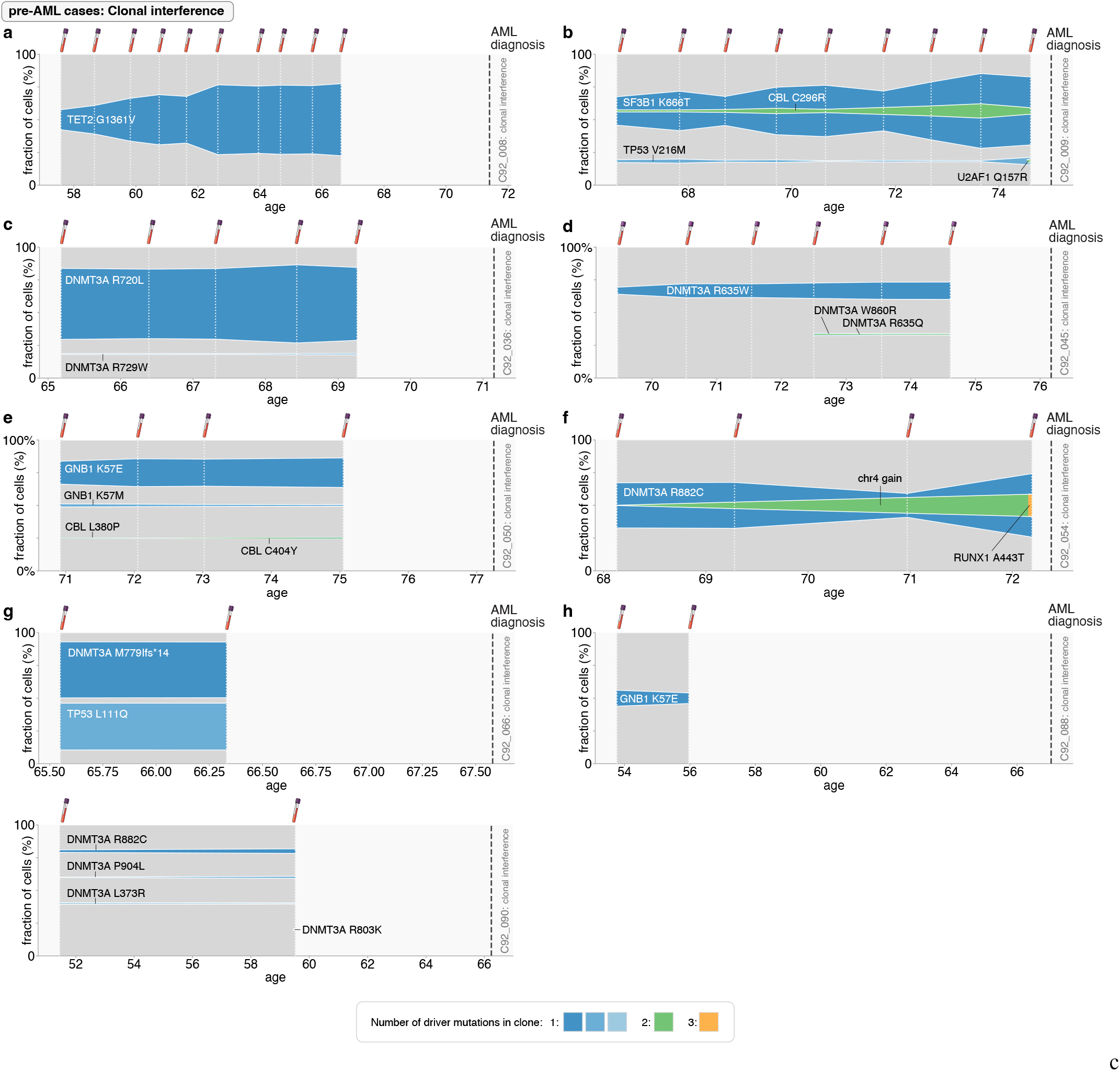
Reconstruction of clonal evolutionary histories: pre-AML cases with clonal interference. Muller plots exhibiting clonal dynamics in pre-AML cases showing “clonal interference”. Colouring denotes the number of observed driver mutations in each clone (see legend). White vertical dashed lines indicate timing of UKCTOCS blood samples. Thick grey vertical dashed line indicates time of AML diagnosis.

**Supplementary Fig. 18.**
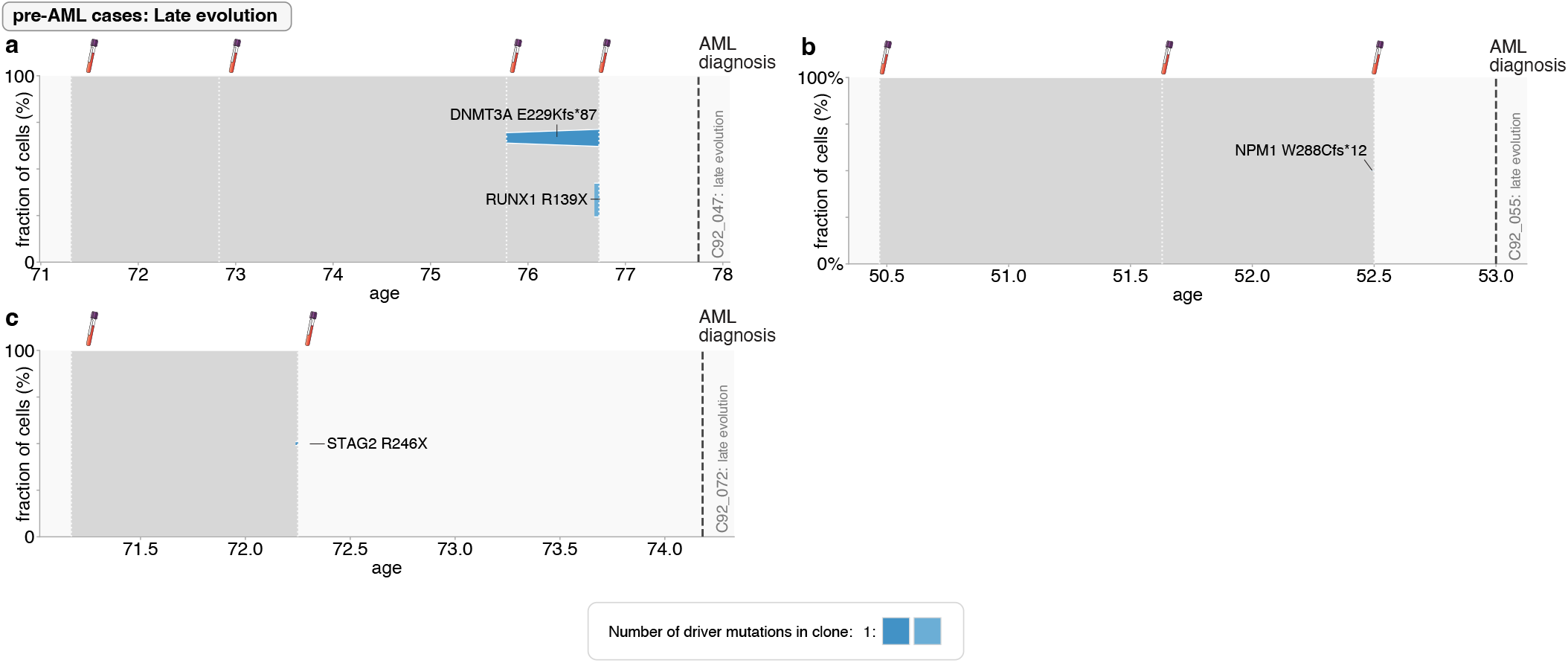
Reconstruction of clonal evolutionary histories: pre-AML cases with late evolution. Muller plots exhibiting clonal dynamics in pre-AML cases showing “late evolution”. Colouring denotes the number of observed driver mutations in each clone (see legend). White vertical dashed lines indicate timing of UKCTOCS blood samples. Thick grey vertical dashed line indicates time of AML diagnosis.

**Supplementary Fig. 19.**
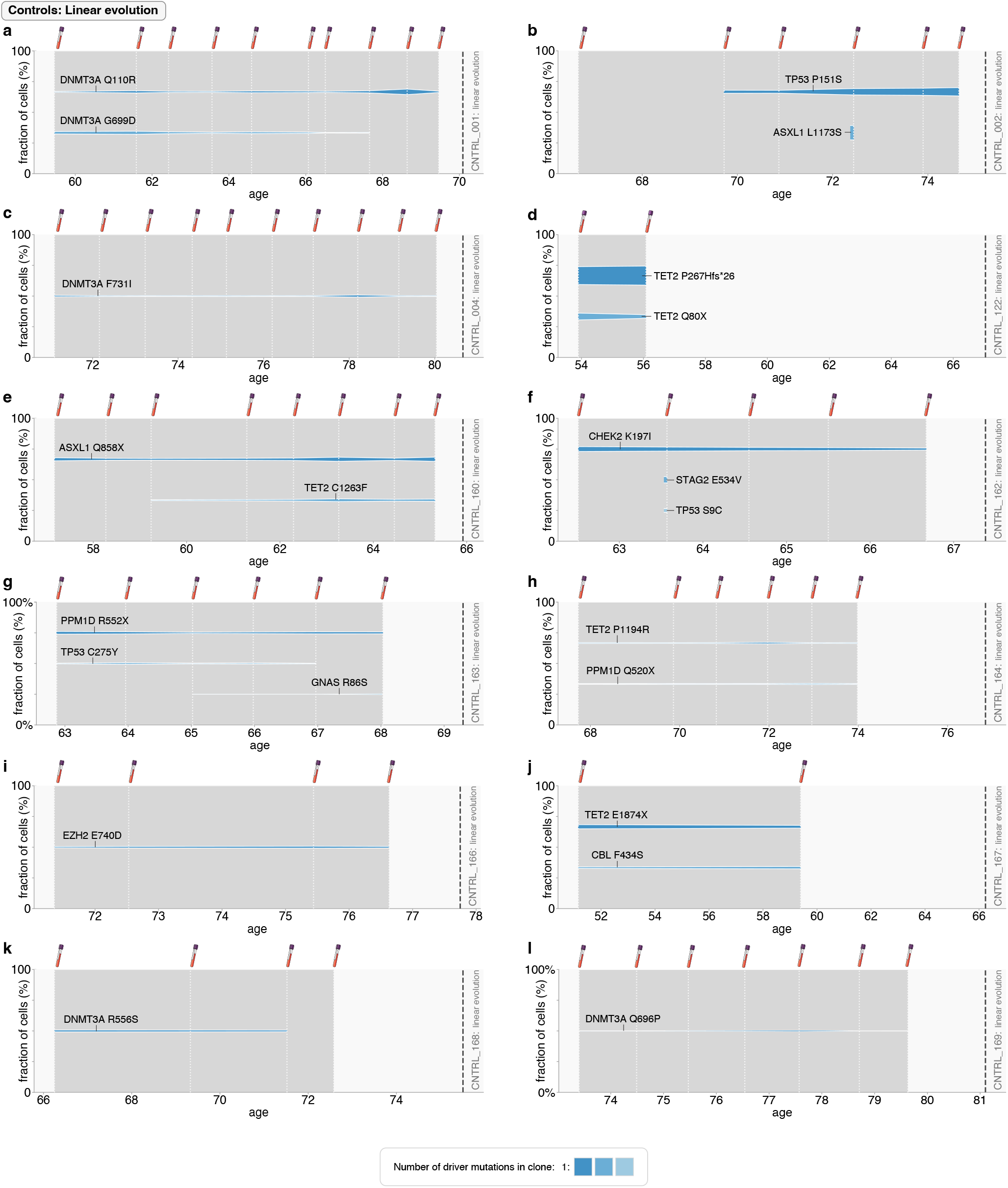
Reconstruction of clonal evolutionary histories: controls with linear evolution part 1. Muller plots exhibiting clonal dynamics in controls showing “linear evolution”. Colouring denotes the number of observed driver mutations in each clone (see legend). White vertical dashed lines indicate timing of UKCTOCS blood samples. Thick grey vertical dashed line indicates time of AML diagnosis in age- and timepoint-matched case.

**Supplementary Fig. 20.**
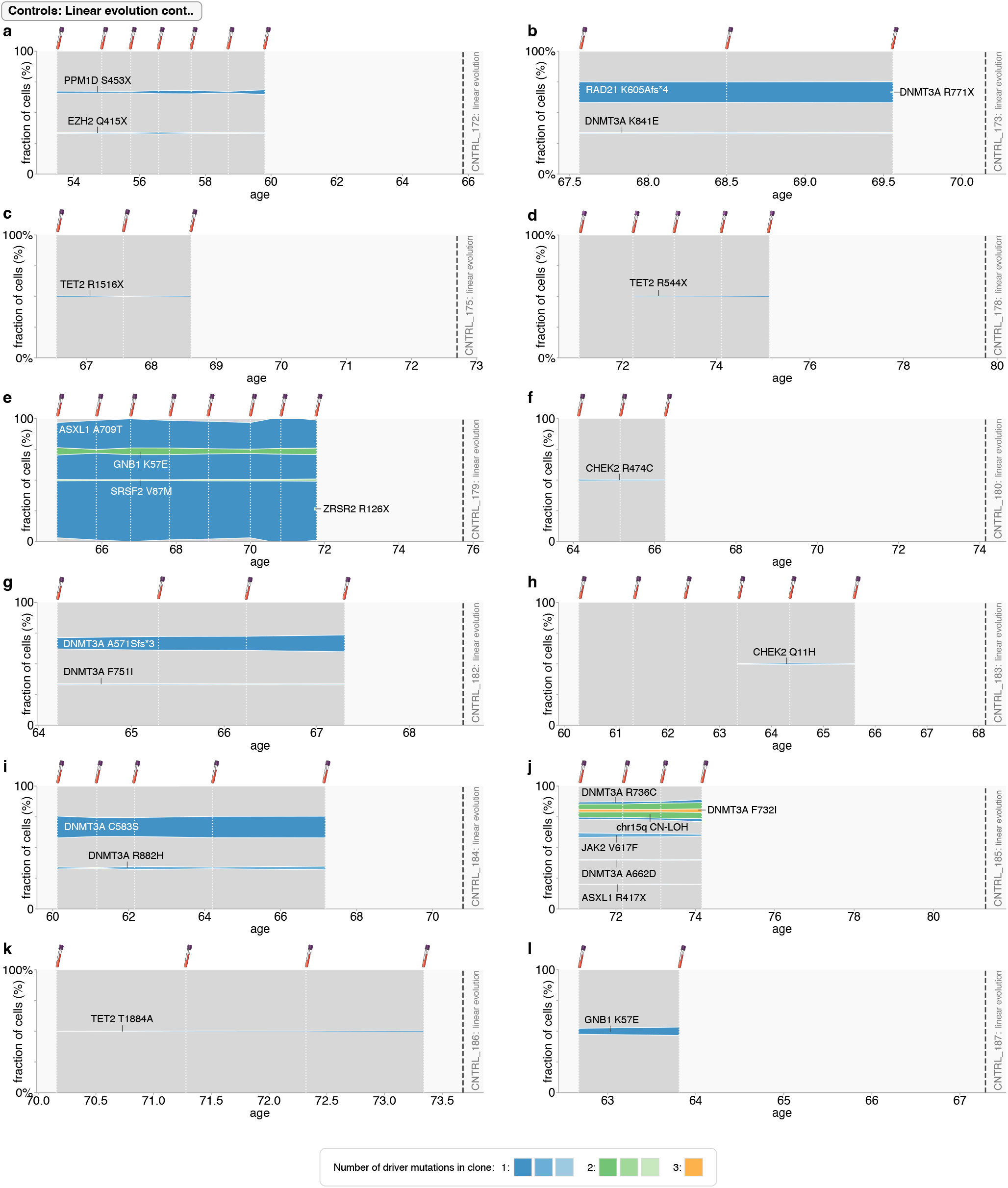
Reconstruction of clonal evolutionary histories: controls with linear evolution part 2. Muller plots exhibiting clonal dynamics in controls showing “linear evolution”. Colouring denotes the number of observed driver mutations in each clone (see legend). White vertical dashed lines indicate timing of UKCTOCS blood samples. Thick grey vertical dashed line indicates time of AML diagnosis in age- and timepoint-matched case.

**Supplementary Fig. 21.**
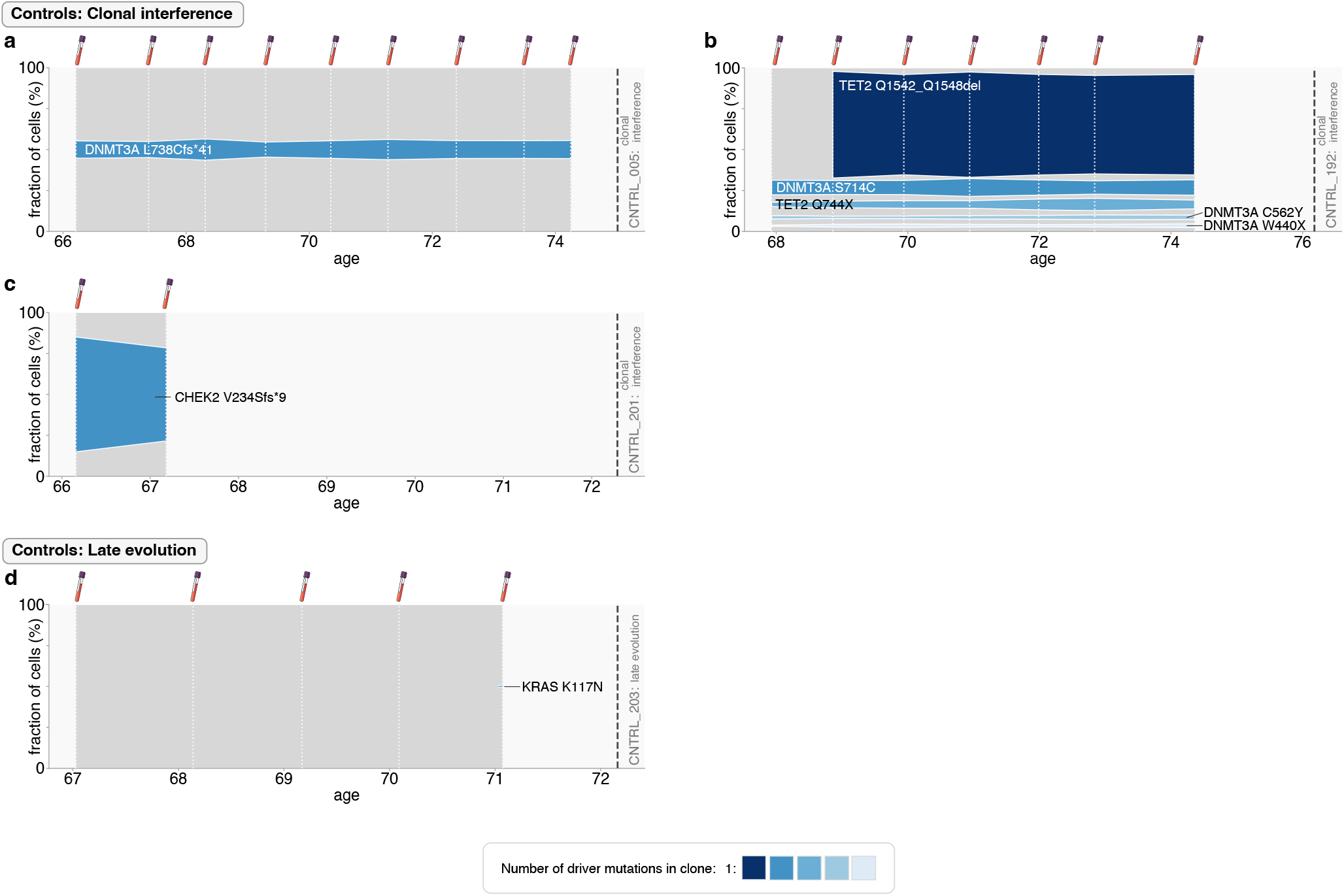
Reconstruction of clonal evolutionary histories: controls with clonal interference or late evolution. Muller plots exhibiting clonal dynamics in controls showing “clonal interference” or “late evolution”. Colouring denotes the number of observed driver mutations in each clone (see legend). White vertical dashed lines indicate timing of UKCTOCS blood samples. Thick grey vertical dashed line indicates time of AML diagnosis in age- and timepoint-matched case.

### 3. Inferring acquisition age and fitness effect of mutations

#### A. Initialisation of the maximum likelihood optimisation

Our maximum likelihood optimisation started with an ‘initial estimate’ for the establishment time and fitness of each clone. If a variant had VAF measurements from ≥2 timepoints that were <40% VAF, the initial fitness estimate was obtained from the gradient of a straight line drawn through the log(VAF) trajectory and the establishment time from where this straight line intercepted the y-axis at 1/*Nτ*s = 10^−5^. If there were VAF measurements from ≥4 timepoints, and the gradient changed by >40% in the second half of the trajectory, only the log(VAF) measurements from the earliest half of the trajectory were used for gradient estimation. If there were VAF measurements from only 1 timepoint, a ‘bounded VAF’ from the preceding and/ or subsequent timepoints (up to a maximum of two timepoints) was calculated as 1/(total depth) on the assumption that the variant was at a VAF lower than the ‘bounded VAF’ at that timepoint, otherwise it would have been detected. The ‘initial estimate’ for establishment time and fitness was then inferred using the measured VAF and bounded VAF(s). If the variant with a VAF measurement from only one timepoint was not a single-mutant clone, the ‘initial estimate’ inferred using the ‘bounded VAF’ was compared to the ‘initial estimate’ for the establishment time and fitness of the predecessor clone. If the ‘initial estimate’ for establishment time and fitness of the predecessor clone was higher, then this was used as the clone’s ‘initial estimate’ instead. The ‘initial estimate’ for establishment times were capped to a minimum of 0 years. If the ‘initial estimate’ for fitness was inferred to be <0, then the ‘initial estimate’ for establishment and fitness was based on the number of mutations in the clone as follows:

**Table S2.** ‘Initial estimate’ used in maximum likelihood optimiser for establishment time and fitness, based on number of mutations in clone. If the sample was from a control, ‘AML diagnosis age’ was that of its matched pre-AML case.

| Number of mutations in clone | Fitness (% per year) | Establishment time (years) |
| --- | --- | --- |
| 1 | 10% | AML diagnosis age / 5 |
| 2 | 30% | AML diagnosis age / 2 |
| 3 | 90% | AML diagnosis age - 10 |
| 4 | 180% | AML diagnosis age - 7 |
| 5 | 300% | AML diagnosis age - 5 |
| 6 | 400% | AML diagnosis age - 3 |

For ‘missing drivers’, the ‘initial estimate’ was based on the number of mutations in the clone. If the ‘missing driver’ was not a single-mutant clone, an ‘initial estimate’ for establishment time and fitness was also inferred using the final two timepoints of the ‘missing driver”s predecessor clone. If this estimated fitness was higher then the predecessor clone based ‘initial estimate’ for establishment time and fitness was used instead of that based on number of mutations. For some samples with very complex clonal architecture, a manual ‘initial estimate’ was used.

#### B. Maximum likelihood optimisation

Once the ‘initial estimate’ for the establishment time and fitness of each clone had been set, a 3-stage maximum likelihood process was employed, each involving progressively finer parameter adjustments with each optimisation stage. With each optimisation, after the log(likelihood) of the measured VAFs (given the predicted VAFs inferred from each variant’s estimated establishment time and fitness) had been calculated, the estimated establishment time and fitness were adjusted and the log(likelihood) recalculated. If the log(likelihood) using the new parameters was improved, then these parameters were accepted as the new estimates for the next optimisation attempt. Adjustments were made by randomly choosing a new fitness or establishment time from a normal distribution centred around the old establishment time or fitness. The standard deviation of the fitness distribution was that of the previous fitness estimate, multiplied by 0.1 for optimisation stage 1, 0.01 for optimisation stage 2 and 0.001 for optimisation stage 3. The standard deviation of the establishment time was 1 for optimisation stage 1, 0.2 for optimisation stage 2 and 0.04 for optimisation stage 3. These optimisation adjustments were repeated 5000 times in optimisation stage 1, 2000 times in optimisation stage 2 and 1000 times in optimisation stage 3. For some samples with very complex clonal architecture, the number of optimisation attempts in the optimisation stage 1 was manually increased to >5000 until there appeared to be no further incremental improvement in the log(likelihood) during that stage. For each sample, the 3-stage maximum likelihood process was repeated for 25 ‘seeds’ and the results of the seed that produced the lowest total log(likelihood) provided the inferred establishment times and fitnesses of all the variants in that sample.

Penalties were applied to the calculated log(likelihoods) in certain situations (i.e. making the estimated establishment time and fitness less likely to be accepted):

- If, from the predicted variant trajectory, a variant’s predicted VAF is greater than the ‘bounded’ non-detected VAF at a particular timepoint, penalise the log(likelihood) by multiplying the log(likelihood) for that variant timepoint by 3. This reduces the chance of fitting a trajectory that should have been detectable at timepoints at which it was not actually detectable.
- For variants that are detected at only one timepoint (and which are not single mutant clones), penalise the log(likelihood) (x3) for the variant trajectory if the inferred fitness effect is lower that it’s predecessor clone or if it’s inferred establishment time is earlier than it’s predecessor clone.
- Penalise the log(likelihood) (+10000) for a variant trajectory if the inferred establishment time is <0 years old, it is earlier than it’s predecessor clone (if it has one) or it is later than the AML diagnosis age.

**Supplementary Fig. 22.**
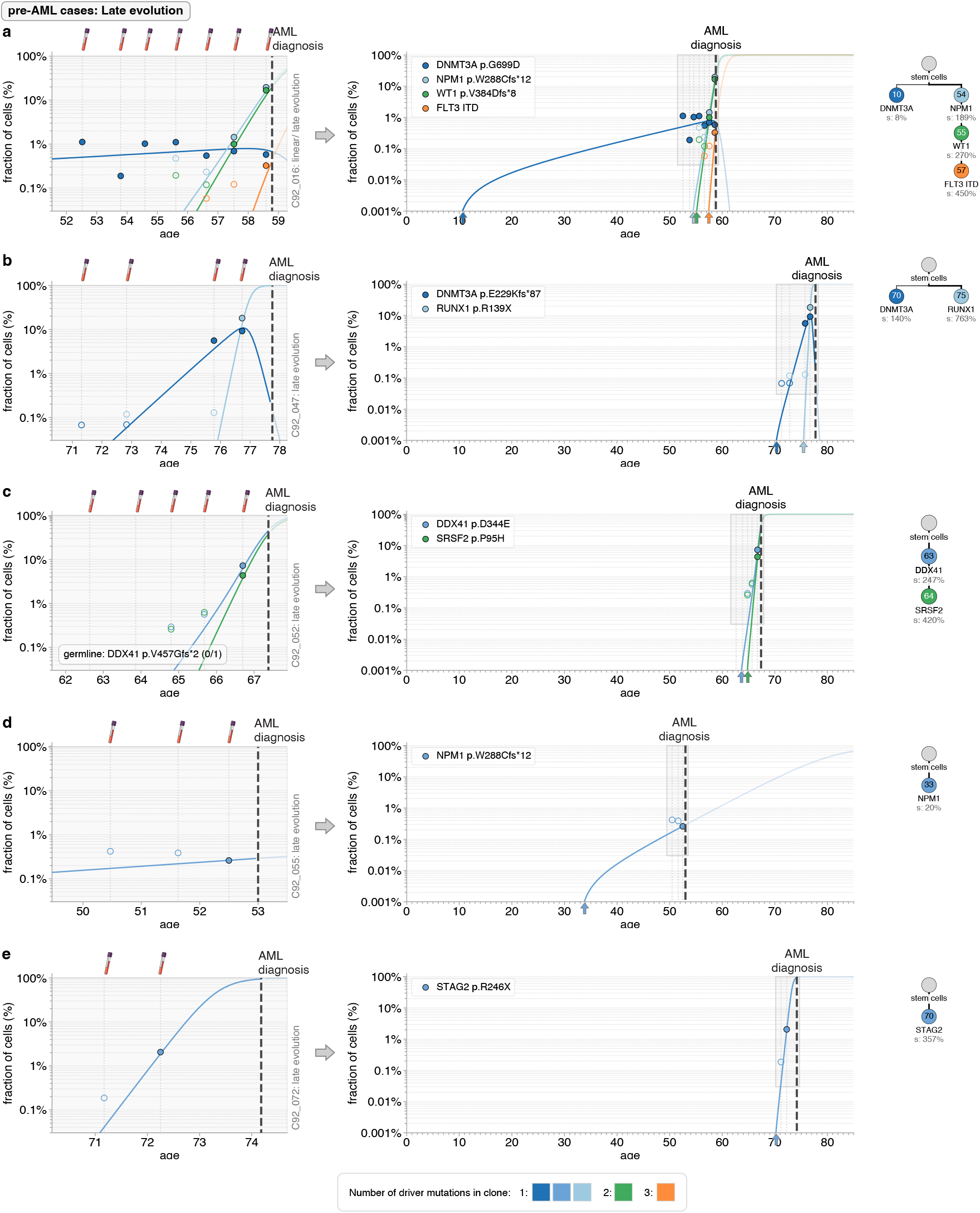
Quantitative dynamics of driver mutations in the decades before AML diagnosis: pre-AML cases with late evolution. **a-e**. Observed variant frequency trajectories (data points) compared with predicted variant frequency trajectories (coloured lines) due to the most likely fitness and occurrence time estimates of clones, across the period of UKCTOCS blood sampling (left-hand-side plots) and since the birth of the individual (right-hand-side plots). For some variants, hollow datapoints are shown to indicate the cell fraction limit of detection (1/DCS depth) at that timepoint (if the variant was not detected at that timepoint, but was detected at other timepoints). Trajectories of any inferred ‘missing drivers’ are indicated by dashed coloured lines. Grey vertical dashed lines indicate timing of blood samples. Thick black vertical dashed line indicates time of AML diagnosis. Phylogenetic trees (right-hand-side) show the inferred clonal composition of the population, occurrence times of each clone (number in circles, years) and their fitness effect (% per year). The number of driver mutations in the clone is indicated by its colour (see legend).

**Supplementary Fig. 23.**
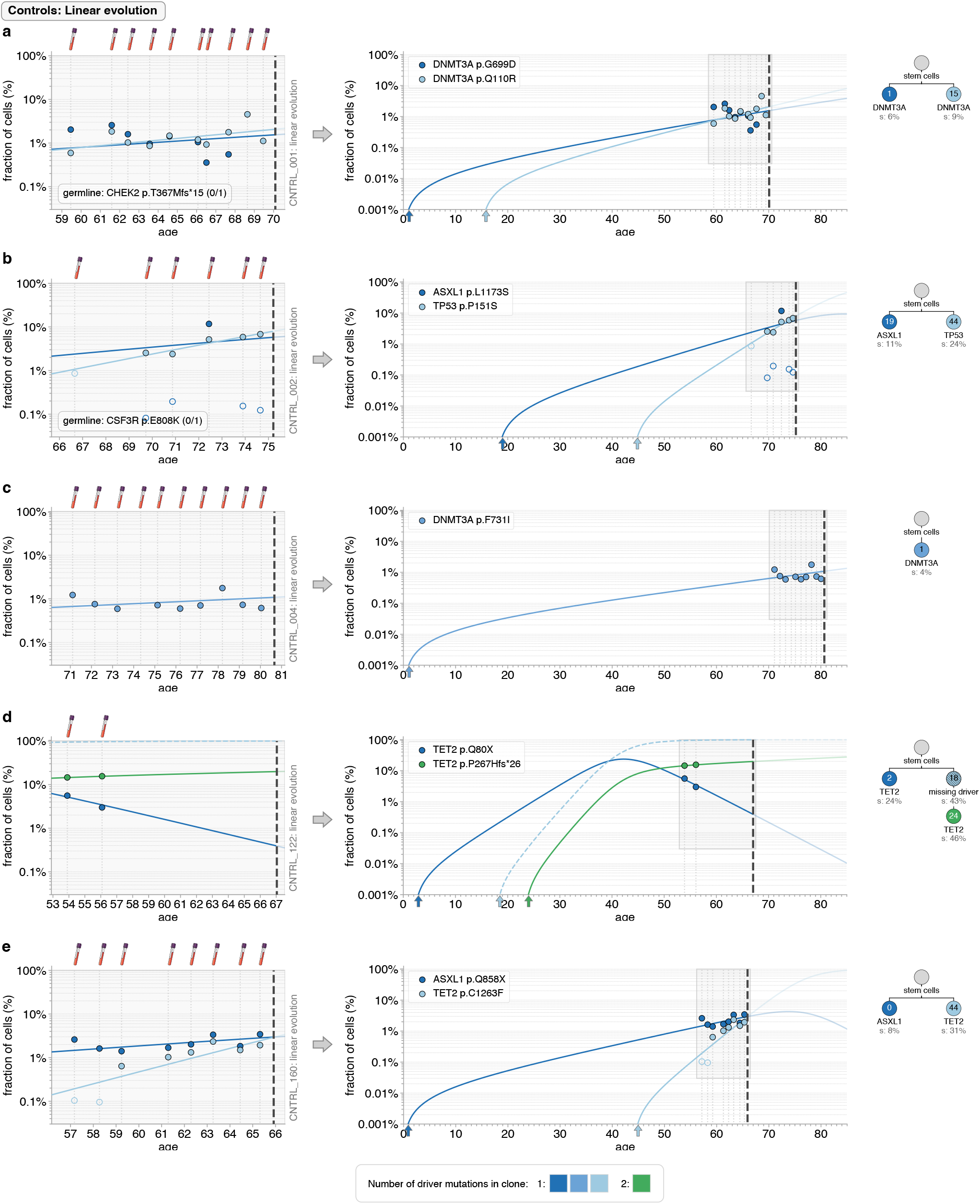
Quantitative dynamics of driver mutations in controls with linear evolution part 1. **a-e**. Observed variant frequency trajectories (data points) compared with predicted variant frequency trajectories (coloured lines) due to the most likely fitness and occurrence time estimates of clones, across the period of UKCTOCS blood sampling (left-hand-side plots) and since the birth of the individual (right-hand-side plots). For some variants, hollow datapoints are shown to indicate the cell fraction limit of detection (1/DCS depth) at that timepoint (if the variant was not detected at that timepoint, but was detected at other timepoints). Trajectories of any inferred ‘missing drivers’ are indicated by dashed coloured lines. Grey vertical dashed lines indicate timing of blood samples. Thick black vertical dashed line indicates time of AML diagnosis in the age- and timepoint-matched case. Phylogenetic trees (right-hand-side) show the inferred clonal composition of the population, occurrence times of each clone (number in circles, years) and their fitness effect (% per year). The number of driver mutations in the clone is indicated by its colour (see legend).

**Supplementary Fig. 24.**
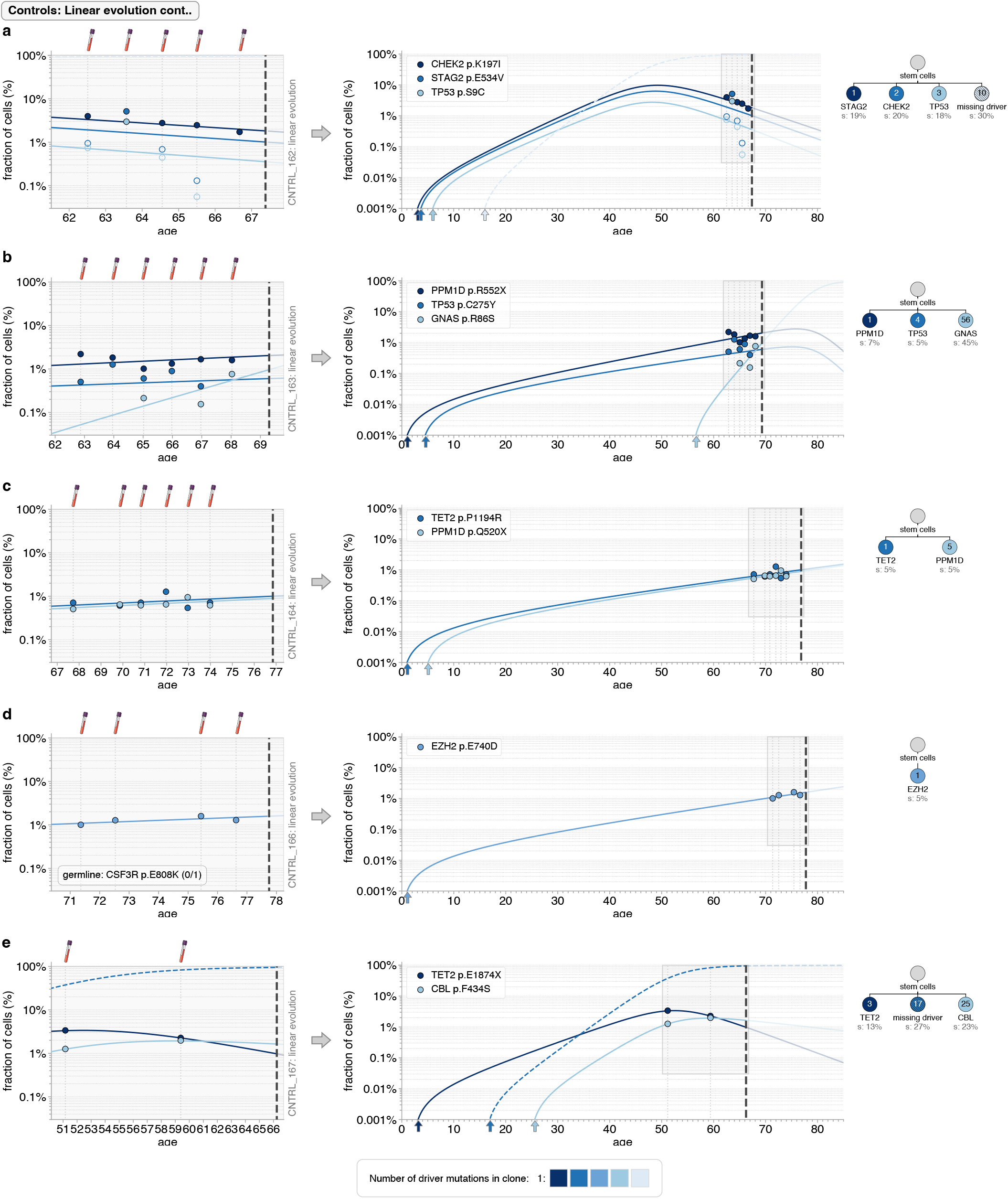
Quantitative dynamics of driver mutations in controls with linear evolution part 2. **a-e**. Observed variant frequency trajectories (data points) compared with predicted variant frequency trajectories (coloured lines) due to the most likely fitness and occurrence time estimates of clones, across the period of UKCTOCS blood sampling (left-hand-side plots) and since the birth of the individual (right-hand-side plots). For some variants, hollow datapoints are shown to indicate the cell fraction limit of detection (1/DCS depth) at that timepoint (if the variant was not detected at that timepoint, but was detected at other timepoints). Trajectories of any inferred ‘missing drivers’ are indicated by dashed coloured lines. Grey vertical dashed lines indicate timing of blood samples. Thick black vertical dashed line indicates time of AML diagnosis in the age- and timepoint-matched case. Phylogenetic trees (right-hand-side) show the inferred clonal composition of the population, occurrence times of each clone (number in circles, years) and their fitness effect (% per year). The number of driver mutations in the clone is indicated by its colour (see legend).

**Supplementary Fig. 25.**
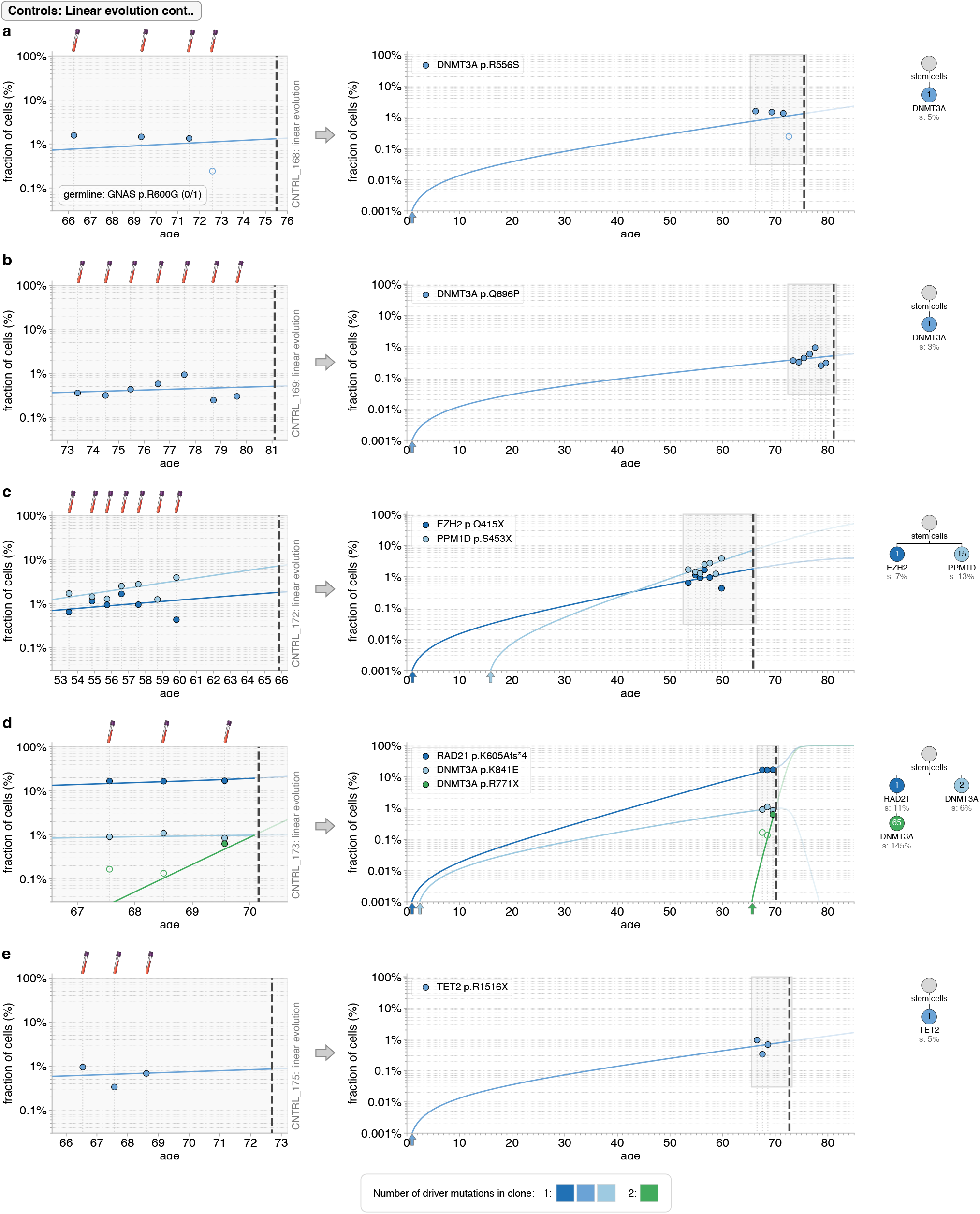
Quantitative dynamics of driver mutations in controls with linear evolution part 3. **a-e**. Observed variant frequency trajectories (data points) compared with predicted variant frequency trajectories (coloured lines) due to the most likely fitness and occurrence time estimates of clones, across the period of UKCTOCS blood sampling (left-hand-side plots) and since the birth of the individual (right-hand-side plots). For some variants, hollow datapoints are shown to indicate the cell fraction limit of detection (1/DCS depth) at that timepoint (if the variant was not detected at that timepoint, but was detected at other timepoints). Trajectories of any inferred ‘missing drivers’ are indicated by dashed coloured lines. Grey vertical dashed lines indicate timing of blood samples. Thick black vertical dashed line indicates time of AML diagnosis in the age- and timepoint-matched case. Phylogenetic trees (right-hand-side) show the inferred clonal composition of the population, occurrence times of each clone (number in circles, years) and their fitness effect (% per year). The number of driver mutations in the clone is indicated by its colour (see legend).

**Supplementary Fig. 26.**
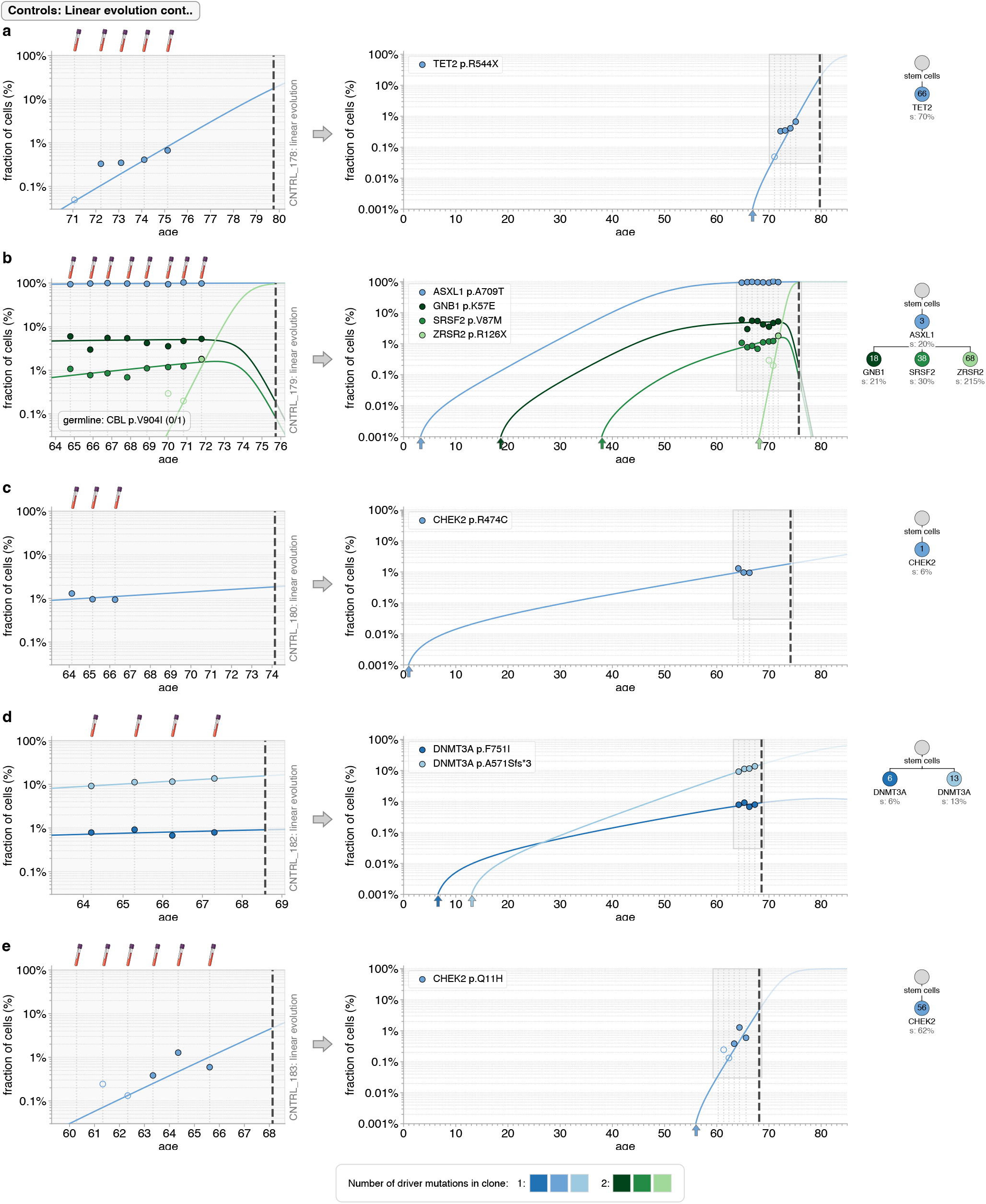
Quantitative dynamics of driver mutations in controls with linear evolution part 4. **a-e**. Observed variant frequency trajectories (data points) compared with predicted variant frequency trajectories (coloured lines) due to the most likely fitness and occurrence time estimates of clones, across the period of UKCTOCS blood sampling (left-hand-side plots) and since the birth of the individual (right-hand-side plots). For some variants, hollow datapoints are shown to indicate the cell fraction limit of detection (1/DCS depth) at that timepoint (if the variant was not detected at that timepoint, but was detected at other timepoints). Trajectories of any inferred ‘missing drivers’ are indicated by dashed coloured lines. Grey vertical dashed lines indicate timing of blood samples. Thick black vertical dashed line indicates time of AML diagnosis in the age- and timepoint-matched case. Phylogenetic trees (right-hand-side) show the inferred clonal composition of the population, occurrence times of each clone (number in circles, years) and their fitness effect (% per year). The number of driver mutations in the clone is indicated by its colour (see legend).

**Supplementary Fig. 27.**
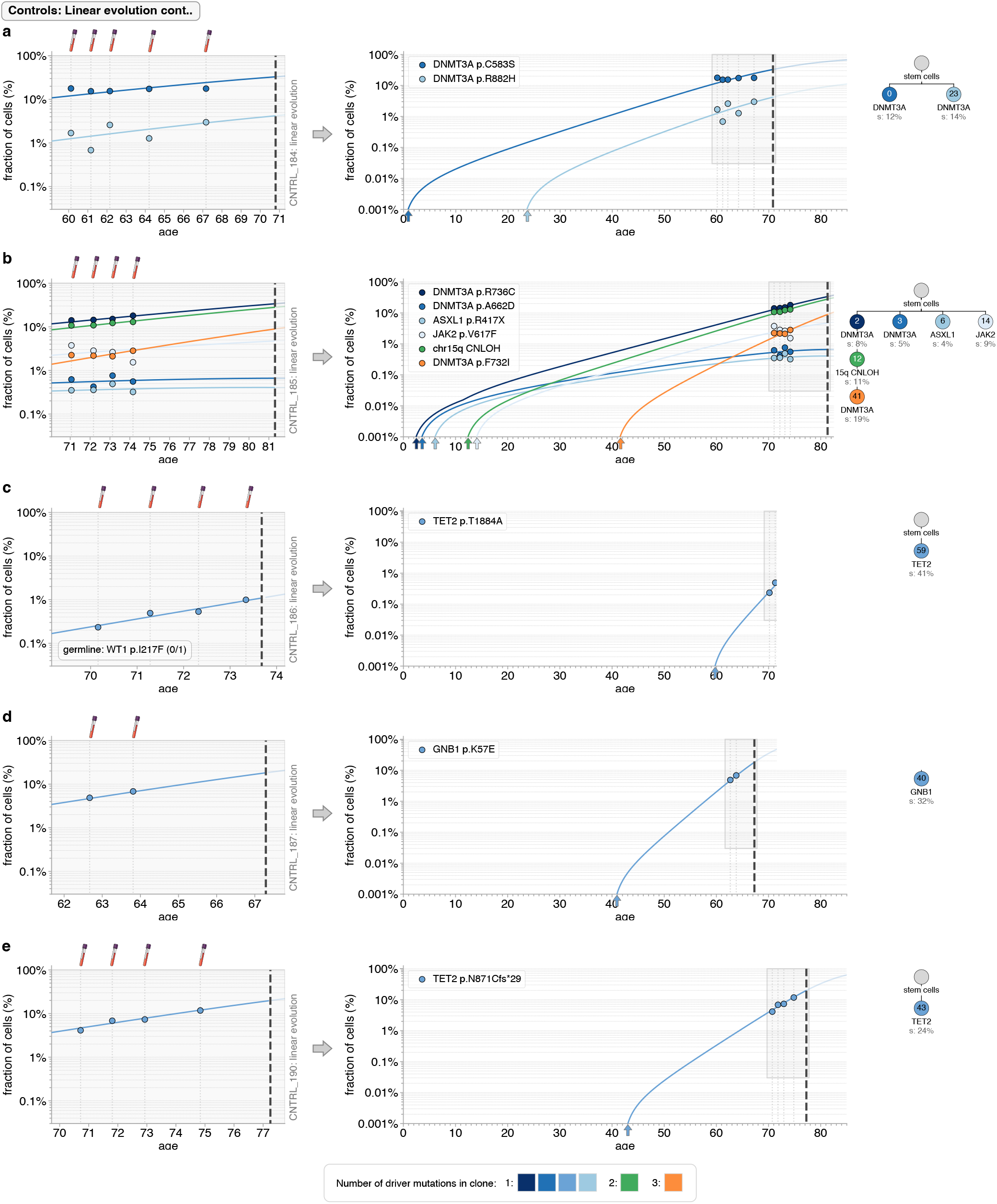
Quantitative dynamics of driver mutations in controls with linear evolution part 5. **a-e**. Observed variant frequency trajectories (data points) compared with predicted variant frequency trajectories (coloured lines) due to the most likely fitness and occurrence time estimates of clones, across the period of UKCTOCS blood sampling (left-hand-side plots) and since the birth of the individual (right-hand-side plots). For some variants, hollow datapoints are shown to indicate the cell fraction limit of detection (1/DCS depth) at that timepoint (if the variant was not detected at that timepoint, but was detected at other timepoints). Trajectories of any inferred ‘missing drivers’ are indicated by dashed coloured lines. Grey vertical dashed lines indicate timing of blood samples. Thick black vertical dashed line indicates time of AML diagnosis in the age- and timepoint-matched case. Phylogenetic trees (right-hand-side) show the inferred clonal composition of the population, occurrence times of each clone (number in circles, years) and their fitness effect (% per year). The number of driver mutations in the clone is indicated by its colour (see legend).

**Supplementary Fig. 28.**
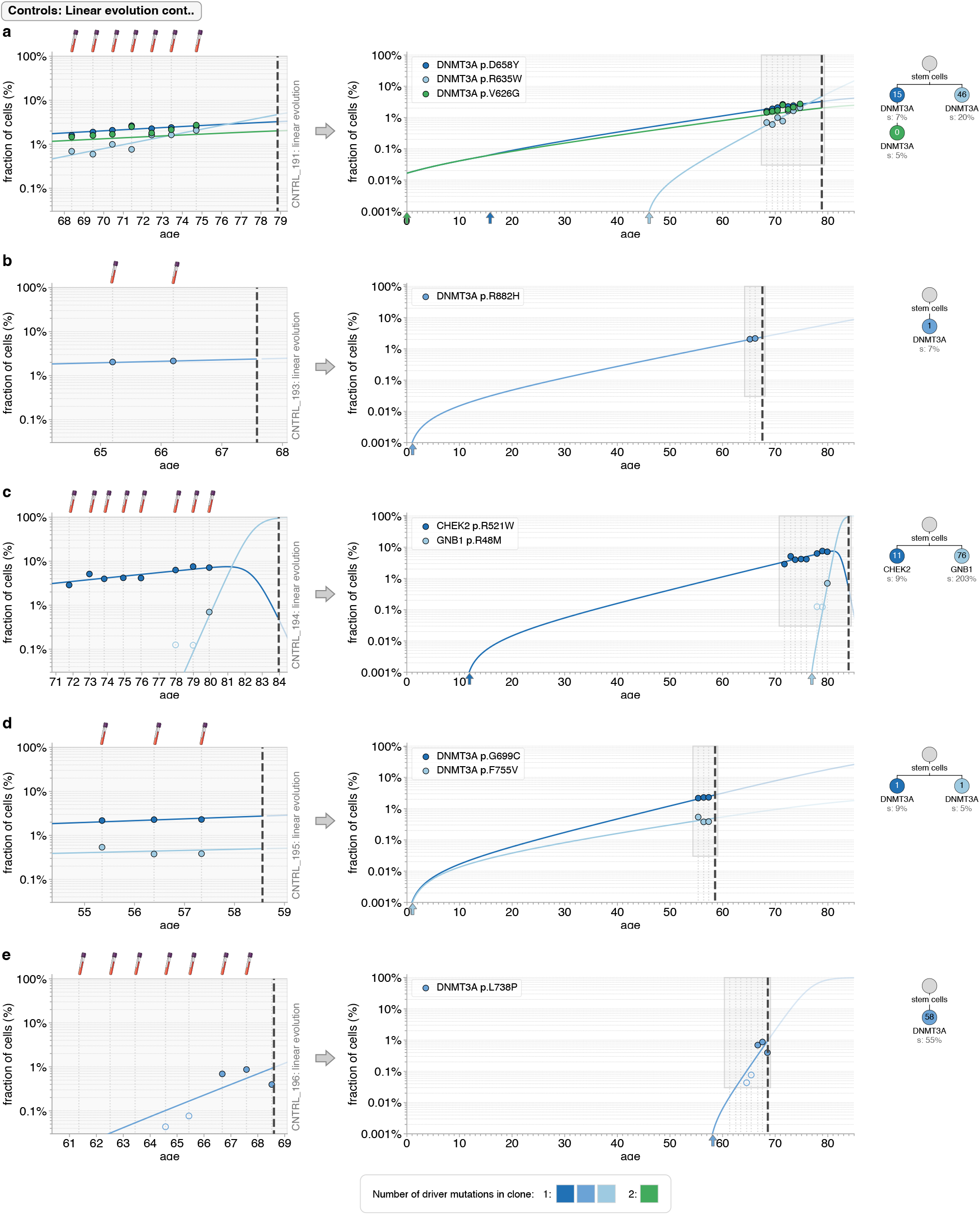
Quantitative dynamics of driver mutations in controls with linear evolution part 6. **a-e**. Observed variant frequency trajectories (data points) compared with predicted variant frequency trajectories (coloured lines) due to the most likely fitness and occurrence time estimates of clones, across the period of UKCTOCS blood sampling (left-hand-side plots) and since the birth of the individual (right-hand-side plots). For some variants, hollow datapoints are shown to indicate the cell fraction limit of detection (1/DCS depth) at that timepoint (if the variant was not detected at that timepoint, but was detected at other timepoints). Trajectories of any inferred ‘missing drivers’ are indicated by dashed coloured lines. Grey vertical dashed lines indicate timing of blood samples. Thick black vertical dashed line indicates time of AML diagnosis in the age- and timepoint-matched case. Phylogenetic trees (right-hand-side) show the inferred clonal composition of the population, occurrence times of each clone (number in circles, years) and their fitness effect (% per year). The number of driver mutations in the clone is indicated by its colour (see legend).

**Supplementary Fig. 29.**
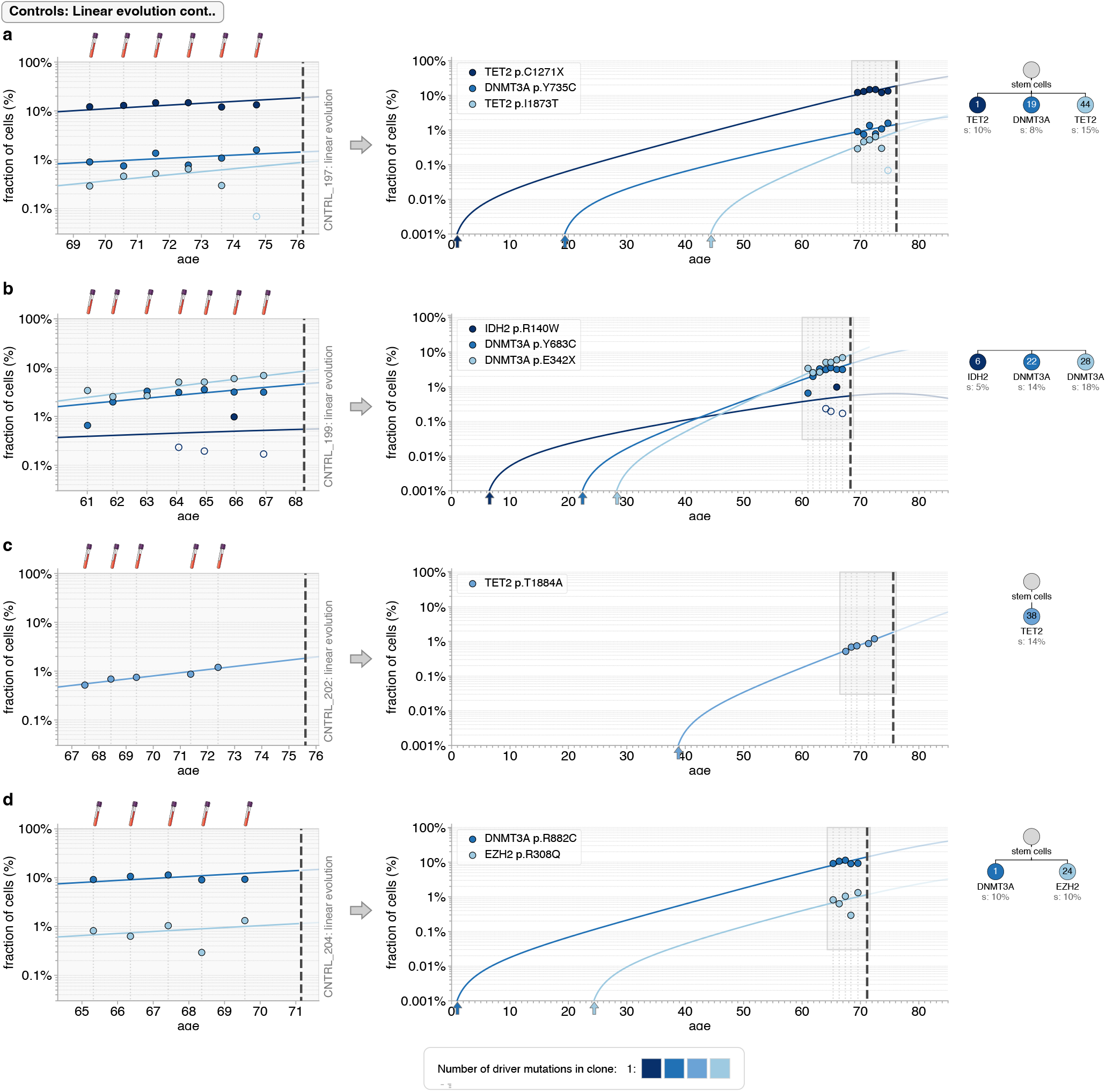
Quantitative dynamics of driver mutations in controls with linear evolution part 7. **a-e**. Observed variant frequency trajectories (data points) compared with predicted variant frequency trajectories (coloured lines) due to the most likely fitness and occurrence time estimates of clones, across the period of UKCTOCS blood sampling (left-hand-side plots) and since the birth of the individual (right-hand-side plots). For some variants, hollow datapoints are shown to indicate the cell fraction limit of detection (1/DCS depth) at that timepoint (if the variant was not detected at that timepoint, but was detected at other timepoints). Trajectories of any inferred ‘missing drivers’ are indicated by dashed coloured lines. Grey vertical dashed lines indicate timing of blood samples. Thick black vertical dashed line indicates time of AML diagnosis in the age- and timepoint-matched case. Phylogenetic trees (right-hand-side) show the inferred clonal composition of the population, occurrence times of each clone (number in circles, years) and their fitness effect (% per year). The number of driver mutations in the clone is indicated by its colour (see legend).

**Supplementary Fig. 30.**
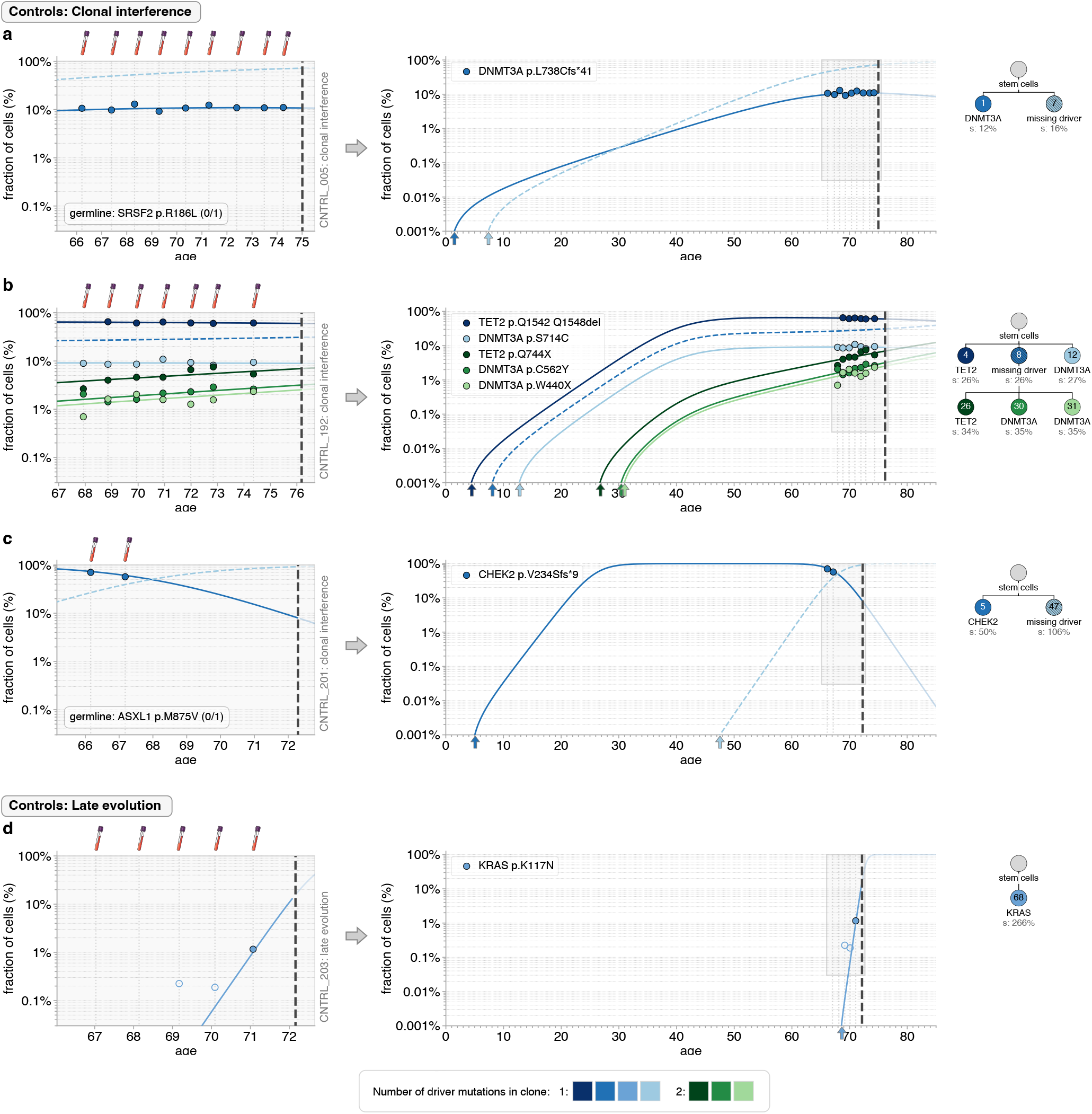
Quantitative dynamics of driver mutations in controls with clonal interference or late evolution. **a-d**. Observed variant frequency trajectories (data points) compared with predicted variant frequency trajectories (coloured lines) due to the most likely fitness and occurrence time estimates of clones, across the period of UKCTOCS blood sampling (left-hand-side plots) and since the birth of the individual (right-hand-side plots). For some variants, hollow datapoints are shown to indicate the cell fraction limit of detection (1/DCS depth) at that timepoint (if the variant was not detected at that timepoint, but was detected at other timepoints). Trajectories of any inferred ‘missing drivers’ are indicated by dashed coloured lines. Grey vertical dashed lines indicate timing of blood samples. Thick black vertical dashed line indicates time of AML diagnosis in the age- and timepoint-matched case. Phylogenetic trees (right-hand-side) show the inferred clonal composition of the population, occurrence times of each clone (number in circles, years) and their fitness effect (% per year). The number of driver mutations in the clone is indicated by its colour (see legend).

